# Latent generative search unlocks de novo design of untapped biomolecular interactions at scale

**DOI:** 10.64898/2026.09.12.751118

**Authors:** Kieran Didi, Danny Reidenbach, Matthew Penner, Supriya Ravichandran, Marshall Case, Mike Nichols, Erik Swanson, Alex Reis, Maggie Prescott, Yue Qian, Dongming Qian, Jingjing Yang, Weiji Li, Le Li, Daichi Shonai, Sean Gay, Bhoomika Basu Mallik, Ho Yeung Chim, Liurong Chen, Miguel Atienza Juanatey, Hubert Klein, Dominic Rieger, Phillip Schlegel, Anna U. Macintyre, Maxim Secor, Daniele Granata, Sooyoung Cha, Zhonglin Cao, Guoqing Zhou, Tomas Geffner, Xi Chen, Micha Livne, Zuobai Zhang, Tianjing Zhang, Kyle Gion, Michael M. Bronstein, Martin Steinegger, Kristine Deibler, Scott Soderling, Clara T. Schoeder, Alena Khmelinskaia, Florian Hollfelder, Christian Dallago, Emine Kucukbenli, Arash Vahdat, Pierce Ogden, Karsten Kreis

## Abstract

De novo protein design has advanced rapidly, yet designing binders to polar, solvent-exposed epitopes and small, flexible ligands remains challenging. Such hydrated surfaces and flexible molecules, including carbohydrates, provide few of the hydrophobic contacts favoured by current methods and have largely resisted de novo binders. To address this challenge, here we introduce latent generative search for binder design, a novel framework that uses reward-guided search at inference time to steer the Proteína-Complexa generative model. The model codesigns sequence and structure — generating them together in a continuous latent space — and thereby removes the inverse-folding step on which current methods rely. In a screen of more than one million designs by multiplexed phage display, latent generative search produced more validated binders than every other method tested, its codesigned sequences surpassing post hoc redesign. It delivered high-affinity binders across therapeutic receptors, a viral attachment protein and intracellular signalling targets. Our approach also accessed previously untapped biology, generating the first de novo proteins that bind a free carbohydrate, including one that discriminates between blood-group antigens — a polar, flexible target class beyond the reach of current design methods.

## 1 Introduction

Across artificial intelligence, the most difficult problems repeatedly led to a common recipe: a generative model that learns the structure of a solution space, paired with a search procedure that explores that space at inference time. Neither component alone offers the same capabilities: the learned prior supplies plausible candidates, while search refines them toward a goal. Search-based methods first achieved landmark successes in constrained environments like board games [1]. More recently, they proved transformative in conjunction with generative models that make challenging, unstructured problems amenable to search, including image generation [2, 3] and language modelling, for instance through test-time reasoning [4, 5].

Designing proteins that bind chosen targets with high affinity and selectivity is a problem of similar difficulty, with broad implications for therapeutics, diagnostics and synthetic biology [6]. Deep learning made it increasingly attainable [7, 8]: protein structure-prediction models provide accurate target structures [9, 10] and filtering metrics [11, 12], while generative methods, such as RFdiffusion [13], or hallucination approaches, such as BindCraft [14], produce candidate backbones. However, the pairing of a generative prior with principled search that has seen success elsewhere has not been realised here. Generative models typically produce a protein structure in a single pass, with no optimisation of the sample, and quality often falls short without extensive downstream filtering [15]. Hallucination methods instead search over sequences without a learned generative prior to guide them, which can make the search brittle and prone to non-physical solutions [16]. Both paradigms also depend on a separate inverse-folding step to assign sequences to backbones, which can degrade electrostatic and other functionally critical interactions [14, 17]. These shortcomings are most acute at polar, solvent-exposed interfaces and against carbohydrates and flexible peptides, where sequence and backbone geometry co-optimisation would enable precise placement of hydrogen bonds — and such targets therefore remain largely out of reach.

Here we bring generative search to protein design, combining a learned protein prior with goal-directed search at inference time beyond filtering. The underlying generative model, Proteína-Complexa, and its inference-time optimization algorithms were introduced previously and shown to be effective for binder design in silico, without wet-lab testing [18]; the model builds on the La-Proteína latent protein representation [19]. We generalise that work into a latent generative search framework for real-world design, defined by application-specific, multi-objective reward functions tailored to each target class, and validate it experimentally at scale across protein–protein, peptide–protein, and protein–carbohydrate interactions (Fig. 1).

**Fig. 1.**
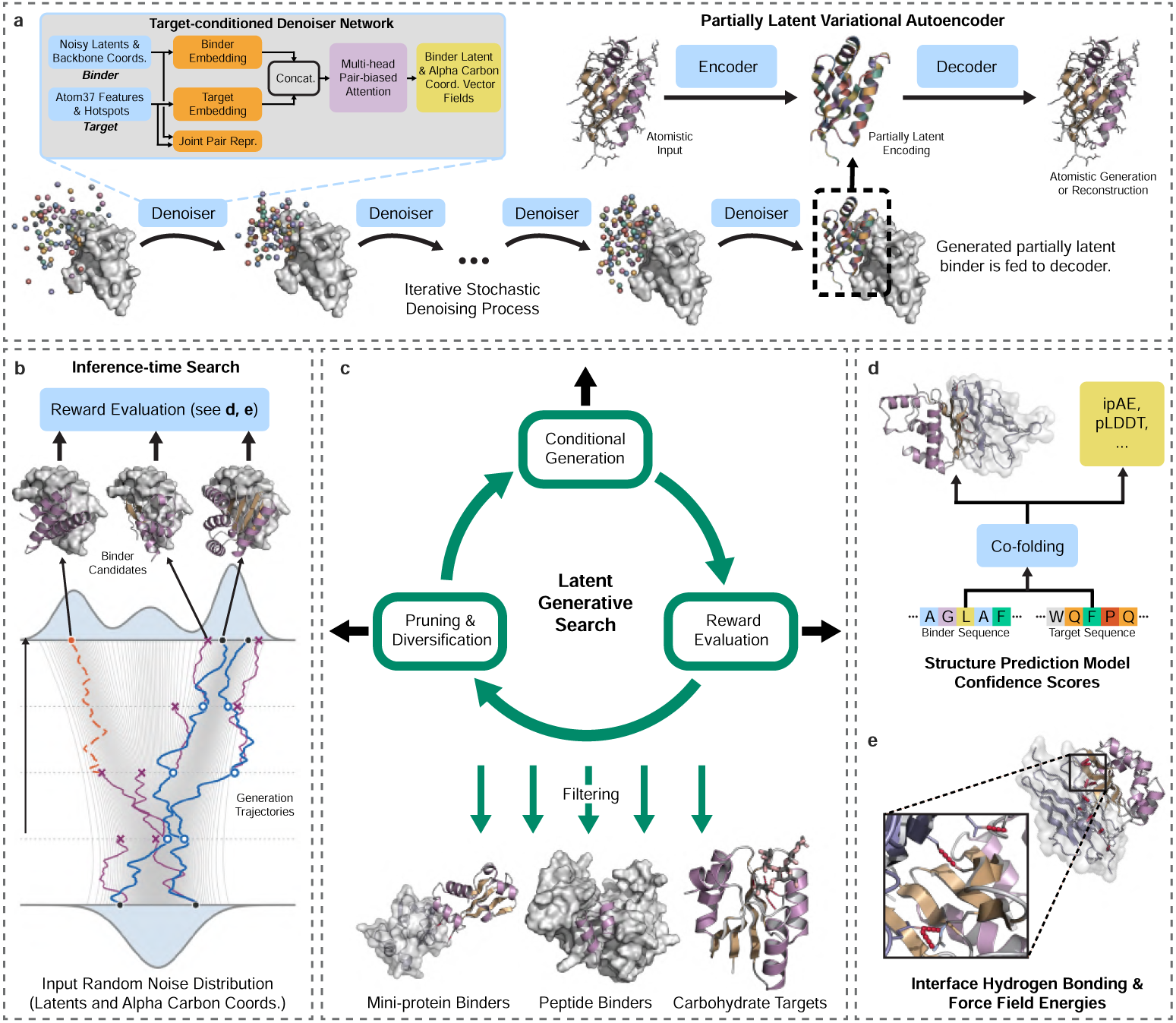
Latent generative search for de novo protein binder design. **a**, The conditional codesign generator, Proteína-Complexa, generates binder structure and sequence together using a partially latent representation [19]. A target-conditioned denoiser network iteratively transforms random noise into backbone coordinates and latent variables, after which a decoder maps the codesigned latent binder representation to a fully atomistic output. An auxiliary encoder is used during training of the variational autoencoder. **b**, Generative sampling trajectories are guided towards high reward regions of the solution space according to task-relevant scoring models in an inference-time search process. Low-scoring trajectories are terminated, whereas promising trajectories are expanded and diversified, for example through beam search. **c**, The complete latent generative search protocol alternates between generative denoising rollouts, reward-based trajectory evaluation, and the subsequent selection and expansion of promising trajectories. The resulting candidates may undergo a final filtering stage using additional scoring functions. The same framework supports diverse design tasks — including monomers, mini-protein binders, peptide binders, and binders to carbohydrate targets — by adapting the conditioning input, reward definition, and scoring models. **d**, Confidence metrics from structure prediction models, including ipAE and pLDDT, can serve as rewards during search or as criteria for final filtering. **e**, Interface hydrogen bonding scores and molecular force fields provide complementary physics-based scoring functions.

Here and throughout, we use *codesign* to mean generating an amino-acid sequence and its all-atom structure together within one generative process, without assigning a new sequence afterward by inverse folding. A *codesigned sequence* is therefore the model’s direct sequence output for its simultaneously generated structure. The framework combines this codesign process with reward-guided search to steer generation toward defined objectives. We evaluate it through biophysical characterisation of designed monomers (Section 3) and initial binder tests against PDGFR, PD-L1, and the Nipah virus G-glycoprotein (Section 4); a benchmark of more than one million designs against 127 diverse and challenging targets (Section 5); nanomolar binders to the muscle-wasting receptor ActRIIA that block signalling in cells (Section 6); peptide and mini-protein kinase binders (Section 7); and, to our knowledge, the first experimentally validated de novo binders to a carbohydrate target, a blood-group antigen (Section 8). Together, these results establish latent generative search as a state-of-the-art framework for protein binder design.

## 2 A latent generative search framework

A central difficulty in atomistic protein generation is that the number of atoms varies with amino-acid identity, complicating the joint modelling of sequence and structure. La-Proteína addresses this with a partially latent representation: backbone C*_α_* coordinates are modelled explicitly, while each residue’s amino-acid identity and remaining atoms are compressed into a fixed-length continuous latent variable by a variational autoencoder [19]. A flow-matching model then learns the joint distribution over backbone coordinates and latents, so that every codesigned model sample represents a complete protein — backbone, side chains and sequence — without requiring a separate sequence-design step [20].

Proteína-Complexa extends this representation to binder–target complexes by conditioning the flow model on a target structure and optional interface hotspot residues, while the autoencoder continues to encode and decode monomeric binders [18] (Fig. 1a). The model is trained in stages: the autoencoder and partially latent flow model are pre-trained on monomer structures, followed by training of the latent prior on Teddymer — a large synthetic dataset of binder–target pairs derived from domain–domain interactions in AlphaFold Database structures [21, 22] — and experimental multimers from the Protein Data Bank [23]. Because the generator is efficient, generation can be combined with reward-guided search at inference time: rather than accepting or rejecting individual samples, the model repeatedly generates and evaluates multiple stochastic denoising trajectories, selecting and expanding promising trajectories to steer generation towards high-quality binders (Fig. 1b,c; details in Methods Section 10). The same process can partially noise an existing binder and code-sign a related backbone and sequence, enabling scaffold-based re-engineering rather than fixed-backbone sequence redesign.

The rewards used in this study encode molecular requirements familiar to biochemists and can be arbitrarily extended. First, hydrogen-bond terms favour correctly placed donor–acceptor interactions and satisfaction of buried polar groups (Fig. 1e) [24, 25]. Second, clash avoidance, shape complementarity, pocket burial and interface packing assess steric and docking compatibility. Third, structure-prediction confidence and Rosetta/Tmol energy terms assess whether the designed protein and complex are likely to fold and remain stable [26, 27]. Fourth, protein-language-model likelihoods favour sequences that are plausible in their local and global context. Fifth, hotspot engagement requires the designed binder to contact specified target residues expected to contribute to binding. Different campaigns combine these molecular terms in different proportions, allowing the same search framework to address protein interfaces, peptides and free carbohydrates.

The latent generative search framework therefore combines complementary advantages: its learned generative prior anchors search in the distribution of plausible proteins, enabling more efficient and robust exploration than prior hallucination methods, while inference-time steering provides greater control over generation than unguided generative approaches. Joint sequence–structure codesign avoids post hoc inverse-folding redesign, which can disrupt functionally important side-chain interactions, while the flexible reward framework adapts the same search process to a broad range of design tasks.

## 3 Designed monomeric proteins

Reliable binder design requires generative models that produce both designable backbones and realistic sequences encoding stable, functional proteins, an end-to-end capability that remains an open experimental challenge. Leading approaches such as RFdiffusion and RFdiffusion2 generate protein geometry and rely on ProteinMPNN or LigandMPNN for the final sequences [13, 28]. Even all-atom, sequence-aware methods often retain post hoc redesign: RFdiffusion3 generates alanine-biased sequences whose sequence–structure consistency improves after ProteinMPNN redesign, while BoltzGen includes a dedicated inverse-folding module to improve predicted refolding and solubility [29, 30]. Because Proteína-Complexa instead codesigns sequence and structure in a single pass, we posed a controlled experimental test: do its codesigned sequences express, fold and remain soluble as well as ProteinMPNN-designed sequences? This comparison tests whether generative codesign can replace inverse folding, rather than only whether the model produces designable backbones. To our knowledge, no generative codesign method had previously demonstrated experimentally that its codesigned sequences perform at parity with a dedicated inverse-folding model.

We characterised 48 designed monomers spanning 100–800 amino acids (Fig. 2a). For 24 designs, sequence and backbone were codesigned; for the other 24, the generated backbones were redesigned using ProteinMPNN. For four backbones shared between the two sets, we characterised both the codesigned and ProteinMPNN-designed sequences, enabling a controlled comparison on identical structures (Fig. 2b); the remaining 20 ProteinMPNN designs had distinct backbones. Under standard *Escherichia coli* expression, 41 of 48 constructs expressed solubly; for the 38 without significant aggregation, size-exclusion chromatography (SEC) with multi-angle light scattering (MALS) confirmed the expected monomeric mass, and circular dichroism (CD) indicated folded, highly thermostable proteins with melting temperatures above 95 °C (Supplementary Figs. S1–S3). Overall, codesigned and ProteinMPNN-designed sequences performed comparably in these analyses, providing, to our knowledge, the first experimental validation of a generative codesign method whose direct sequence outputs match the performance of ProteinMPNN without post hoc inverse folding.

**Fig. 2.**
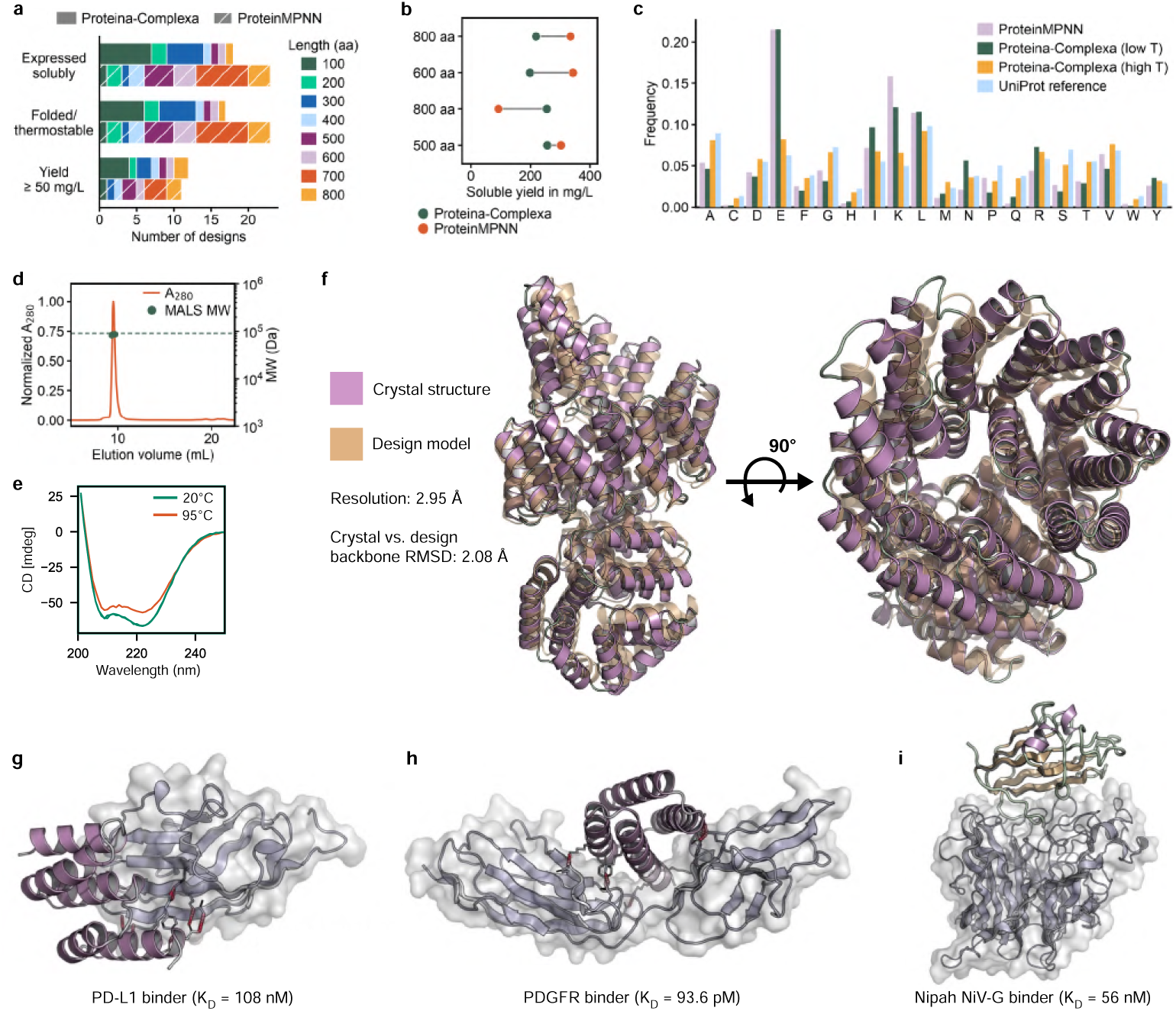
Designed monomers and protein binders. **a**, Outcomes for 48 designed monomers (100–800 aa): soluble expression, thermostable folding and yield, comparing codesigned (Proteína-Complexa) and ProteinMPNN-designed sequences. **b**, Expression yield comparison between Proteína-Complexa codesigned and ProteinMPNN-designed sequences for four shared backbones. **c**, Amino-acid composition of codesigned sequences (Proteína-Complexa, low and high sampling temperature) versus ProteinMPNN-designed sequences and a UniProt reference. **d**, Size-exclusion chromatography (SEC) profile overlaid with the molecular weight determined by multi-angle light scattering (MALS) for a representative 800-residue design. **e**, Circular dichroism (CD) spectra at 20°C and 95°C for the same design. **f**, Crystal structure of the same design (purple) overlaid on its computational model (wheat). The structure was determined at 2.95 Å resolution and resolved 794 of 800 residues; only loop residues 311–316 were unresolved. The backbone RMSD between the resolved experimental structure and the design model is 2.08 Å; per-residue RMSD in Supplementary Fig. S5. **g**, PD-L1 binder in complex; target represented with transparent surface (10.4% hit rate; *K*_D_ 108 nM–3.71 *µ*M). **h**, PDGFR binder in complex (63.5% hit rate; *K*_D_ 93.6 pM–1.34 *µ*M). **i**, De novo Nipah NiV-G binder in complex (*K*_D_ 56 nM).

Proteína-Complexa’s sampling temperature provided tunable control over sequence character: lower temperatures yielded ProteinMPNN-like, readily expressible sequences, whereas higher temperatures produced amino-acid distributions closer to those of natural proteins (Fig. 2c). Among 13 high-temperature designs selected for low sequence identity (≤ 40%) and MSA-supported structure predictions, two expressed solubly, and both showed cooperative thermal transitions below 95,°C (Supplementary Fig. S4). These data suggest a temperature-dependent trade-off between the hyperstability typical of conventionally designed proteins and more natural-like sequence composition and thermal behaviour.

To confirm that the generated backbones are physically realizable and not merely high-scoring in silico, we determined a 2.95 Å-resolution crystal structure of a representative 800-residue design, a length at which most design methods fail to yield successful samples. The structure resolved 794 of 800 residues (99.3%), with only loop residues 311–316 unresolved, and matched the computational model with a backbone RMSD of 2.08 Å across the resolved residues (SEC–MALS results and CD spectra for the crystallised design in Fig. 2d,e; structure in Fig. 2f), to our knowledge the first time a generative model has produced such structures and sequences via codesign.

## 4 Protein binders

PDGFR [31, 32] and PD-L1 [12] provided our first direct tests of Proteína-Complexa’s binder-design capabilities. PDGFR signalling drives proliferative and fibrotic disease, and its extracellular domain presents the kind of polar, solvent-exposed surface that contact-driven design has struggled to engage; PD-L1 is a central immune-checkpoint target in oncology and a common benchmarking target for binder-design pipelines. Interface hotspots were identified by Rosetta alanine scanning of known complexes, and several hotspot combinations were evaluated per target rather than committing to a single in-silico prediction. For each combination, 1,500 candidates were generated by beam search with an AlphaFold2 interface-pAE reward and passed through a two-stage filter on physicochemical and confidence metrics followed by structural validation (Methods); 192 designs per target were then expressed and tested. Of these, 191 (PDGFR) and 192 (PD-L1) expressed and purified, and surface plasmon resonance identified 122 designs with detectable binding to PDGFR and 20 to PD-L1. Measured affinities spanned 93.6 pM to 1.34 *µ*M for PDGFR and 108 nM to 3.71 *µ*M for PD-L1, corresponding to hit rates of 63.5% and 10.4%, thereby experimentally validating Proteína-Complexa’s binder-design capabilities within the latent generative search framework with sequence-structure codesign. Representative PD-L1 and PDGFR complexes are shown in Fig. 2g,h.

Having established binder capability on PDGFR and PD-L1, we turned to the Nipah virus attachment glycoprotein (NiV-G) as a harder test case and a use case for scaffold-based re-engineering. NiV-G is a World Health Organization priority pathogen target with no approved therapeutics, and its receptor-binding site is recessed and difficult to engage [33]. We tested two modes within the same framework. The first was fully de novo design, guided by a multi-objective reward combining Boltz-2 interface confidence with shape complementarity, interface hydrophobicity and hydrogen-bond satisfaction [34]. The second was codesign-based scaffold re-engineering, in which an existing binder is partially noised and a related backbone and sequence are codesigned; this capability follows naturally from generating sequence and structure together and contrasts with fixed-backbone redesign, which can only change the sequence. All 20 designs were expressed, purified and screened by surface plasmon resonance: the de novo set yielded a binder at *K*_D_ = 56 nM, and five of six re-engineered designs bound with nanomolar affinity, all targeting the receptor-binding site (Fig. 2i). The same framework therefore supports both fully de novo design and the optimisation of existing binders without any change of method.

## 5 Benchmarking design methods at scale

The individual campaigns above show that our Proteína-Complexa-based latent generative search framework produces folded proteins and high-affinity binders on selected targets. We next asked how it behaves at scale and how it compares with other methods under matched computational resources. Binder design methods are typically reported on a handful of targets, often with bespoke filtering and compute budgets, which makes head-to-head comparisons difficult and can overstate generality. We therefore designed a benchmark large enough to compare methods under standardized conditions and to probe specificity directly. A 127-target panel was assembled from targets used in publicly reported binder-design studies and benchmarks, including those associated with RFdiffusion, BindCraft and mBER, maximizing overlap with established test cases while spanning diverse protein classes and surface geometries [13, 14, 35].

In total, we screened 1,015,870 designed sequences, generated across more than 1,000 design tasks, against 127 targets by multiplexed phage display. The screened library comprised three non-overlapping subsets dedicated to distinct analyses: 492,636 Proteína-Complexa sequences spanning multiple hotspot combinations for panel-wide target and epitope coverage; 390,569 sequences from all evaluated method and sequence-design configurations for a compute-matched comparison; and 132,665 additional Proteína-Complexa sequences generated at alternative sampling temperatures. All three subsets were assayed in a single experimental campaign.

Each Proteína-Complexa target–hotspot task used standard beam search guided by an AlphaFold2 interface-pAE reward, followed by confidence, clash and structural-diversity filtering (Supplementary Fig. S6). Crucially, the multiplexed format resolves both on-target activity and off-target engagement across the whole panel simultaneously, providing a view of specificity, rather than on-target activity alone, that is inaccessible in small-scale studies where each binder is tested against its intended target alone. Generation and in silico filtering across the campaign required more than 140„000 GPU hours, demonstrating the deployment of the framework at a large computational scale.

High-throughput screening of the 492,636 Proteína-Complexa-generated sequence subset across the 127-target panel with diverse epitopes revealed broad target coverage, with binding events strongly enriched toward the intended targets (Fig. 3a). Proteína-Complexa produced at least one on-target hit for 115 of 127 targets (91%); 92 targets (72%, Fig. 3b) yielded target-specific binders and 78 yielded poly-specific binders engaging two to four targets; these categories were not mutually exclusive. Counting each binder once, 1,982 of 23,966 binders with detected activity (8.3%) engaged their intended target, a 10.5-fold enrichment over the 0.79% expected under uniform binding across the panel. Despite this on-target enrichment, off-target binding was also widespread, with every panel target specifically bound by at least one sequence designed against another target, indicating that a single design campaign can yield binding activity well beyond the intended targets. The choice of interface hotspot used to condition generation had a substantial effect on success, and the computationally top-ranked hotspot did not always give the highest experimental hit rate (Supplementary Fig. S7), supporting the evaluation of several hotspot combinations per target rather than a single in silico prediction. The results demonstrate the effectiveness of latent generative search for binder design at scale across diverse targets.

**Fig. 3.**
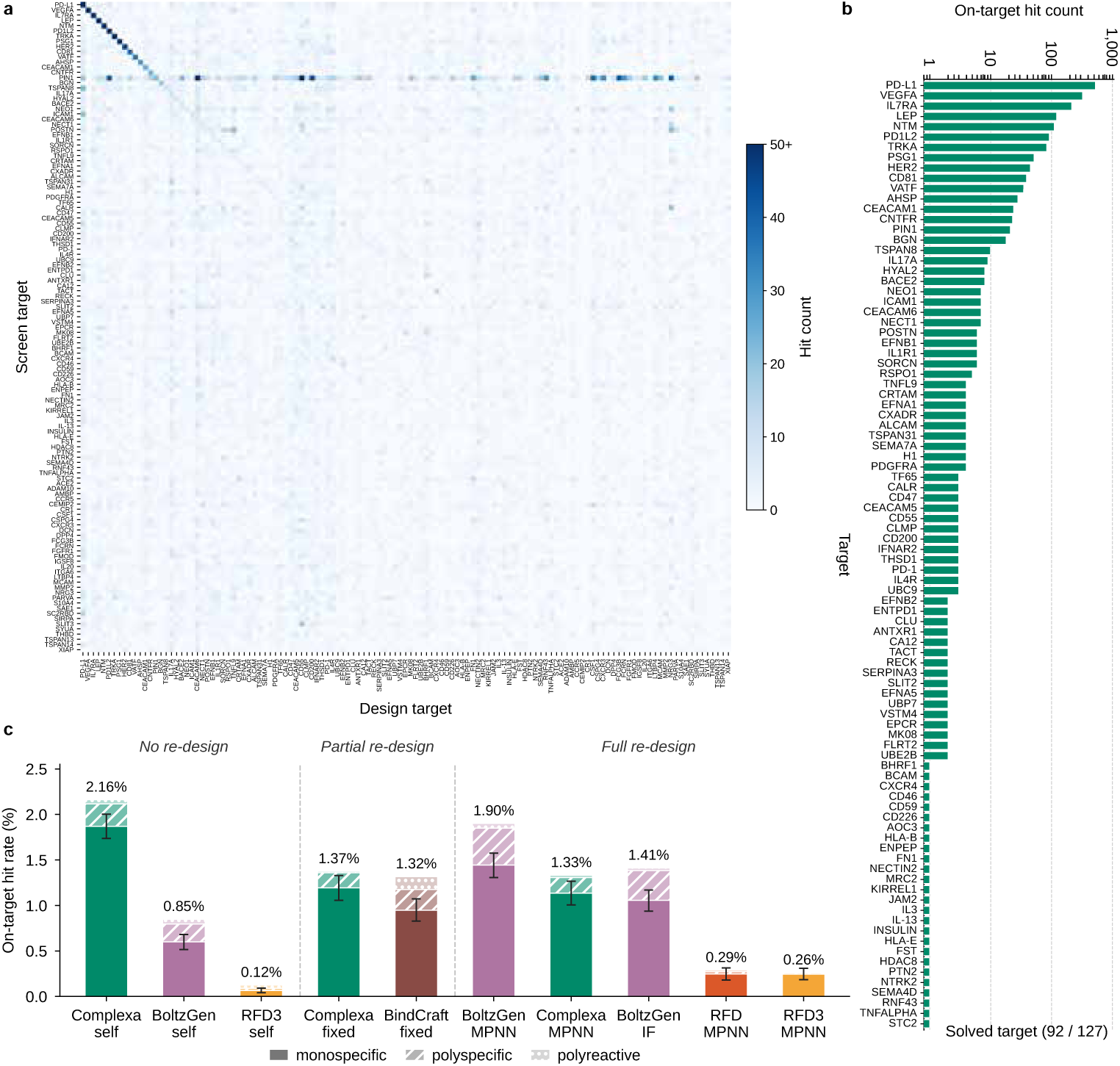
Benchmarking designs at scale against 127 targets by multiplexed phage display. **a**, Cross-reactivity matrix of hit counts for 492,636 Proteína-Complexa-generated sequences (design target, columns) screened against all 127 targets (screen target, rows); on-target hits lie on the diagonal and off-target (cross-reactive) hits off it. **b**, On-target specific hit count for each target, ranked; 92 of 127 targets were solved (at least one on-target specific hit). **c**, On-target specific hit rate across design methods (Proteína-Complexa with latent generative search, BoltzGen, RFdiffusion3, BindCraft, RFdiffusion) and sequence design strategies for the matched-compute comparison sequence set, grouped by native model output (no post hoc redesign), partial redesign (fixed-interface) and full redesign (ProteinMPNN/inverse folding); bars are split into monospecific, poly-specific and poly-reactive fractions.

We then compared methods by specific on-target hit rate in a compute-matched analysis restricted to the top-ranked in silico hotspot for each of the 83 targets for which at least one method produced an on-target hit (Fig. 3c). This analysis used the dedicated subset of 390,569 sequences comprising designs from all approaches. For each target, every method was assigned the same candidate budget, and generation was performed under comparable computational budgets. Using codesigned sequences, Proteína-Complexa achieved a specific hit rate of 1.87%, exceeding both the next-best method with native model sequences, BoltzGen (0.60%), and the best method with fully redesigned sequences, BoltzGen with ProteinMPNN (1.44%) [30, 36]. In absolute terms, Proteína-Complexa yielded 702 on-target hits, of which 607 were specific, from 32,469 codesigned sequences. The corresponding on-target hit counts (specific hits in parentheses) were 511 (388) for BoltzGen with ProteinMPNN, 323 (232) for BindCraft [14], 61 (57) for RFdiffusion3 [29] and 61 (52) for RFdiffusion [13]. The sequence-design strategy mattered as much as the backbone-generation method. For Proteína-Complexa, codesigned sequences (702 on-target hits, 607 specific; 1.87% specific hit rate) outperformed both ProteinMPNN redesign (323 hits, 275 specific; 1.13%) and fixed-interface redesign (333 hits, 289 specific; 1.19%) on the same backbones. This ordering did not hold for the other methods: BoltzGen performed best with ProteinMPNN redesign (1.44%) rather than its native sequences (0.60%), and RFdiffusion3 native sequences reached only 0.07%. In this benchmark, Proteína-Complexa with our latent generative search framework was therefore the only method whose codesigned sequences consistently exceeded post hoc redesign strategies, indicating that codesign can reduce reliance on separate inverse-folding models at scale. In silico confidence metrics were predictive of experimental success, with hits enriched among designs with high interface confidence and pLDDT (Supplementary Fig. S8). Finally, sampling temperature ablations incorporating the remaining subset of 132,665 sequences revealed complementary, target-dependent performance across settings, supporting the use of multiple temperatures to broaden target coverage (Supplementary Table S1).

## 6 Binders to a muscle-wasting receptor

Activin receptor type IIA (ActRIIA) suppresses skeletal-muscle growth through Smad2/3 signalling and is a target in muscle-wasting disorders, including the lean-mass loss that can accompany the GLP-1 receptor agonist therapies now in widespread use [37]. A designed mini-protein that blocks this receptor could help preserve muscle during weight loss, but doing so demands both high affinity and the ability to interrupt downstream signalling in living cells (Fig. 4a).

**Fig. 4.**
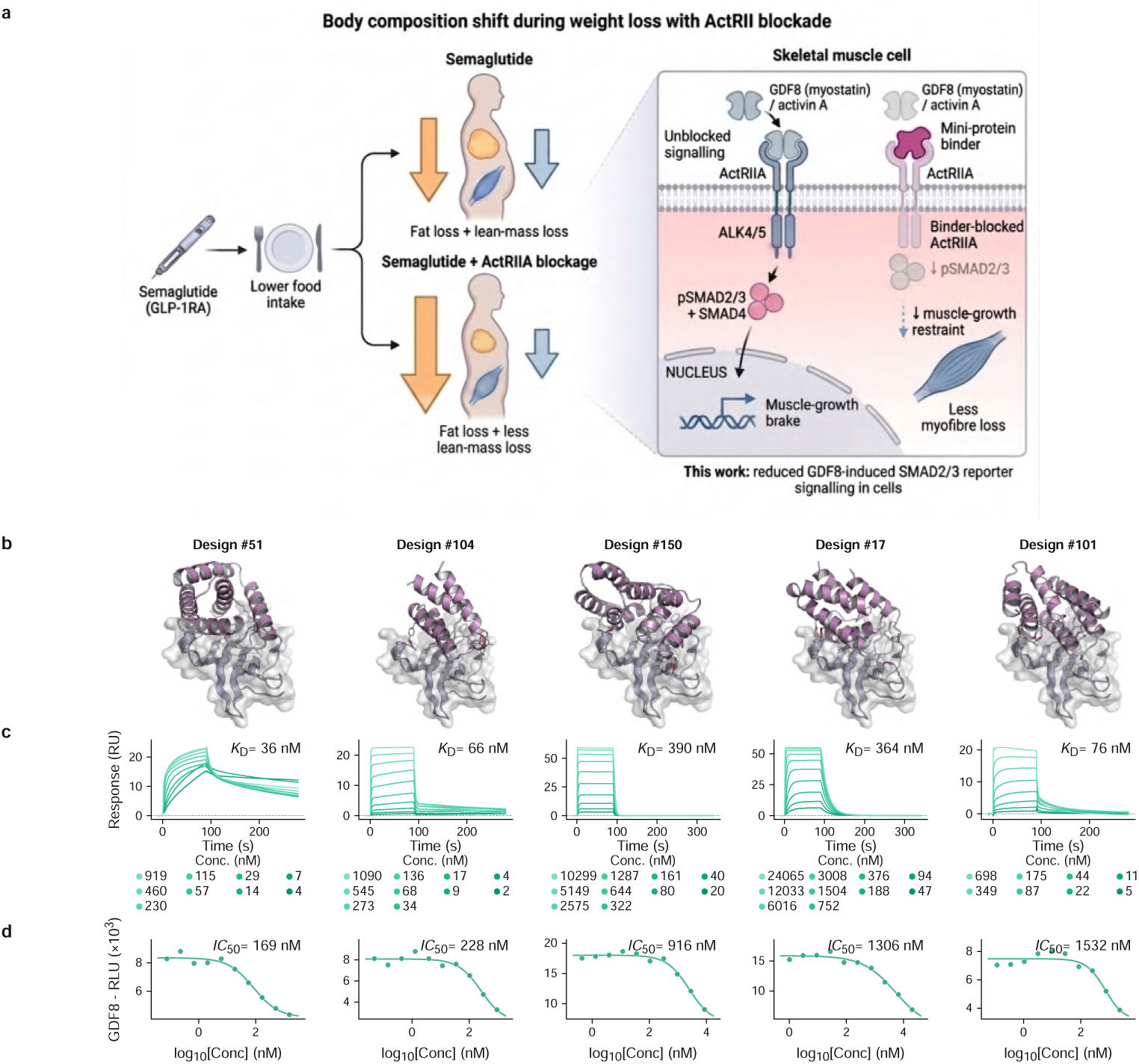
De novo binders to the muscle-wasting receptor ActRIIA. **a**, Rationale for ActRIIA blockade during semaglutide-induced weight loss and schematic of binder-mediated inhibition of GDF8/Activin signalling through Smad2/3. **b**, Predicted structures of five representative designs (#51, #104, #150, #17 and #101; purple) bound to ActRIIA (grey); the binders share a helical binding mode at low pairwise sequence identity (6–18%). **c**, Surface plasmon resonance sensorgrams (analyte concentration series, legends in nM) with fitted *K*_D_ (36–390 nM; tightest design 36 nM). **d**, Inhibition of GDF8 (myostatin)-induced Smad2/3 luciferase signalling in cells, shown as GDF8-driven luminescence versus binder concentration, with fitted IC_50_ (169–1,532 nM).

Across seven *in silico* design rounds we varied sampling size, hotspot conditioning and the search algorithm — beam search, Feynman–Kac steering or Monte Carlo tree search. Hotspots were drawn from the ActRIIA–bimagrumab and ActRIIA–Activin A interfaces, and candidates were ranked by an averaged complex interface score before scaffold clustering. Of 200 selected designs, 192 expressed and purified. Surface plasmon resonance identified 16 binders, 8 with sub-micromolar affinity and a tightest design at *K*_D_ = 36 nM (Fig. 4c). The sub-micromolar hits shared low sequence identity (6–18%) despite a common helical binding mode (Fig. 4b), again pointing to diverse solutions rather than a single convergent design. Crucially, function tracked affinity: in a Smad2/3 luciferase reporter assay in HEK293 cells, designs inhibited myostatin-induced signalling with half-maximal inhibitory concentrations from 169 to 1,532 nM (Fig. 4d). That computationally designed mini-proteins disrupt receptor signalling in cells — directly from design and without any experimental affinity maturation — shows that the latent generative search framework yields not just binders but functional antagonists of a therapeutically important receptor.

## 7 Peptide and mini-protein kinase binders

Most de novo binder design has focused on mini-proteins of roughly 50–100 residues, where a folded scaffold provides a rigid, preorganised interface. Shorter peptides are both biologically and therapeutically attractive — they are synthetically accessible and can reach buried or constrained sites — but they are far harder to design: with fewer residues and a smaller, more flexible interface, sequence and backbone geometry must be specified together with little tolerance for error, exactly the regime in which codesign should be beneficial. To test generality across this size spectrum, we designed binders to the catalytic domains of two kinases: mini-protein binders (49–74 amino acids) for PAK1 [38, 39] and short peptide binders (*<*31 amino acids) for CK1*δ* [40, 41] (Fig. 5a,b), choosing different length windows for both targets to probe the capabilities of the method in different scenarios.

**Fig. 5.**
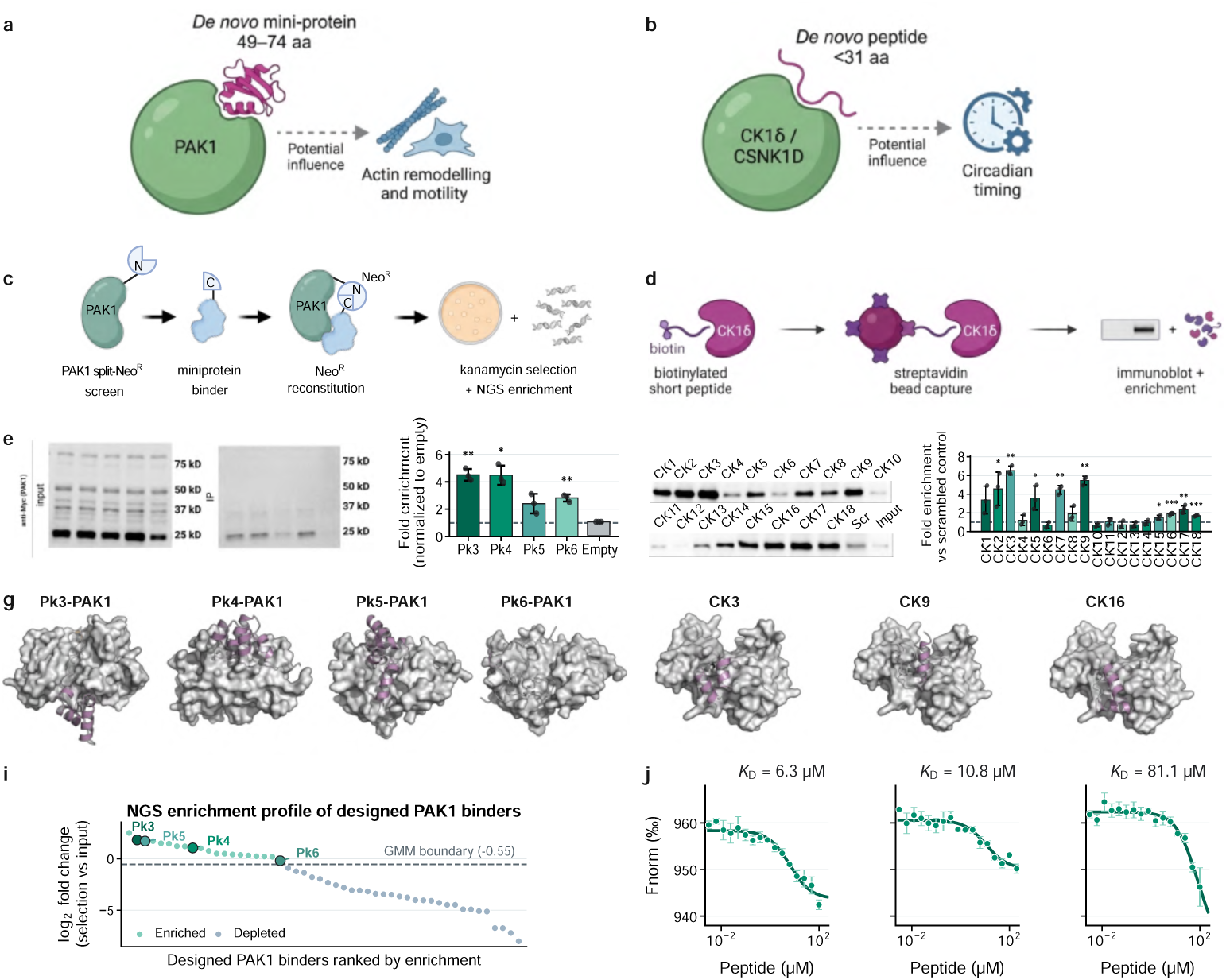
De novo peptide and mini-protein kinase binders. The left column concerns PAK1 mini-proteins (**a**,**c**,**e**,**g**,**i**) and the right column CK1*δ* peptides (**b**,**d**,**f**,**h**,**j**). **a**, Signalling context for PAK1, a Rho-family GTPase effector, and the 49–74-aa mini-protein design regime. **b**, Signalling context for CK1*δ*, a constitutive Ser/Thr kinase, and the *<*31-aa peptide design regime. **c**, Split-neomycin-resistance complementation assay for PAK1 mini-proteins, read out by next-generation sequencing (NGS). **d**, Streptavidin-bead pull-down assay for biotinylated CK1*δ* peptides. **e**, Mammalian validation of PAK1 binders by representative anti-Myc Western blot and fold enrichment relative to empty vector (mean *±* SD, *n* = 3). **f**, CK1*δ* pull-down blots and binding enrichment for CK1–CK18 relative to scrambled control (mean *±* SD, *n* = 3–4); 9 of 18 are significant (*p <* 0.05). **g**, AlphaFold2-Multimer predicted PAK1 complexes for Pk3–Pk6 (PAK1 surface, binder cartoon); details in Supplementary Table S2. **h**, Predicted CK3, CK9 and CK16 peptide–CK1*δ* complexes. **i**, Split-NeoR NGS enrichment profile for the 50 PAK1 designs, ranked by fold change, with the Gaussian-mixture decision boundary and Pk3–Pk6 indicated. **j**, Microscale-thermophoresis dose– response curves for CK3, CK9 and CK16 (*K*_D_ = 6.3, 10.8 and 81.1 *µ*M, respectively; mean *±* SEM, *n* = 3–4).

For PAK1, 50 mini-protein candidates were generated by beam search with an AlphaFold2 interface-pAE reward and screened by a split-neomycin-resistance complementation assay in *E. coli* [42, 43] (Fig. 5c). Next-generation sequencing classified 20 of 50 designs (40%) as enriched (Fig. 5i), and co-immunoprecipitation in HEK293T cells confirmed binding for top candidates, with the strongest designs reaching roughly 4.5-fold enrichment over control (*P <* 0.01; Fig. 5e). The corresponding predicted PAK1 complexes show distinct binding modes (Fig. 5g). For CK1*δ*, 18 synthesised peptides were assayed by streptavidin pull-down against the purified kinase (Fig. 5d); 9 of 18 (50%) showed significant enrichment over a scrambled control (*P <* 0.05), with the strongest binders exceeding five-fold enrichment (Fig. 5f); all binders in Supplementary Table S3. Predicted CK3, CK9 and CK16 complexes are shown in Fig. 5h, for which we measured low-to-mid micromolar dissociation constants by microscale thermophoresis (Fig. 5j). High success rates in both regimes indicate that codesign generalises across both peptide and protein binders. The binding affinities of CK3, CK9, and CK16 for CK1*δ* were determined by microscale thermophoresis using a Monolith X instrument (NanoTemper Technologies). Normalized fluorescence (*F*_norm_) was measured across a 16-point serial dilution series and fitted to a three-parameter sigmoidal binding model to derive dissociation constants (*K*_d_; Fig. 8e). CK3 exhibited a *K*_d_ of 6.27 ± 1.81 *µ*M, CK9 a *K*_d_ of 10.78 ± 2.83 *µ*M, and CK16 a *K*_d_ of 81.06 31.48 *µ*M (mean ± SD). Together, these data demonstrate that Proteina-Complexa-predicted peptide binders engage the CK1*δ* kinase domain with low-to-mid-micromolar affinity.

## 8 De novo carbohydrate-antigen binders

Carbohydrates carry much of the information at the human cell surface — governing blood-group identity, host–pathogen defence and immune signalling — yet they remain almost untouched by de novo design. The ABO system is the canonical example: the A antigen terminates in N-acetylgalactosamine (GalNAc), whereas the B antigen terminates in galactose (Gal), and this single chemical difference is a primary determinant of blood-transfusion and organ-transplant compatibility [44]. In reverse blood typing, patient red blood cells are mixed with antibodies of known specificity and antigen-dependent agglutination reveals the blood group. These antibodies are exquisitely specific but large (*>*150 kDa), thermally fragile and costly to produce. Smaller carbohydrate-binding molecules bind both antigens without sufficient discrimination, typically with only tens-to-hundreds-of-micromolar affinity, and often require structural metal ions, making them poor typing reagents. A small, hyperstable protein that binds one antigen specifically would enable cheap, cold-chain-free blood-typing reagents for point-of-care and field use, and more broadly a route to programmable carbohydrate recognition. Designing such a binder, however, requires recognition of a small, flexible, densely polar sugar — a hydroxyl-rich surface with little hydrophobic character and a substantial desolvation penalty — such that even natural lectins bind free sugars only weakly [45, 46]. These are exactly the properties that defeat contact-driven design and ProteinMPNN [12]; instead, sequence and backbone geometry must be co-optimised to place a hydrogen-bond network precisely around the ligand. We are not aware of a prior de novo protein that binds a free carbohydrate with measurable affinity (Fig. 6a).

**Fig. 6.**
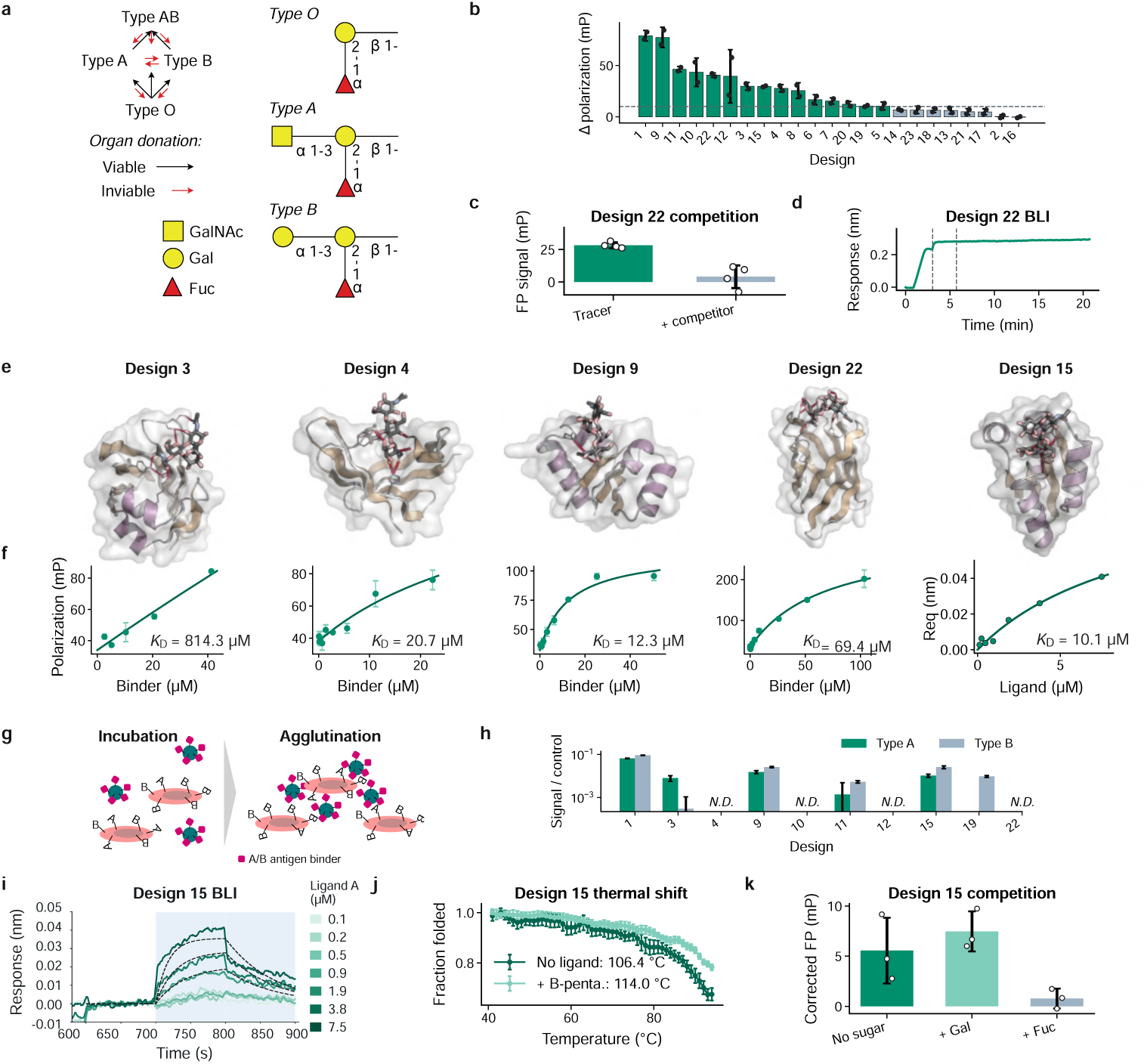
De novo binders that recognise a blood-group carbohydrate antigen. **a**, Principle of ABO blood-group compatibility and schematic structures of the A, B and O(H) antigens; the A antigen terminates in N-acetylgalactosamine (GalNAc), whereas the B antigen terminates in galactose (Gal) (glycan symbols per standard glycomics nomenclature). **b**, FP screen of designs 1–23 against fluorescently labelled type II A pentasaccharide. Δpolarization is relative to the tracer-only control; bars show duplicate means and error bars show s.d. Fifteen designs pass the +10 mP threshold. **c**, Excess unlabelled type II A competes design 22 binding to the fluorescent probe back to baseline. **d**, BLI trace of design 22 against immobilised biotinylated type II A glycan, with association and dissociation phases indicated. **e**, Predicted glycan-bound structures of designs 3, 4, 9, 15 and 22. **f**, Concentration-dependent binding of the same five designs, measured by FP (designs 3, 4, 9 and 22) or BLI equilibrium response (design 15), with fitted *K*_D_; the design 3 estimate lies above the measured concentration range. **g**, Agglutination assay in which His_6_-tagged binders are immobilised on Ni-NTA agarose beads to confer multivalency before incubation with red blood cells. **h**, Type A_2_ and type B red-blood-cell agglutination relative to blank-bead controls (error bars, s.d. of triplicate wells; N.D., no agglutination above the negative control). Design 3 agglutinates A_2_ but not B. **i**, BLI of immobilised design 15 against a type II A pentasaccharide–DNA conjugate, reference-subtracted against a buffer-only control. **j**, Circular-dichroism thermal shift of design 15 with and without type B pentasaccharide. **k**, FP competition for design 15: 30 mM unlabelled D-galactose does not compete, whereas 250 *µ*M L-fucose abolishes the signal (error bars, s.d. of triplicate measurements).

Because carbohydrate recognition depends on precise polar contacts while sugars remain conformationally flexible, we adapted latent generative search to address both properties explicitly. We conditioned Proteína-Complexa on the type II A pentasaccharide and used pocket-burial terms together with physics-based Tmol hydrogen-bond rewards during search to favour geometrically precise polar interactions [25]. At every search step, RosettaFold3 (RF3) predicted the bound complex, and the RF3-predicted sugar conformation was fed back into the next generative step. The search therefore updated the ligand pose as the binder evolved, accounting for binding-dependent glycan flexibility rather than treating the sugar as rigid [27].

We expressed 23 designs, each smaller than 10 kDa, and screened the purified proteins against a fluorescently labelled type II A pentasaccharide by fluorescence polarization (FP). Fifteen designs exceeded a +10 mP threshold (Fig. 6b). Excess unlabelled pentasaccharide competed the FP signal back to baseline (Fig. 6c), and biolayer interferometry (BLI) against immobilised glycan, referenced against non-specific sensor binding, showed a clear on-rate and a very slow off-rate, confirming glycan binding independently of the fluorescent tracer (Fig. 6d). Concentration-series measurements were obtained for ten putative binders and yielded five well-constrained dissociation constants of approximately 12–340 *µ*M in the initial FP analysis. Predicted complexes illustrate the pockets formed around the glycan (Fig. 6e). Five representative designs are shown in Fig. 6f: designs 3, 4, 9 and 22 measured by FP and design 15 measured by BLI equilibrium response. Their fitted *K*_D_ values span 10.1–814.3 *µ*M; the design 3 value is an extrapolated estimate because the fitted *K*_D_ lies above the measured concentration range.

To ask whether these binders can type red blood cells, we conferred multivalency on the His-tagged designs by immobilising them on Ni-NTA beads and measured agglutination of type A_2_ or type B cells by OD_600_ (Fig. 6g). Design 3 agglutinated type A_2_ cells but not type B, discriminating between the two antigens (Fig. 6h); this is a demanding test, because A_2_ cells display the A antigen at low density while the type B cells present a competing, higher-valency surface. Five further designs (1, 9, 11, 15 and 19) agglutinated type A and/or type B cells without a clear preference.

We used design 15, which agglutinated both cell types, as a case study in poly-specificity. It bound the A antigen by BLI (Fig. 6i) and was thermally stabilised by the B pentasaccharide (Fig. 6j), pointing to recognition of a feature shared by both antigens. Competition experiments localised that feature: free galactose (30 mM) did not inhibit FP, whereas L-fucose (250 *µ*M) abolished it (Fig. 6k), and the model predicts a hydrophobic stacking contact between design 15 and the fucose ring. Because fucose is common to the A, B and O antigens, this off-target interaction with a shared monosaccharide compresses the dynamic range of the cell assay — a reminder that screening in the relevant cell type is essential to reveal biologically important off-target effects.

Design 3, by contrast, distinguishes A_2_ from B. Its model places a tyrosine on a *β*-strand to hydrogen-bond the terminal N-acetyl group unique to the A antigen — an interaction that cannot form with the galactose of the B antigen — providing a structural rationale for the observed specificity, and it contacts fucose through only two hydrogen bonds rather than a stacking interaction. Design 3 has no detectable sequence homologue in the non-redundant database and no structural relative by Foldseek (no hit at E *<* 10*^−^*^2^) [47], indicating that Proteína-Complexa reached a region of sequence and fold space without a known natural non-antibody solution to A/B discrimination. A complementary BLI screen of the FP-positive designs against type-A and type-B glycans is consistent with antigen-preferential binding (Supplementary Fig. S9).

## 9 Discussion

The results show that codesign with reward-guided search in a continuous latent space produces functional binders across a wide range of targets and scales. A large-scale screening campaign testing over a million binder candidates demonstrated broad on-target success across a diverse panel of 127 targets. In what is, to our knowledge, the largest experimentally validated benchmark of protein binder design methods to date, our codesign-based latent generative search framework outperformed competing generative models, an optimisation baseline and post hoc redesign under compute-matched conditions, with the advantage over sequence redesign indicating that generating backbone and sequence together yields measurable experimental gains. Individual campaigns further demonstrate the breadth of the approach across modalities: picomolar binders to PDGFR; ActRIIA binders that block signalling in cells without affinity maturation; nanomolar Nipah virus binders by de novo design and scaffold re-engineering; kinase binders across the peptide-to-protein size range; and de novo binders to a free carbohydrate, including a design that discriminates between blood-group antigens.

Open questions remain. Codesign outperforms sequence redesign on average, but the margin varies across targets, and which interface properties benefit most from joint optimisation is not yet clear. The carbohydrate results motivate extension to other non-protein modalities. More broadly, latent generative search — our framework pairing a learned generative prior with adaptive inference-time search — translates a principle established in other areas of artificial intelligence into experimentally validated protein design across target classes that have been difficult to address. With an expanded repertoire of scoring functions and optimisation objectives, latent generative search could unlock further new classes of protein design problems. Code and model weights are released to support reproduction and extension (Methods Section 10).

## 10 Methods

### Latent generative search and base model

The base generative model, Proteína-Complexa, and the inference-time search algorithms used here were introduced previously [18]; we summarise them for completeness and refer to that work and to La-Proteína [19] for full specifications. The model builds on La-Proteína’s partially latent representation, in which backbone C*_α_* coordinates are modelled explicitly and each residue’s amino-acid identity and remaining atoms are encoded into a continuous latent variable by a variational autoencoder; a rectified-flow flow-matching model generates backbone coordinates and latents jointly [20]. The denoiser is a transformer with pair-biased attention over a concatenated sequence of noisy binder tokens (C*_α_* coordinates and latents) and clean target tokens (Atom37 coordinates, amino-acid identity and binary hotspot tokens); target conditioning is applied to the flow model only, leaving the autoencoder target-agnostic. During training the binder C*_α_* coordinates are perturbed with a global translation (*c_d_*= 0.2 nm) so that the model reasons over binder placement. We use separate models for protein-protein interactions and for binder design to ligand targets. Extended model details are presented in Supplementary S1; full architecture and training hyperparameters are given in ref. [18] (its Appendix G, Table 14) and are reproduced in Supplementary Table S4.

### Training data

Four datasets are used [18]: Foldseek AFDB monomer cluster representatives [47]; Teddymer dimer cluster representatives (filtered for interface pLDDT *>* 70, interface predicted aligned error *<* 10 and interface length *>* 10); protein multimers filtered from the PDB [23]; and the filtered PLINDER protein–ligand dataset [48]. Teddymer is derived from AFDB50 structures with TED domain annotations [22, 49] (47M structures), split into domains, filtered for spatial proximity and complete CATH annotation [50] to give 10M dimers, then clustered to 3.5M representatives [21]. The variational autoencoder is pretrained on AFDB monomers (500k steps, 16 GPUs) and fine-tuned on 110,976 PDB chains (length 50–256, resolution better than 5.0 Å; 40k steps). The flow model is pretrained on AFDB monomers and then trained on Teddymer and PDB multimers for protein targets; small-molecule targets use PLINDER and AFDB monomers with low-rank adaptation [51].

### Inference-time search

We use best-of-*N* sampling, beam search, Feynman–Kac steering and Monte Carlo tree search adapted to flow models [52–55]. Candidate states are rolled out to clean structures, decoded, folded and scored, with the search applied every *K* denoising steps. Default hyperparameters are beam width *N* = 4, branching factor *L* = 4 and step length *K* = 100; Feynman–Kac steering additionally uses inverse temperature *β* = 10 [18]. Scaffold re-engineering (Nipah) uses a diffuse–denoise procedure that partially noises an input binder and codesigns a related backbone and sequence.

### Reward functions and success criteria

Search and filtering use combinations of complementary rewards (Supplementary Table S5). In-silico success for protein targets follows AlphaFold2-Multimer interface metrics (pLDDT *>* 90, interface predicted aligned error *<* 7.0, binder RMSD *<* 1.5 Å) via ColabDesign [26, 56]; small-molecule and carbohydrate targets use RosettaFold3 (minimum interface predicted aligned error *<* 2, binder RMSD *<* 2 Å, ligand RMSD *<* 5 Å) [27]. Hydrogen-bond energies are computed with the Rosetta energy function via Tmol and hydrogen bonds detected with HBPlus [24, 25, 57]. Per-campaign reward mixtures were as follows. PDGFR/PD-L1, the massive-scale benchmark and PAK1 used beam search with an AlphaFold2 interface-pAE reward. Nipah de novo design used beam search with a Boltz-2 maximum interface score combined with shape complementarity, interface hydrophobicity and hydrogen-bond satisfaction [34]. ActRIIA used averaged complex interface scores for ranking across beam search, Feynman–Kac steering and tree search. Carbohydrate design used pocket-burial and physics-based Tmol hydrogen-bond rewards together with RosettaFold3 interface confidence. After every search step, the RosettaFold3-predicted bound sugar conformation replaced the preceding ligand pose before the next generative step, allowing the search to account for binding-dependent glycan flexibility.

### Designed monomers

Designs were filtered by AlphaFold2 single-sequence prediction (mean pLDDT *>* 90, RMSD *<* 2 Å to the design model). Genes were codon-optimised, cloned by Golden Gate assembly and expressed in *E. coli* BL21(DE3) by auto-induction. Proteins were purified by Ni-NTA affinity and size-exclusion chromatography (SEC); yields were determined from A_280_. Oligomeric state was assessed by SEC coupled to multi-angle light scattering, and secondary structure and thermostability by circular dichroism (25–95 °C). A representative design was crystallised and its structure solved by X-ray crystallography.

### PDGFR and PD-L1

Interface hotspots were identified by Rosetta alanine scanning of known complexes (retaining positions with favourable binding contribution; top nine per target). For each hotspot combination, 1,500 candidates (55–95 aa) were generated by beam search with an AlphaFold2 interface-pAE reward. Stage-one filtering used isoelectric point, ESM pseudo-likelihood, minimum interface-pAE, complex interface-pTM and pLDDT, and cysteine count, followed by TM-score clustering at 0.8; stage two validated monomeric structures with OpenFold3 and Boltz-2 (pLDDT *>* 80, RMSD *<* 2 Å). 192 designs per target were expressed in *E. coli*, purified, and assayed by surface plasmon resonance with kinetic or steady-state fitting of *K*_D_.

### Nipah virus

De novo designs were generated conditioned on hotspots at the NiV-G receptor-binding site using beam search with the multi-objective reward above; scaffold re-engineering used the diffuse–denoise procedure. Designs were expressed in *E. coli* and purified by Ni-affinity and size-exclusion chromatography, and binding was measured by surface plasmon resonance against NiV-G with *K*_D_ from kinetic or steady-state fitting.

### Massive-scale benchmark

Targets were sourced commercially (Supplementary Table S6). For each target, design tasks spanned multiple hotspot combinations (about ten where literature hotspots were unavailable), with an overall screening budget of 10,000 sequences per target. Per task, a single Proteína-Complexa run with standard beam search and an AlphaFold2 interface-pAE reward produced about 27,000 candidates, filtered to the top 1,000 by predicted aligned error; in-silico success required minimum interface-pAE *<* 2, complex pLDDT *>* 0.8, interface-pTM *>* 0.7 and fewer than five C*_α_* clashes. For the screening of diverse Proteína-Complexa-generated sequences across all targets and hotspots, diversity selection clustered candidates by predicted structure and cycled ranking criteria across clusters and hotspots (Supplementary Fig. S6).

For the separate baseline comparison experiment using the top-ranked in silico hotspots only both Proteína-Complexa with latent generative search and the baseline methods (BindCraft, RFdiffusion, RFdiffusion3, BoltzGen with varying sequence design approaches) were run under a matched budget of about 32 GPU hours per target with minimum interface score as the primary metric and eight cycles of soluble ProteinMPNN redesign [13, 14, 29, 30, 36]. For each method, sequence design setting and target, 400 sequences were tested. Here, the final candidates were selected greedily based on minimum ipSAE without clustering (details in Supplementary S4) [58]. The same protocol was used in the sampling temperature ablation experiments. The overall campaign used more than 140,000 GPU hours in total for candidate generation and filtering.

Designs were synthesised as oligo pools, cloned into a phagemid and screened by multiplexed phage display [35]; sequencing reads were merged [59] and counted by exact match. Hits were called by two methods whose union was used: a quantile-ratio test (hit if the minimum across replicates exceeds the upper-tail quantile, *q* = 20) and an enrichment *z*-test on output/input ratios with a robust null (median and median absolute deviation) and Benjamini–Hochberg control of the false-discovery rate (Supplementary S4.2).

### ActRIIA

Hotspots were taken from ActRIIA complexes with bimagrumab and Activin A. Across seven design rounds varying sampling size, search algorithm and hotspot conditioning, candidates were ranked by averaged complex interface score and clustered by scaffold (Foldseek) to select 200 designs. Proteins were expressed in *E. coli* BL21(DE3), purified by Ni-affinity with TEV cleavage and a second Ni step, and assayed by surface plasmon resonance (Biacore) against immobilised ActRIIA over a concentration series in two independent runs. Cellular activity was measured in a Smad2/3-responsive luciferase reporter in HEK293 cells stimulated with Activin A (20 ng mL*^−^*^1^) or GDF8/myostatin (200 ng mL*^−^*^1^), with three-fold serial dilutions of each design.

### Kinase binders

PAK1 mini-proteins (49–74 aa) were generated by beam search with an AlphaFold2 interface-pAE reward and screened by a split-neomycin-resistance complementation assay in *E. coli* [42, 43]; enrichment was computed from next-generation sequencing as log_2_ fold change and classified by a two-component Gaussian mixture model. Selected candidates were validated by HA-Trap co-immunoprecipitation in HEK293T cells with quantitative Western blotting (one-sample *t*-test versus 1). CK1*δ* peptides (*<*31 aa) were synthesised by Fmoc solid-phase synthesis with a biotinylated lysine, captured on streptavidin beads, and assayed for binding to purified GST–CK1*δ* by Western blot (one-tailed *t*-test versus scrambled control). *K*_D_ values for CK1 were determined with microscale thermophoresis on a Monolith X instrument (NanoTemper Technologies) using the 670 nm detection channel.

### Carbohydrate binders

Proteína-Complexa was conditioned on the type II A pentasaccharide. Pocket burial and physics-based Tmol hydrogen-bond energies rewarded deep pockets and precise polar contacts during search. RosettaFold3 was run without ligand templating after every search step; its predicted bound sugar conformation was fed back as the ligand pose for the next step so that binder generation and glycan conformation co-evolved. Twenty-three designs (all *<*10 kDa) were cloned into a T7 vector, expressed in *E. coli* BL21(DE3) and purified by Ni-NTA. Purified proteins were screened against a fluorescently labelled (iFluor488) type II A pentasaccharide by fluorescence polarization (FP) in duplicate; designs exceeding +10 mP over a tracer-only control were counted as hits, and dissociation constants were obtained from FP titration series normalised to the tracer-only control. FP specificity was tested by competition with excess unlabelled type II A pentasaccharide and, for design 15, with 30 mM D-galactose and 250 *µ*M L-fucose. Orthogonal binding was measured by biolayer interferometry, both with protein as analyte over immobilised biotinylated glycan (referenced against an analyte-only control) and with immobilised binder against a glycan–DNA conjugate (pentasaccharide coupled to an amine-modified oligonucleotide by reductive amination; a trivalent analyte poorly described by a 1:1 model). Agglutination was assessed by immobilising His_6_-tagged binders on Ni-NTA agarose beads to confer multivalency, incubating with type A_2_ or type B red blood cells (commercial source) in a plate format, and reading OD_600_ in triplicate against blank-bead controls, with significance from a Z-score. Thermostability and ligand-induced stabilisation were measured by circular dichroism at 222 nm with and without the type B pentasaccharide. Structural and sequence novelty of design 3 was assessed by search against the non-redundant sequence database and by Foldseek structural search [47].

## Code and model availability

Inference code for latent generative search and model weights for Proteína-Complexa are released [18], see https://research.nvidia.com/labs/genair/proteina-complexa/ and https://github.com/NVIDIA-Digital-Bio/proteina-complexa.

## Acknowledgements

We thank our collaborators and funding bodies. Macromolecular crystallography measurements were carried out at beamline MX14.1 at the BESSY II electron storage ring. We would like to thank Melanie Oelker for assistance during the experiment.

## Declarations

### Ethics approval and consent to participate

The agglutination assay used commercially sourced human type-A_2_ and type-B red blood cells; no human or animal subjects were involved.

### Author contributions

K.D.: Conceptualization, Methodology, Software, Investigation, Validation, Formal analysis, Visualization, Data curation, Writing – Original Draft, Writing – Review & Editing, Project administration. D.R.: Methodology, Software, Investigation, Validation, Formal analysis, Visualization, Data curation, Writing – Original Draft, Writing – Review & Editing. M.P. (M. Penner): Methodology, Validation, Formal analysis, Investigation. S.R.: Methodology, Experimental Validation, Data Generation, Investigation. M.C.: Methodology, Formal Analysis, Data Curation, Investigation. M.N.: Conceptualization, Methodology. E.S.: Conceptualization, Methodology. A.R.: Methodology, Data Curation. M.Pr.: Experimental Validation, Data Generation. Y.Q.: Conceptualization, Methodology, Resources, Supervision, Writing – Original Draft. D.Q.: Validation, Data curation. J.Y.: Validation, Data curation. W.L.: Validation, Data curation. L.L.: Formal analysis, Methodology. D.S.: Formal analysis, Investigation, Visualization, Writing – Original Draft. S.G.: Formal analysis, Investigation, Visualization, Writing – Original Draft. B.B.M.: Validation, Data curation, Formal analysis, Investigation, Visualization, Writing – Original Draft. H.C.: Validation, Data curation, Formal analysis, Investigation, Visualization, Writing – Original Draft. L.C.: Validation, Data curation, Formal analysis, Investigation, Visualization, Writing – Original Draft. M.A.J.: Validation, Data curation, Formal analysis, Investigation, Visualization, Writing – Original Draft. H.K.: Validation, Data curation, Formal analysis, Investigation, Visualization, Writing – Original Draft. D.Ri.: Validation, Data curation, Formal analysis, Investigation. P.Sc.: Validation, Data curation, Formal analysis, Investigation. A.U.M.: Formal analysis, Visualization, Writing – Original Draft. M.Se.: Formal analysis, Visualization, Writing – Original Draft. D.G.: Supervision. S.Ch.: Data curation. Z.C.: Software, Investigation, Validation. G.Z.: Software, Investigation, Validation. T.G.: Software, Investigation, Validation. X.C.: Investigation. M.L.: Software, Investigation, Validation. Z.Z.: Methodology, Software, Validation. T.Z.: Project administration. K.G.: Project administration, Writing – Review & Editing. M.M.B.: Supervision. M.St.: Data curation. K.De.: Conceptualization, Resources, Project administration, Supervision, Writing – Original Draft. S.S.: Conceptualization, Methodology, Resources, Supervision. C.T.S.: Conceptualization, Methodology, Resources, Supervision. A.K.: Conceptualization, Methodology, Resources, Supervision, Writing – Original Draft. F.H.: Conceptualization, Methodology, Resources, Supervision. C.D.: Conceptualization, Investigation, Project administration, Writing – Review & Editing. E.K.: Conceptualization, Resources, Supervision, Project administration. A.V.: Resources, Supervision, Project administration. P.O.: Conceptualization, Methodology, Resources, Supervision, Project administration. K.K.: Conceptualization, Methodology, Visualization, Writing – Original Draft, Writing – Review & Editing, Supervision, Project administration.

### Materials availability

Plasmids and designed proteins are available from the corresponding authors on reasonable request. The crystallographic dataset of LP46 has been deposited in the PDB (Accession code: 32GJ).

## Supplementary contents

*S1* Model architecture and latent generative search (p. 32).

*S2* Biophysical characterization of designed monomers (p. 37).

*S3* PDGFR and PD-L1 filtering and hotspot analysis (p. 39).

*S4* Massive-scale benchmark: extended analysis and hit calling (p. 40).

*S5* ActRIIA experimental methods (p. 46).

*S6* PAK1 and CK1*δ* experimental methods (p. 48).

*S7* Carbohydrate-binder experimental methods (p. 51).

*S8* Supplementary figures (p. 54).

*S9* Supplementary tables (p. 70).

## S1 Model architecture and latent generative search

This section gives a self-contained mathematical summary of the model used in this study. It follows the notation of Proteína-Complexa throughout and condenses the full treatments in La-Proteína and Proteína-Complexa [18, 19]. The purpose is to make precise how a complete binder sequence and all-atom structure are generated, how a fixed target conditions that distribution, and how external rewards alter sampling without retraining the base model.

### S1.1 Protein representation and notation

Let a binder contain *N* residues. We write **x***^Cα^∈* ℝ*^N×^*^3^ for its C coordinates, **x***^¬C_α_^ ∈* ℝ*^N×^*^36^*^×^*^3^ for the remaining Atom37 coordinates, and **s** *∈* **{**0*,…,* 19}*^N^* for its amino-acid sequence. Atom37 is a fixed-size storage representation. The residue-dependent mask **m**(**s**) *∈* {0, 1}*^N^*^×36^ indicates which non-C_α_ atom slots are present for each residue type. The per-residue latent representation is **z** *∈* ℝ*^N^*^×*d_z_*^, with *d_z_* = 8. A complete binder is abbreviated by

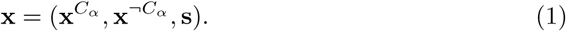

The fixed target context is denoted by **c**^target^. For protein targets it contains Atom37 coordinates, residue identities, chain and residue indices, and optional binary interface-hotspot indicators. For ligand targets it contains atom types and names, coordinates, charges, graph-Laplacian positional encodings, bond masks, and bond orders.

The partially latent construction preserves the spatial backbone explicitly while placing residue identity and all non-C*_α_* geometry in a fixed-dimensional continuous latent space. The joint conditional density of a binder and its latent representation factorizes as

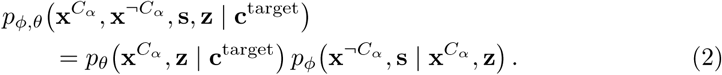

The corresponding conditional density over complete binders is obtained by marginalizing the latent variable:

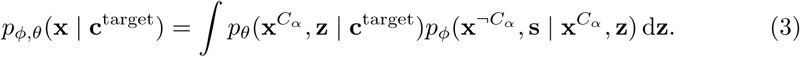

The first factor is generated by a continuous flow, and the second factor is the autoencoder decoder. Thus the main generator does not have to evolve a mixed categorical–continuous state or a residue-dependent number of atoms.

where coordinate-valued quantities are represented in the model’s internal length units.

Complex generation additionally uses a global translation

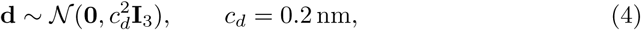

which is independent of the reference variables and is broadcast over all binder residues as **1***_N_* **d**^T^. Separate interpolation times *t_x_, t_z_* ∈ [0, 1] are used for the backbone and latent channels:

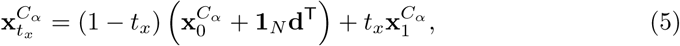

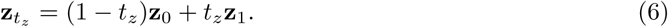

Translation noise is specific to complex generation: it makes absolute binder placement relative to the target a learned part of denoising rather than a nearly fixed low-frequency feature.

The conditional velocity network has two outputs,

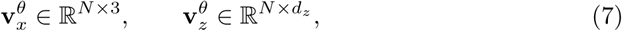

and receives the interpolated binder state, target context, and both times. Because the interpolants are linear, their target velocities are

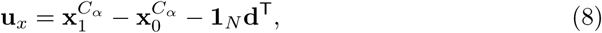

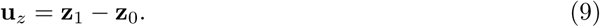

The conditional flow-matching objective is

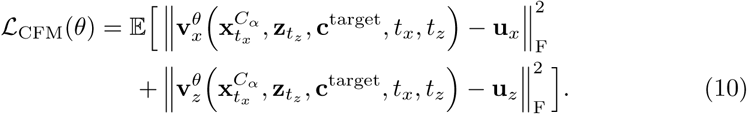

The expectation is over binder–target pairs (**x**, **c**^target^) *∼ p*_data_, encoder samples **z**_1_ *∼ q_ψ_*(**z** | **x**), independent reference noise 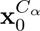 and **z**_0_, translation noise **d**, and independently sampled interpolation times.

Following La-Proteína, define the time densities

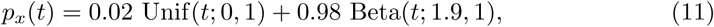

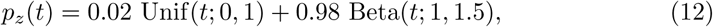

and draw

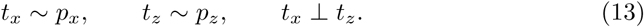

The unequal schedules emphasize different noise regimes for global backbone geometry and local atomistic detail.

### S1.2 Target conditioning and neural architecture

The denoiser operates on a single concatenated sequence containing noisy binder tokens and clean target tokens. Binder token *i* contains its interpolated C*_α_* coordinate 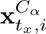, its interpolated latent vector **z***_tz,i_*, and a positional encoding of its sequence index. Protein-target tokens contain amino-acid identity, fixed Atom37 geometry, chain identity, and optional hotspot indicators. Ligand-target tokens instead represent individual target atoms using the element, charge, aromaticity, and other atom-level features defined above. All token types are projected into a common hidden dimension before being processed jointly.

Supplementary Fig. S10a shows the construction of the initial sequence representation. Sinusoidal encodings of sequence indices are linearly projected and added to learned projections of the coordinate and feature channels. The coordinate projections allow the network to distinguish the current noisy binder geometry from the fixed target geometry, while the feature projections distinguish binder latent states, protein-target residues, and ligand-target atoms. A small set of learned register tokens is appended to the physical tokens. These registers provide sequence-level workspace without being associated with a residue, atom, or generated coordinate.

The two interpolation times enter through a separate conditioning pathway (Supplementary Fig. S10b). Sinusoidal embeddings of *t_x_* and *t_z_* are concatenated and transformed by an MLP to obtain a conditioning vector **c***_t_*. This vector is aligned with the binder and target tokens and is used throughout the transformer to modulate normalization and residual updates. The conditioning entries associated with register tokens are set to zero. Keeping *t_x_*and *t_z_*separate allows the network to adapt its computation independently to the noise levels of the backbone and latent channels.

The initial pair representation is constructed as shown in Supplementary Fig. S10c. For protein pairs, its inputs include coordinate-derived pair distances, relative sequence separation, chain identity, and the distinction between binder and target tokens. Ligand pairs additionally contain bond connectivity and bond order, and protein–ligand pairs encode their current cross-interface geometry. The geometric and sequence-derived features are concatenated and adaptively normalized using the time representation. A transformed embedding of *t_x_* and *t_z_* is then concatenated with these features. Rows and columns introduced by the register tokens are zero-padded so that registers participate in sequence attention without introducing artificial geometric relationships.

Each transformer block maintains a sequence representation **H** *∈* ℝ*^L×ds^* and a pair representation **P** *∈* R*^L×L×dp^*, where *L* includes the binder, target, and register tokens. As illustrated in Supplementary Fig. S11a, adaptively normalized sequence states are projected into queries, keys, and values. A normalized linear projection of the pair representation supplies an additive bias to each attention head:

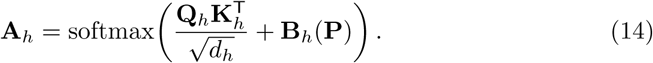

Consequently, attention between two tokens depends both on their sequence representations and on their pairwise geometric or chemical relationship. The attention outputs are concatenated, linearly projected, adaptively scaled, and added to the sequence residual stream. A second adaptively normalized sublayer applies a linear–SwiGLU transition followed by another conditioned residual update.

The pair representation is updated after the sequence attention operation (Supplementary Fig. S11b). Two learned projections of the normalized sequence states are combined elementwise to produce a sequence-derived pair update. This update is added to **P**, after which outgoing and incoming multiplicative triangle layers propagate information over token triplets. A normalized feed-forward pair transition, implemented as linear–ReLU–linear transformations, completes the pair update. Residual connections around these operations preserve the previous pair state while allowing geometric and interaction information to accumulate over successive blocks.

The adaptive operations used in the sequence pathway are detailed in Supplementary Fig. S11c–d. Suppressing token and channel indices, their action can be summarized as

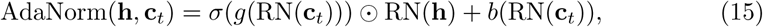

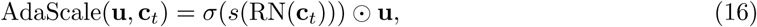

where *g*, *b*, and *s* are learned projections and RN denotes the base normalization operation. The first operator conditions both the scale and shift of a normalized activation, whereas the second controls the magnitude of a residual branch. This makes the computation in every block explicitly dependent on both flow times.

After the final transformer block, only the binder-token representations are passed to two output projections. The coordinate head produces 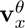, with one three-dimensional velocity per binder residue, and the latent head produces 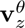, with one latent-space velocity per binder residue. Target and register outputs are discarded. Target coordinates remain fixed during integration and influence generation only through the joint sequence representation, pair features, and attention biases.

Only the conditional flow model receives **c**^target^. The atomistic autoencoder remains a target-independent monomer encoder and decoder, allowing its pretrained sequence–structure representation to be reused unchanged for conditional generation. Training proceeds from broad protein priors to interaction-specific data. The autoencoder is pretrained on AlphaFold Database monomers and fine-tuned on PDB chains. The conditional flow combines AFDB monomers, Teddymer synthetic domain–domain interfaces, and filtered experimental PDB multimers [21–23]. For ligand-conditioned models, filtered PLINDER complexes are added and the protein model is adapted with low-rank updates [48, 51]. Target coordinates are held fixed throughout generation.

### S1.3 Sampling dynamics and decoding

At inference, Gaussian binder states are advanced from (*t_x_, t_z_*) = (0, 0) to (1, 1). For the Gaussian probability paths used here, the learned velocity determines the scores ***ζ****_x_* and ***ζ****_z_*, the gradients of the log intermediate densities. The stochastic sampler may be written

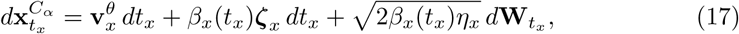

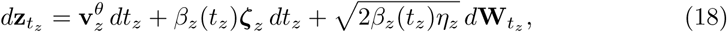

where 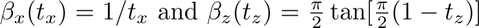. Noise scales *η_x_, η_z_ <* 1, typically 0.1, implement low-temperature sampling. Once the terminal state is reached, the decoder maps 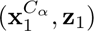 to a categorical sequence and non-C*_α_* Atom37 coordinates. Consequently each trajectory directly returns a sequence–structure pair; no inverse-folding model is required.

### S1.4 Reward-guided inference-time search

Let *p^θ^*(**x c**^target^) denote the terminal distribution of the conditional generator, *R*(**x**) a real-valued reward, and *S*(**x**) a Boolean success criterion. Rewards used here include structure-prediction interface confidence, hydrogen-bond energy and satisfaction, shape complementarity, pocket burial and physicochemical penalties. A reward for an intermediate state is approximated by stochastically rolling that state to *t_x_*= *t_z_*= 1, decoding it, and evaluating the resulting clean complex. This keeps the scoring model outside the generator and permits non-differentiable rewards.

#### Best-of-N

Independent terminal samples are ranked by *R*. This increases inference compute without changing any trajectory and provides the reference case for adaptive methods.

#### Beam search

Let *B_k_* be a beam of *N* partially denoised states at search stage *k*. Each state is advanced for *K* discretized steps along *L* independent stochastic branches, producing *NL* candidates *C_k_*_+1_. With rollout reward *R̂*, the next beam is

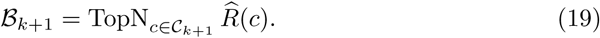

Completed rollout samples satisfying *S* are retained rather than discarded. Unless stated otherwise, *N* = 4, *L* = 4, *K* = 100, and the base trajectory contains 400 discretized denoising steps.

#### Feynman–Kac steering

Feynman–Kac steering targets the exponentially tilted terminal density

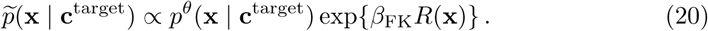

It uses the same branch-and-rollout construction as beam search, but resamples *N* particles from *C_k_*_+1_ with probabilities

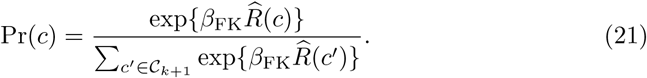

We use *β*_FK_ = 10. Deterministic top-*N* selection concentrates more aggressively, whereas resampling preserves reward-weighted diversity.

#### Monte Carlo tree search

MCTS interprets discretized denoising as a tree whose nodes are partially latent states. Selection, stochastic expansion, terminal rollout and reward backpropagation are repeated. For a child *j* of node *i*, the exploration–exploitation score has the form

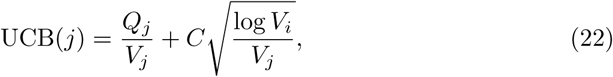

where *Q_j_*is accumulated rollout reward, *V_j_*is the child visit count, *V_i_* is the parent visit count and *C* is the exploration constant. Unlike beam search, MCTS can revisit an earlier branching decision when later rollouts reveal that it is promising.

### S1.5 Codesign-based scaffold re-engineering

For scaffold re-engineering, an observed binder is encoded, then moved to an intermediate noise level (*τ_x_, τ_z_*) using Eqs. (5) and (6). Denoising from that state regenerates both backbone and latent variables conditional on the same target. The pair (*τ_x_, τ_z_*) controls the edit radius: small values retain little of the input, whereas values near one produce local sequence and geometry changes. Because the decoder returns both modalities, this procedure is codesign rather than fixed-backbone sequence redesign. The Nipah campaign applies this diffuse–denoise construction alongside fully de novo sampling.

## S2 Biophysical characterization of designed monomers: experimental methods

### Design filtering

Structure predictions for La-Proteína co-designed and ProteinMPNN-designed sequences for the unconditional design benchmark set were generated in single-sequence mode with 3 recycles using the *alphafold2_ptm_model_1* of AF2 in its ColabFold implementation. Designs were considered successful if the predicted structures had a mean pLDDT *>* 90 and an RMSD *<* 2 Å from the design model. All successful designs were subsequently manually screened; a subset of 49 designs were selected for experimental analysis. La-Proteína co-generated sequences were preferred whenever metrics matched those of the respective Protein-MPNN sequence. Sequence identity and structural similarity were further analysed using HHblits and the Foldseek web server, respectively.

For the high-temperature generation set, designs were predicted both in single-sequence mode and with MSA. Designs were selected for further analysis if they were confidently predicted with MSA (mean pLDDT *>* 80 and RMSD *<* 2 Å from the design model), while being poorly predicted in single-sequence mode (mean pLDDT *<* 80 and RMSD *>* 4 Å). Successful designs were manually screened and filtered based on sequence identity (not higher than 40%), yielding a final set of 13 designs.

### Cloning, expression, purification, and yield quantification

Genes coding for the designed proteins were codon-optimised for *Escherichia coli* K-12 and synthesised by GenScript. Gene fragments were cloned into the LM627 bacterial expression vector using BsaI-mediated Golden Gate assembly, replacing the vector’s *ccdB* selection marker. The assembly products were transformed into *E. coli* NEB5*α* cells (New England Biolabs) through heat-shock transformation and selected on LB agar (Carl Roth) supplemented with kanamycin (VWR). Plasmids were purified from overnight cultures using the QIAprep Spin Miniprep Kit (Qiagen) and verified through Sanger sequencing (Eurofins Genomics). The sequence-verified plasmids were subsequently transformed into *E. coli* BL21(DE3) cells (New England Biolabs) for protein expression.

Proteins were expressed by auto-induction in TBII medium (MP Biomedicals), supplemented with kanamycin (VWR), a 50×5052 autoinduction mixture, 20 mM MgSO_4_, and a trace metal mix. The cultures were initially grown at 37 °C for 8 hours, followed by incubation at 18 °C for 24 hours with shaking. Cells were harvested by centrifugation at 5,000 *g*, resuspended in lysis buffer (50 mM Tris, pH 8.0, 250 mM NaCl, 20 mM imidazole) containing lysozyme and DNase I (Sigma-Aldrich), and lysed by sonication. Clarified lysates obtained by centrifugation were purified using immobilised metal affinity chromatography with HisPur Ni-NTA resin (Thermo Fisher Scientific). Samples were washed with wash buffer (50 mM Tris, pH 8.0, 250 mM NaCl, 20 mM imidazole) and eluted using the same buffer containing 500 mM imidazole. Eluted proteins underwent further purification by size-exclusion chromatography (SEC) on a Superdex 75 Increase 10/300 GL column (Cytiva), using an ÄKTA Pure system (Cytiva) equilibrated in 25 mM Tris, pH 8.0, and 150 mM NaCl.

Protein yields were estimated from the SEC chromatograms recorded at 280 nm by integrating the absorbance peak corresponding to the expected elution of the target species. Peak areas were converted to protein mass using sequence-derived extinction coefficients, corrected for the ÄKTA flow-cell path length, and scaled from the injected sample to the total Ni-NTA eluate. To compare soluble expression yields between different sequence design strategies, a subset of constructs sharing the same backbone but differing in sequence (co-generated by La-Proteína or fixed-backbone design with ProteinMPNN) was expressed and purified in three independent replicates.

### Size-exclusion chromatography coupled to multi-angle light scattering (SEC–MALS)

Where needed, SEC-purified proteins were concentrated prior to further analysis to ensure sufficient scattering signal. Molecular weight was determined by SEC–MALS using an ÄKTA Pure system equipped with a Superdex^TM^ 75 10/300 GL column and a Wyatt miniDAWN^TM^ detector. Extinction coefficients were calculated from the protein sequence and used by the ASTRA software to determine molecular weights.

### Circular dichroism (CD)

Proteins were diluted and/or buffer-exchanged into PBS (100 mM KCl, 10 mM phosphate) buffer to concentrations between 0.1–0.3 mg/mL prior to analysis and measured in a 2 mm Hellma QS cuvette. A JASCO J-1500 spectrometer equipped with a Xe lamp was used in combination with the Spectra Manager^TM^ CFR software. Spectra between 200 and 250 nm were initially obtained for each sample at 25 °C to confirm protein folding. Subsequently, temperature-ramp measurements at three distinct wavelengths (208, 222, and 230 nm) were recorded in steps of 10 °C between 25 °C and 95 °C. Once overall stability was confirmed, full spectra in the range of 200–250 nm were obtained at the start and end points (25 °C, 95 °C, and return to 25 °C). All data were baseline-corrected with PBS.

### Crystallization, data collection, and structure determination

Protein crystals of LP46 (3.9 mg/mL) were obtained by hanging drop vapor diffusion at room temperature, using 1 *µ*L protein in 20 mM TRIS (pH 8.0), 150 mM NaCl buffer, and 1 *µ*L reservoir solution. The protein sample crystallized in 0.1 M NaAcet (pH 4.3), 32.5 % (v/v) ethylene glycol within 3 days. To improve crystal growth, microseeding was applied using protein crystals of the same condition in the subsequent finescreen. The crystal samples were flash-frozen in liquid nitrogen.

The collection of crystallographic data was conducted at the BESSY II Beamline MX14.1 [60]. Diffraction data were processed using xds, and anisotropy corrected and merged using STARANISO [61, 62]. Phases were determined by Phaser molecular replacement, using the designed structure of LP46 as search model [63]. Manual model refinement was performed in Coot (0.9.8) and automated refinement was carried out with PHENIX.refine (v2.1) [64, 65]. Data collection, phasing, and refinement statistics are summarized in Table S7. Data processing software packages were used from the SBGrid software environment, which provides curated, version-controlled structural biology applications and reproducible execution environments across platforms [66].

The crystal structure is shown in Fig. 2f and a detailed per-residue plot of the root mean square deviation between the experimentally determined structure and the computational design generated by La-Proteína is presented in Supplementary Fig. S5.

### Data processing

All data were processed and plotted using Python.

## S3 PDGFR and PD-L1 binder design: filtering and hotspot analysis

### S3.1 Design filtering pipeline

As described in the protein-binder section of the main text (Section 4), a two-stage filtering pipeline was applied to Proteína-Complexa-generated candidates for PDGFR and PD-L1. In the initial screening stage, designs were required to meet the following quantitative thresholds:

*Isoelectric Point ≤* 6.0, selected to ensure favorable solubility and charge profiles under physiological conditions.

*ESM Pseudolikelihood ≤* 10.0, utilized as a metric for sequence fitness and evolutionary plausibility.

*Minimum IPAE ≤* 1.9, ensuring high-confidence relative positioning and orientation of the binder at the target interface.

*Complex iPTM ≥* 0.80, prioritizing designs with a high probability of correct interface assembly.

*Complex pLDDT ≥* 0.88, ensuring high local structural confidence across the designed binder.

*Cysteine Count* exactly 0, to prevent the formation of off-target disulfide bonds and simplify experimental expression and purification.

To ensure broad exploration of the structural landscape, the remaining candidates were clustered at a TM-score threshold of 0.8, with a single representative from each cluster retained for final selection.

### S3.2 Hotspot conditioning analysis

Multiple hotspot combinations were evaluated for both targets to assess the influence of interface residue selection on design success (Table S8). For PDGFR, hotspot sets that included residues A77, A82, and A11—which contribute hydrophobic contacts at the core of the interface—consistently yielded the highest binder rates and affinities. The best-performing combination (NN-PDGFR-6: A139, A77, A82) achieved a 71.1% hit rate with the tightest binder reaching -log(*K*_D_) = *10.03*, corresponding to sub-100 pM affinity. Conversely, the set restricted to peripheral residues (NN-PDGFR-3: A139, A116, A142, A134) yielded a substantially lower hit rate (31.3%) and weaker affinities, confirming that hotspot selection critically shapes design outcomes.

For PD-L1, overall hit rates were lower (5–18%), consistent with the more challenging, flatter interface. Nevertheless, hotspot conditioning had a marked effect: the combination NN-PDL1-4 (A39, A98, A51) achieved the highest hit rate of 17.9%, while the single-residue set NN-PDL1-2 (A39 alone) yielded only 5.3%. This indicates that even on difficult targets, multi-residue hotspot conditioning concentrating on key interface positions substantially improves design success.

## S4 Massive-scale benchmark: extended analysis

### S4.1 Large-scale design campaign

Following [35], Manifold Bio selected 127 design targets to define a diverse and biologically relevant benchmark panel for evaluating Proteína-Complexa alongside recent generative and hallucination-based binder design methods. For targets included in prior in vitro studies (e.g., [67]), we used hotspot definitions reported in the literature. For the remaining targets, where rigorously validated hotspots were unavailable for more than 93% of cases, Manifold Bio proposed approximately 10 single-residue hotspots per target and paired each hotspot with distance-based target crops, yielding more than 1,200 formal design tasks.

For each target, we set out to screen 10,000 sequences. For targets with only one known hotspot, this budget was doubled to maintain comparable search breadth. As described in Section 5, approximately half of the total sequence budget was allocated to a diverse hotspot sweep using Proteína-Complexa. The remaining half was split among baseline comparisons with matched compute budgets, Proteína-Complexa sequence-redesign ablations, and Proteína-Complexa sampling-temperature ablations. For each task, we ran one instance of Proteína-Complexa with standard beam search and AlphaFold2 interface predicted aligned error (ipAE) as the reward, generating approximately 27,000 candidate binders. We filtered this pool to the top 1,000 candidates ranked by pAE. For each hotspot task, we recorded candidates satisfying all of the following criteria: minimum interface pAE *<* 2, complex pLDDT *>* 0.8, ipTM *>* 0.7, and fewer than 5 backbone C*α* clashes with the uncropped target (with a clash defined as any atom pair within 1 Å).

For each target, the hotspot with the largest number of candidates passing these criteria was designated the *top hotspot*. This top-ranked in silico hotspot was used for all baseline experiments shown in Fig. 3c. An analysis of the experimental on-target hit fraction against the in silico hotspot ranking is shown in Supplementary Fig. S7. For targets with only one hotspot, we repeated the search twice and retained the best 1,000 sequences from each run. We did not evaluate samples from hotspots that yielded no in silico success.

In the experiment testing in total 492,636 Proteína-Complexa sequences spanning multiple hotspot combinations for panel-wide target and epitope coverage, to screen a structurally diverse binder set for each target, we pooled all generated binders from hotspots with at least one *in silico* successful design and clustered them at the target level by predicted binder structure. We then filled the evaluation budget using a staged selection procedure that cycled across hotspots and sampled without replacement from these target-level structure clusters. Specifically, we iterated through four ranking regimes in sequence: minimum ipSAE, ipTM, minimum ipAE, and random sampling. Within each regime, the top cluster candidate was selected from previously unselected clusters whenever possible, thereby promoting diversity across both hotspot conditions and predicted binding modes while still prioritizing high-confidence AlphaFold2 predictions (see Supplementary Fig. S6). The resulting screen, including both on-target and off-target hits, is shown in Fig. 3a.

In the experiment testing in total 390,569 sequences from all evaluated method and sequence-design configurations in a compute-matched comparison, we used only the top-ranked in-silico hotspot and we ranked the 1,000 retained Proteína-Complexa candidates by minimum ipSAE and selected the top 400 without clustering. For the baseline methods we similarly chose the top 400 candidates by minimum ipSAE without clustering from the pool of samples generated with the same computational budget across all methods. Hence, the baseline method allocation within the candidate budget for experimental testing was distributed evenly across methods. The same selection procedure was applied to sampling-temperature and sequence-redesign ablations using the top hotspot condition. Minimum ipSAE [58] was chosen here as the primary selection metric to align with recent in vitro practice and to provide a consistent criterion across methods. For sequence-redesign ablations, we followed [18], applying soluble ProteinMPNN for both full-sequence redesign and fixed-interface redesign. For temperature ablations, we started from the default backbone/local latent temperatures of 0.1, 0.1 and evaluated 0.1, 0.4, 0.4, 0.1, and 0.4, 0.4.

Overall, the campaign required more than 140,000 GPU hours and produced a curated set of 1,015,870 binders that were successfully experimentally tested. Because Proteína-Complexa was run across more than 1,200 simultaneous design tasks, we standardized beam search hyperparameters to generate roughly 27,000 binders per task. Under this setup, 80% of tasks finished within 20–32 GPU hours, with higher costs for longer target crops. To match compute across methods, each baseline received approximately 32 GPU hours per target and hotspot. Because downstream experimental processing introduced modest differences in final tested counts, we report tested sequence counts alongside hit counts and hit rates. For the initial designed set on the 83 solvable baseline targets (Fig. 3c), Proteína-Complexa used 26 GPU hours on average, with 80% of targets under 32 GPU hours and a maximum runtime of approximately 40 GPU hours. Additional baseline implementation details are provided in Supplementary S4.6.

### S4.2 Experimental testing: Manifold Bio large-scale all-by-all binder experiment

#### Protein sourcing

Recombinant proteins were obtained from commercial vendors (Supplementary Table S6) and accompanied by vendor quality-control documentation (e.g., SDS-PAGE purity and activity/binding checks). Proteins were reconstituted and stored according to manufacturer specifications before use.

#### Library cloning, phage production, and display

All binder polypeptide sequences were back-translated using Manifold Bio’s internal codon optimization tool and ordered as oligopools (Twist Biosciences). Libraries were cloned into a pADL-100 derivative phagemid (Antibody Design Laboratories) using Gibson Assembly. Insert/vector mixtures were assembled under standard conditions, electroporated into bacterial cells, and assessed for library coverage. Libraries with adequate coverage were prepared and stored at −80 °C. Phage display protocols followed [35]. Sequencing libraries were indexed and run on an Illumina NovaSeq X (Broad Clinical Labs, MA). Paired-end reads (2 × 150 bp) were merged with BBMerge [59]. Mean sequence quality was Q40 (error rate ≈ 10*^−^*^4^ per base). Only exact matches to designed binder variants were counted.

#### Computational binder selection and hit calling

Hits were called from enrichment statistics computed against the pre-selection input library, then classified by the number of distinct screen targets called positive relative to the intended design target. Single-target hits were labeled *on-target specific* (match to design target) or *off-target specific* (single hit against non-design target). Hits spanning two to four targets were labeled *on-target cross-reactive* (design target plus one to three additional targets) or *off-target cross-reactive* (two to four non-design targets). Read counts were converted to frequencies under a Dirichlet–multinomial model with a symmetric prior (*α* = 1), giving posterior-mean frequency *f_i_* = (*n_i_* + *α*)*/*(*N*_reads_ + *Mα*) and posterior standard deviation *σ_i_* = 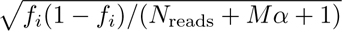 for each of the *M* binders. The prior acts as Laplace smoothing so that no binder has an estimated frequency of exactly zero, which is required for the ratios below. Frequencies were scaled to concentrations, yielding input concentration *c_i_* with error *e_ci_* and, for each of the *k* replicate wells *r* of target group *g*, output concentration *c_o,i,g,r_*. For binder *i* and target group *g* the enrichment ratio and its propagated error are

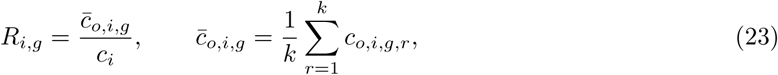

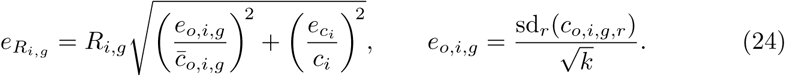

Normalizing to the pre-selection input library is what distinguishes this statistic from a rank threshold on output signal alone: a binder abundant in the output because it was already abundant in the input is not enriched, and a low-input binder that survives selection is recovered even though its absolute output signal is small.

Because the great majority of library members are not selected, the null enrichment level is estimated robustly from the data itself,

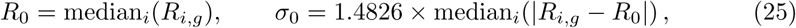

which is insensitive to contamination by up to 50% true hits. A single global (*R*_0_*, σ*_0_) per sub-library is not adequate, however: *R_i,g_* = *c̄_o,i,g_/c_i_* is inflated by small denominators, so low-abundance binders are systematically over-called against a null dominated by high-abundance ones. We therefore stratify the null by input abundance. Binders are sorted by log_10_ *c_i_* and divided into *B* equal-count quantile bins (*B* = min(50, ⌊*n/*100⌋), at least 3); 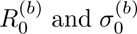 are computed per bin by Eq. 25 and linearly interpolated to give per-binder *R*_0*,i*_ and *σ*_0*,i*_, with *σ*_0*,i*_ floored at 10% of the global value for numerical stability. The null is estimated separately within each sub-library, so sub-libraries of differing quality do not borrow each other’s background. The test statistic is the studentized enrichment

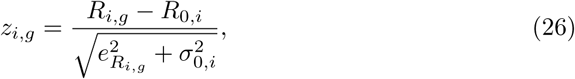

which adds replicate measurement error and null spread in quadrature, so a noisily measured binder must deviate further to reach the same significance. Only right-tail deviations are tested, since depletion below the null is counter-selection rather than binding, and in an earlier two-sided formulation every hit drawn from the depleted tail was off-target:

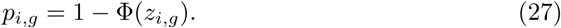

Magnitude is reported separately as FC*_i,g_* = log_2_(*R_i,g_/R*_0*,i*_).

A binder was called a hit for target group *g* when *p_i,g_ <* 0.05 and FC*_i,g_ >* 2, and it passed exactly one target gene across the panel. The magnitude term is required alongside significance because with *M ∼* 10^5^ binders per group a small but highly reproducible ratio can be statistically significant without being biologically meaningful. Because the uniqueness term depends on how many of the 129 targets a binder passed, it is a panel-global property and is evaluated in a single pass over the full panel rather than reconstructed from any one target’s statistics.

Called hits were assessed against the negative-control (“empty”) wells carried on every plate, which are real outputs of the same selection, amplification and sequencing process with no target protein present and therefore no target-specific enrichment. For each hit we compare its mean output across the target’s *k* replicate wells to its mean across all control wells. Over the 40 highest-yielding target groups of a representative screen (18,536 calls), hits sit a median 4.2× above their own control-well level (inter-quartile range 3.8–4.8×; 2.0–18.2× across targets, every target at ≥2×), and for 39 of 40 targets fewer than 4% of calls have a control-well mean that reaches their target mean (median 0.7%). Applying the identical statistic to *k* control wells drawn as a decoy pseudo-target, on the same binders, yields a median ratio of 0.79× — so the separation is specific to the presence of the target and is not a property of the statistic or of the binders themselves.

Specificity across the panel is the second line of evidence, and it is built into the hit rule rather than applied afterwards: a binder enriched on more than one target gene is not called for any of them. Because the 129 targets are screened simultaneously on the same library, each target’s panel-mates act as mutual controls, so a binder that clears the significance and magnitude terms broadly is recognised as non-specific rather than counted once per target.

### S4.3 Per-target hotspot analysis

To characterize hotspot-conditioning effects on design success, we analyzed hit distributions across hotspot combinations for targets with multiple tested hotspots (Supplementary Fig. S7). For each target, cells report the fraction of designs yielding on-target specific hits for a given hotspot, with in silico hotspot rank annotated. Targets are ordered by hotspot entropy, which quantifies how evenly success is distributed across hotspot choices. Targets at the top are robust to hotspot choice, whereas targets at the bottom are hotspot-sensitive. Notably, the top-ranked computational hotspot is not always experimentally optimal, supporting multi-hotspot evaluation per target.

### S4.4 Model temperature ablation

To assess sensitivity to sampling temperature, we ran an ablation across four backbone/local latent temperature pairs on the 83 solvable baseline targets, designing against each target’s top-ranked in silico hotspot. The default setting (0.1, 0.1) yielded the highest on-target hit count (702) and specificity (86.5%), while solving 35 targets (see Supplementary Table S1). Increasing local latent temperature to 0.4 while keeping backbone temperature at 0.1 solved 34 targets and produced fewer total hits (495), although the solved targets were different ones (new/lost columns in table). Each temperature setting uniquely solved 3–6 targets not solved by the others. Specificity remained stable (73.0–86.5%), suggesting elevated temperatures did not materially reduce selectivity. These results support a mixed-temperature strategy to expand solvable-target coverage while preserving hit quality.

### S4.5 Compositional dropout from the phage input pool

Overall, approximately 1.4 million minibinder sequences were designed and ordered across four length-stratified pools. Deep sequencing of the phage input pool before selection recovered 73.2% of designs at one or more reads, resulting in the 1,015,870 successfully tested binders. Two sequence features strongly predicted dropout: sequence length and fractional lysine+glutamate (K+E) content.

Across amino acids, lysine and glutamate showed the strongest negative Spearman correlation with detection (K: *ρ* = —0.268, E: *ρ* = −0.217; Supplementary Fig. S12b). Joint binning by length and K+E fraction (Supplementary Fig. S12a) showed detection rates from 97.9% at 50 aa with K+E *<* 0.08 to 0.19% at 88 aa with K+E *>* 0.50.

We tested three possible explanations. (i) Net charge alone does not explain the trend: arginine (+1, *ρ* = +0.090) and aspartate (—1, *ρ* = +0.063) were positively correlated with detection. (ii) Helix propensity alone does not explain the trend: alanine, methionine, and leucine have comparable/high *P_α_* values without similarly negative correlations (*ρ* = +0.071, —0.038, —0.050). (iii) Adenine-rich codons alone do not explain the trend: asparagine and glutamine share related codon motifs yet show no corresponding negative behavior (*ρ* = +0.011, +0.145). No single property recapitulated the K+E dropout pattern.

In a PaCMAP embedding of all ≈1.4 million designs in 20-dimensional amino-acid composition space (Supplementary Fig. S12c,d), undetected designs concentrated in the high-K+E region. Among the 1,015,870 detected designs, input abundance spanned 0.011× to 12.3× the median (2nd–98th percentile; 1092-fold), with lower abundance in the same high-K+E region.

An enrichment of K+E among non-recovered de novo designs was independently observed in a CAR-T proliferation assay [68], where combined helical-region K+E content was the strongest predictor of non-recovery (ROC-AUC = 0.91). This suggests a shared compositional liability across both bacterial phage display and mammalian surface-expression contexts.

### S4.6 Details on use of prior methods for benchmarking

#### BindCraft

We used the code and checkpoints provided in the public BindCraft repository. We followed the default default_4stage_multimer_hardtarget configuration for setting and filters. For each target, we used the top-ranked hotspot for the hotspot conditioning of BindCraft. To ensure fair comparison under the constraint of 32 GPU hour computation budget while maintaining a minimum number of samples for meaningful statistical analysis, we adopted the following procedure of sample selection. First, we generated *M* binders with a 32 GPU hour budget. Second, we re-designed and evaluated the generated binder sequence 8 times using soluble weights of ProteinMPNN [36] and held interface residues fixed. We re-designed 8 times to be consistent with the re-designing protocol of all models benchmarked in this paper. Then we randomly trim the generated binder set size to *N* (*N ≤ M*) such that the total generation+redesign+evaluation pipeline could fit in the computation budget. After that, we selected the top-1 re-designed sequence from each generated binder based on the best *minimum ipSAE* [58] metric. If *N <* 400 for large targets that required more compute, we backfilled to 400 samples using the best top-2 (or even top-3 if need) re-designed sequences. In this way, we obtained at least 400 samples per target in the final BindCraft sample set.

#### RFDiffusion

We utilized the open-source implementation of RFdiffusion in the public RFdiffusion repository, and followed the protein-protein interaction (PPI) design settings provided in the official documentation. For each target, we first generated backbone structures and subsequently employed the soluble weights of ProteinMPNN to generate 8 binder sequences per backbone. We performed 8 redesign cycles to maintain consistency with the benchmarking protocol used for all models in this study. A total computational budget of 32 GPU-hours was allocated for the full pipeline, including backbone generation, sequence redesign, and AlphaFold2-based evaluation. From the resulting pool, we selected the top-1 redesigned sequence for each generated binder based on the best *minimum ipSAE* metric, yielding a final set of 400 binders per target.

#### RFDiffusion3

We adopted the protein binder design protocol following the production settings in the RosettaCommons Foundry repository for RFdiffusion3. The procedure involved an initial generation of binder structures, followed by the application of soluble ProteinMPNN to produce 8 redesigned sequences per structure. We evaluated both the self-generated sequences and the 8 redesigned variants. Under the 32 GPU-hour budget, we selected the top 400 self-generated sequences and the top 400 redesigned sequences based on the *minimum ipSAE* metric. This procedure resulted in a total of 800 binders per target.

#### BoltzGen

We employed the BoltzGen protein-anything protocol to design binders using the code from the public BoltzGen repository. This unified all-atom diffusion approach integrates structure generation (BoltzGen core) and inverse folding (BoltzIF). For each design trajectory, we generated one sequence via Boltz inverse folding and an additional 8 sequences using the soluble weights of ProteinMPNN. We evaluated the self-generated sequences, the Boltz inverse-designed sequences, and the 8 ProteinMPNN-redesigned sequences. The 32 GPU-hour budget encompassed the entire pipeline: self-generation, Boltz inverse design, ProteinMPNN redesign, and the subsequent evaluation of all resulting sequences. Following the standard selection rules based on *minimum ipSAE*, we identified the top 400 sequences from each of the three categories (self-generated, Boltz inverse-designed, and soluble inverse-designed), resulting in a total of 1,200 binders per target.

## S5 ActRIIA binder design: experimental methods

### Expression and purification protocol

Plasmids encoding the top 200 designs were transformed into *E. coli* BL21(DE3) cells. A single colony was inoculated into 100 mL LB medium supplemented with 100 *µ*g/mL ampicillin and cultured at 37 °C until the optical density at 600 nm (OD_600_) reached approximately 0.6 (3–4 hours).

The temperature was then reduced to 18 °C, and protein expression was induced with 0.1 mM IPTG. Cultures were incubated at 18 °C for an additional 16 hours. The overall timeline from gene synthesis to purified protein was approximately three weeks.

To obtain high-quality protein samples for downstream analysis, affinity tags were removed for selected constructs. Cell pellets were resuspended in lysis buffer (1×PBS, pH 7.4), lysed by sonication, and centrifuged to remove cell debris. The clarified supernatant was loaded onto a nickel-affinity column for purification. Eluted fractions were treated with TEV protease to remove the affinity tag and subsequently passed through a second nickel column to separate cleaved protein from uncleaved protein and the His-tagged TEV protease. The target protein was then buffer-exchanged into PBS, and protein concentrations were determined prior to surface plasmon resonance (SPR) analysis. A summary of expression yields for both tagged and tagless constructs is provided in Supplementary Table S9; of the 200 designs, 192 expressed solubly and 110 achieved yields exceeding 50 mg/L.

### SPR protocol

The surface plasmon resonance (SPR) experiments were conducted on a Biacore 8K+ instrument at 25 °C (both sample and analysis temperature) to characterize the binding kinetics of 80 analytes and a Bimagrumab control to the ligand ActRIIA(20–135)-Avi. The ligand was immobilized on all eight channels of a Streptavidin (SA) sensor chip at a concentration of 0.5 *µ*g/mL with a flow rate of 5 *µ*L/min for 52 seconds. Assays were performed using a running buffer of 10 mM HEPES pH 7.4, 150 mM NaCl, 3 mM EDTA, 0.05% Tween-20, and 2% DMSO. The experimental workflow included 15 start-up cycles to stabilize the surface prior to analysis, and data were collected at a flow rate of 30 *µ*L/min using multi-cycle kinetics (MCK) with association/dissociation times of 90/180 seconds. Two independent SPR runs were performed; binding measurements were largely consistent across both experiments. The only design showing noticeable variability was #51, which exhibited fluctuations in both *K*_D_ and *R*_max_ values, possibly arising from nonspecific surface interactions or partial self-aggregation during the assay. Representative sensorgrams from both runs are shown in Supplementary Fig. S13, and the full Activin A functional dose-response panel across all eight lead designs is shown in Supplementary Fig. S14.

### Smad2/3 luciferase reporter assay protocol

A Smad2/3-responsive luciferase reporter assay was performed using a stable HEK293 TGF-*β*/Activin/Myostatin-responsive reporter cell line in a 384-well format to evaluate the inhibitory activity of candidate binders. Reporter cells were seeded at 10,000 cells per well in 20 *µ*L of DMEM-based culture medium in white, clear-bottom 384-well plates and incubated at 37 °C with 5% CO_2_ for approximately 18 h to allow cell attachment. Test proteins, including engineered mini-binder candidates and the reference antibody bimagrumab, were added in 5 *µ*L volumes and pre-incubated with the cells for 4 h. Subsequently, 5 *µ*L of ligand solution was added to stimulate the pathway, using either human Activin A (20 ng/mL final concentration) or GDF8/myostatin (200 ng/mL final concentration). Test molecules were evaluated in three-fold serial dilutions across the plate. Control wells included ligand-stimulated cells (assay medium + ligand) and unstimulated cells (assay medium only) to define maximal activation and basal signaling levels, respectively. After overnight incubation (*∼*18 h) to allow transcriptional activation of the Smad2/3-responsive reporter, 30 *µ*L of luciferase detection reagent was added to each well, and luminescence was measured using a plate luminometer. Inhibition of pathway signaling by test proteins was quantified as the reduction in luminescence relative to ligand-stimulated control wells.

## S6 PAK1 and CK1*δ* kinase binder design: experimental methods

### S6.1 De novo binder design

Candidate binders targeting the PAK1 catalytic domain (PAK1c; UniProt Q13153, amino acids 273–498) or the CK1*δ* catalytic domain (CK1*δ*c; UniProt P48730, amino acids 9–277) were designed using Proteína-Complexa. For the split-NeoR complementation assay, a library of 50 PAK1c binders (49–74 amino acids) was generated. Binder open reading frames were flanked by constant sequences for downstream PCR amplification and cloned into the split-NeoR selection vector. For CK1*δ*c, 18 short peptide binders (*<*31 amino acids) were designed and synthesized as biotin-conjugated peptides.

### S6.2 Split-NeoR bacterial selection

The split-neomycin resistance (split-NeoR) protein-fragment complementation assay was used to select PAK1 binders based on reconstitution of aminoglycoside phosphotransferase (APH(3*^′^*)-II) activity [42, 43]. Binder candidates were fused to the C-terminal fragment of NeoR, and PAK1c was fused to the N-terminal fragment; both fusion proteins were co-expressed in *E. coli* from a single plasmid. The pooled binder library was transformed into NEB Stable Competent *E. coli* (New England Biolabs, C3040) by chemical transformation. Transformants were plated on LB agar without kanamycin (input control) or LB agar supplemented with 25 *µ*g/mL kanamycin (selection) and incubated overnight. Colonies from both conditions were pooled and plasmid DNA was isolated by miniprep for downstream NGS analysis.

### S6.3 Next-generation sequencing and enrichment analysis

Binder-encoding amplicons were prepared by PCR using KAPA HiFi 2× ReadyMix (Roche, KK2601) with library-specific primers flanking the binder open reading frames. PCR conditions were 98 °C for 3 min; 15 cycles of 98 °C for 20 s, 65 °C for 15 s, and 72 °C for 30 s; and 72 °C for 1 min. Template DNA was normalized to 3.0 ng/*µ*L (30 ng input per reaction). Amplicon libraries were submitted for paired-end sequencing (Azenta AmpliconEZ). Reads were mapped to the reference library using a 30 bp ORF-prefix matching strategy anchored to the 5*^′^* constant sequence (5*^′^*-CACTCAGGGTCCGGT-3*^′^*), achieving mapping rates of 91.7–94.7% across samples. Raw read counts were normalized to counts per million (CPM). Enrichment was calculated as log_2_ fold change:

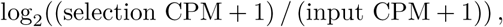

where a pseudocount of 1 CPM was added to avoid division by zero. To classify binders as enriched or depleted, a two-component Gaussian mixture model (GMM) was fitted to the log_2_ fold-change values using expectation-maximization. Two components were chosen based on the observed bimodal distribution. Each binder was assigned to the component with the higher posterior probability, and the decision boundary was defined as the log_2_ fold-change value at which the posterior probabilities of the two components were equal.

### S6.4 Structure prediction

Predicted structures of binder–PAK1 complexes were generated using AlphaFold2 Multimer (v2.3.2) with five models and five seeds per model (25 predictions total per complex). The combined interface predicted TM-score and predicted TM-score (ipTM+pTM) from the best-ranked model was used as the metric of predicted binding confidence.

### S6.5 GST-tagged protein purification

GST-tagged CK1*δ*c was expressed from pGEX-GST-CK1*δ* as an N-terminal fusion in BL21(DE3) *E. coli*, induced with 0.1 mM IPTG, and grown at 20 °C for two nights. Cells were lysed by sonication in lysis buffer (50 mM Tris pH 7.5, 100 mM NaCl, 1 mM EDTA, 1 mM EGTA, 2% IGEPAL CA-630, 3 mM DTT, and protease inhibitors) and clarified by centrifugation and 0.45 *µ*m filtration. Proteins were purified using glutathione-Sepharose resin (Glutathione Sepharose^TM^ 4B, Cytiva) at 4 °C and eluted in 50 mM Tris pH 7.5, 150 mM NaCl, 3 mM DTT, and 25 mM reduced glutathione, as previously described [69]. Protein-containing fractions were identified by Bradford assay, pooled, concentrated by ultrafiltration with PBS washes, and stored at *−*80 °C.

### S6.6 Biotinylated peptide synthesis and streptavidin bead pull-down

Biotinylated peptides were synthesized on a MultiPep 1 automated peptide synthesizer (CEM Corporation) using standard Fmoc solid-phase chemistry. Biotin was incorporated at the N-terminus via a biotinylated lysine followed by two glycine spacers. For pull-down assays, all steps were carried out at 4 °C unless otherwise noted. Streptavidin magnetic beads (Pierce^TM^ Streptavidin Magnetic Beads) were washed three times with 500 *µ*L Bead Wash Buffer (BWB; 25 mM HEPES pH 7.4, 150 mM NaCl, 0.01% Tween-20, and 1 mM DTT). For peptide capture, 17 *µ*L of bead slurry was incubated with 1 mL of 3 *µ*M biotinylated peptide with end-over-end mixing for 30–45 min at 4 °C, then washed three times for 3 min with 500 *µ*L BWB. Purified GST-tagged kinase domain (0.75 *µ*g) was added to peptide-loaded beads in 1 mL BWB+ (BWB supplemented with protease inhibitor) and incubated for 30–40 min at 4 °C with end-over-end rotation. Beads were washed four times with 500 *µ*L BWB+ (10 min each). Bound proteins were eluted in 20 *µ*L of 2× Laemmli Sample Buffer (Bio-Rad) containing *β*-mercaptoethanol at 95 °C for 5 min.

### S6.7 Co-immunoprecipitation

HEK293T cells were seeded in 12-well plates and transfected 24 hours later using PEI MAX 40000 (Polyethylenimine HCl MAX, Linear, Mw 40,000; Kyfora Bio, 24765-100). Each well received 500 ng of binder plasmid (pDS261: pCAG-GFP-3×Flag-P2A-3 HA-binder) and 500 ng of bait plasmid (pDS217: pCAG-PAK1-Myc), with PEI MAX at 20 *µ*L per condition. DNA–PEI MAX complexes were formed in Opti-MEM for 15 min at room temperature and added to cells. Medium was replaced 6 hours post-transfection; cells were harvested 48 hours post-transfection. Cells were washed once with cold PBS and lysed in 300 *µ*L of lysis buffer (20 mM HEPES pH 7.4, 150 mM NaCl, 0.5% NP-40, and protease inhibitor cocktail) on ice for 5–10 min. Lysates were clarified by centrifugation (15,000 × *g*, 10 min, 4 °C), and 5–10% of clarified lysate was saved as input. HA-tagged proteins were immunoprecipitated using 15 *µ*L ChromoTek HA-Trap Magnetic Agarose (Proteintech, cat. no. atma) with rotation for 90 min at 4 °C. Beads were washed three times with 500 *µ*L wash buffer (20 mM HEPES pH 7.4, 150 mM NaCl, and 0.1% NP-40). Bound proteins were eluted in 30 *µ*L 2× Laemmli sample buffer at 95 °C for 5 min. An empty pDS261 vector served as the negative control. Three independent biological replicates were performed.

### S6.8 Western blotting

For PAK1 co-IP experiments, input and eluate samples were resolved by SDS-PAGE on Mini-PROTEAN precast gels (4–20%; Bio-Rad, 4561096) and transferred to membranes. Membranes were probed with anti-Myc chicken polyclonal antibody (Invitrogen, A-21281) to detect Myc-PAK1, followed by IRDye 800CW donkey anti-chicken IgY secondary antibody. Anti-Flag antibody was used to detect GFP-Flag binder expression (IRDye 680RD secondary, 700 nm channel). Blots were imaged using a LI-COR Odyssey system with simultaneous dual-color detection (700 and 800 nm channels). For CK1*δ* pull-down experiments, proteins were transferred to membrane and probed with anti-GST antibody (Bethyl Laboratories GST Tag Polyclonal Antibody, HRP, 1:1000). Standard western blot procedures were followed for blocking, primary antibody incubation, and chemiluminescent detection (SuperSignal^TM^ West Femto Maximum Sensitivity Substrate).

### S6.9 Microscale thermophoresis binding assays

Purified CK1*δ* kinase domain was fluorescently labeled in PBS (pH 7.4) using the RED-NHS 2nd Generation Protein Labeling Kit (NanoTemper Technologies, cat. no. MO-L011) according to the manufacturer’s protocol. Microscale thermophoresis (MST) binding experiments were performed on a Monolith X instrument (NanoTemper Technologies) equipped with a Nano RED detector. Labeled CK1*δ* was diluted to 80 nM in assay buffer (0.14% Pluronic F-127 in PBS, pH 7.4). Peptide binders were prepared as 16-point, twofold serial dilution series in assay buffer, with top concentrations of 100 *µ*M for CK3 and CK16 and 200 *µ*M for CK9. Each peptide dilution was mixed 1:1 (v/v) with labeled CK1*δ* and incubated for 15–30 min at room temperature, protected from light, before loading.

Samples were loaded into Monolith Premium Capillaries (NanoTemper Technologies, cat. no. MO-K025) and analyzed at 25 °C using the 670 nm detection channel and a 1.5 s retention time. Binding was quantified using normalized fluorescence (*F*_norm_), defined as the ratio of fluorescence intensity in the heated region after IR-laser activation to the pre-heating baseline fluorescence (López-Méndez et al., 2021). Averaged *F*_norm_ values were plotted against log_10_([ligand]) and fitted by nonlinear least-squares regression using the Levenberg–Marquardt algorithm to a three-parameter logistic model with the Hill slope fixed at 1. Dissociation-constant (*K*_d_) measurements were performed in triplicate for each binder. Capillaries with fluorescence counts outside the 500–1,400 range or with excessive aggregation were excluded before curve fitting.

### S6.10 Image quantification and statistical analysis

Band intensities were quantified using Fiji/ImageJ. For PAK1 co-IP experiments, co-IP efficiency was calculated as the ratio of anti-Myc signal in the eluate to the anti-Myc signal in the input (elution/input ratio), then normalized to the empty-vector control within each replicate to obtain fold-enrichment values. Statistical significance was assessed by one-sample *t*-test (two-tailed) against a null hypothesis of fold enrichment = 1.0. Data are presented as mean ± SD from three independent biological replicates. For CK1*δ* pull-down experiments, samples were normalized to the scrambled control for their respective run. Statistical significance was assessed by one-sample *t*-test (one-tailed) against a null hypothesis of fold enrichment = 1.0. Data are presented as mean *±* SD from three to four independent biological replicates.

## S7 Carbohydrate binder design: experimental methods

### S7.1 Design generation and filtering

Candidate binders targeting the type II A pentasaccharide were generated using Proteína-Complexa conditioned on the sugar ligand structure. To promote precise hydrogen-bonding networks and deep pocket burial—both critical for overcoming the large desolvation penalty of carbohydrate binding—inference-time rewards included pocket burial scores and hydrogen-bond counts at the binder–ligand interface. Twenty-four candidates were selected for expression testing; 23 designs, all smaller than 10 kDa, advanced to the binding screen reported in the main text (Supplementary Fig. S15).

### S7.2 Protein cloning and expression

The 24 initial candidates were gene-optimized for *E. coli* expression and ordered as gene strings with 20 bp overhangs compatible with the acceptor vector T7-pRSF. The T7-pRSF vector was linearized by PCR, and residual circular template was removed by DpnI (NEB) digestion. Linear product was purified by SPRI beads (Beckman) and eluted at a concentration of 50 ng/*µ*L. Genes were ligated into the vector using Gibson assembly and transformed into BL21(DE3) cells (NEB) under kanamycin selection. Colonies were verified by whole-plasmid sequencing.

Cells were grown at 37 °C in 5 mL LB-kanamycin medium to an OD_600_ of 0.4–0.8, at which point expression was induced with 1 mM IPTG. Cultures were transferred to 20 °C with agitation at 1,000 rpm on a flat shaker-incubator, and expression was carried out for 14 hours. Cells were pelleted by centrifugation (5 min, 4,000 × *g*) and lysed in 2 mL lysis buffer (10× BugBuster (Millipore), diluted in PBS supplemented with CaCl_2_ and MgCl_2_ (Thermo)). Lysates were agitated at 1,000 rpm for 30 min at 20 °C and clarified by centrifugation (10 min, 4,000×*g*).

The supernatant was collected and incubated with Ni-NTA agarose beads (200 *µ*L slurry per protein) with agitation at 1,000 rpm for 30 min at 20 °C. After centrifugation (3 min, 300 × *g*), the supernatant was discarded and the bead pellet washed with 1 mL PBS. Proteins were eluted in 200 *µ*L elution buffer (PBS, 250 mM imidazole) with agitation for 10 min. The eluate was subjected to two rounds of buffer exchange into PBS using 3 kDa size-exclusion filters (Amicon), with 10-fold dilution and re-concentration per round. Final protein concentrations, quantified by A_280_ (NanoDrop), ranged from 0.8 to 3.5 mg/mL. Proteins were stored at 4 °C and assayed within three days of purification.

### S7.3 Red blood cell agglutination assay

Commercially sourced reagent red blood cells (types A_2_ and B) were tested separately. Each neat stock was diluted 1:100 in PBS and washed once by centrifugation (3 min, 300×*g*) without further dilution. A Ni-NTA agarose bead suspension was prepared at a ratio of 2:48 (neat resin:PBS, v/v). For each sample and replicate, 50 *µ*L of bead suspension was loaded with 2 *µ*L of purified binder solution, and 50 *µ*L of the relevant RBC suspension was added. Negative controls included RBCs + beads (no protein), RBCs + protein (no beads), and beads + protein (no RBCs). Samples were incubated in U-bottom 96-deep-well plates (Greiner), with a separate plate per replicate. After 10 min, samples were inspected macroscopically and microscopically for visible agglutination. After 2 hours of incubation, samples were imaged using an EVOS-FL microscope at the region of highest bead density per well, then agitated and OD_600_ measured using a plate reader (SpectraMax). Data are plotted as mean ± standard deviation of three independent replicates. A construct containing the B-binding CBM51 domain served as the positive control, and RBCs + beads without protein served as the negative control baseline.

### S7.4 Glycan–DNA conjugation for BLI

To produce a glycan conjugate of sufficient molecular weight to elicit a detectable biolayer interferometry (BLI) response, the blood group A pentasaccharide (type II; Biosynth) was conjugated to a modified DNA oligonucleotide. An oligonucleotide bearing three amine-modified positions ([AmC6]GA[AmC6dT]GA[AmC6dT]CTC-GACGCTCTCCCTTATGCGACTCC; Integrated DNA Technologies) was mixed with the pentasaccharide at a 1:5 molar ratio in 50% DMSO containing 100 mM 2-picoline borane (Sigma). The reductive amination reaction was incubated for 3 hours at 37 °C. The product was purified and buffer-exchanged into PBS using PD-10 columns (Sephadex G-25, 3 mL elution). Recovered DNA yield was estimated by A_260_ (NanoDrop).

### S7.5 Biolayer interferometry

BLI experiments were performed on an Octet K2 (Sartorius) two-channel system using HIS1K biosensors (Sartorius, 18-5120). Interactions were measured in flat black 96-well plates (Nunc; 200 *µ*L/well) with shaking at 1,000 rpm. The assay sequence consisted of: baseline (PBS, 120 s), loading (protein in PBS at 0.1 mg/mL, 120 s), baseline (PBS, 120 s), association (four sequential steps of 120 s each at glycan–DNA conjugate concentrations of 2, 4, 6, and 8 *µ*M in PBS), and dissociation (PBS, 120 s). A reference sensor was treated identically but without protein in the loading step. Data were acquired at 5 Hz and reference-subtracted.

### S7.6 Circular dichroism

CD spectroscopy was performed on an Aviv 410 CD spectrometer using a quartz cuvette (1 mm path length). Design NV15 (S15) was loaded at a final concentration of 0.25 mg/mL in PBS. A wavelength scan (200–250 nm) was carried out at 25 °C with a bandwidth of 1.00 nm to confirm the presence of secondary structure elements. For thermal stability analysis, CD signal was monitored at 222 nm with a bandwidth of 10 nm. Dynode voltage was recorded concurrently to check for aggregation, which was not observed. An initial scan from 40 to 95 °C identified the inflection region, followed by a higher-resolution scan from 70 to 98 °C. Secondary structure elements remained stable beyond 100 °C, precluding accurate determination of a melting temperature within the accessible range.

## S8 Supplementary figures

The following figures supplement the main text and Supplementary Methods.

**Fig. S1.**
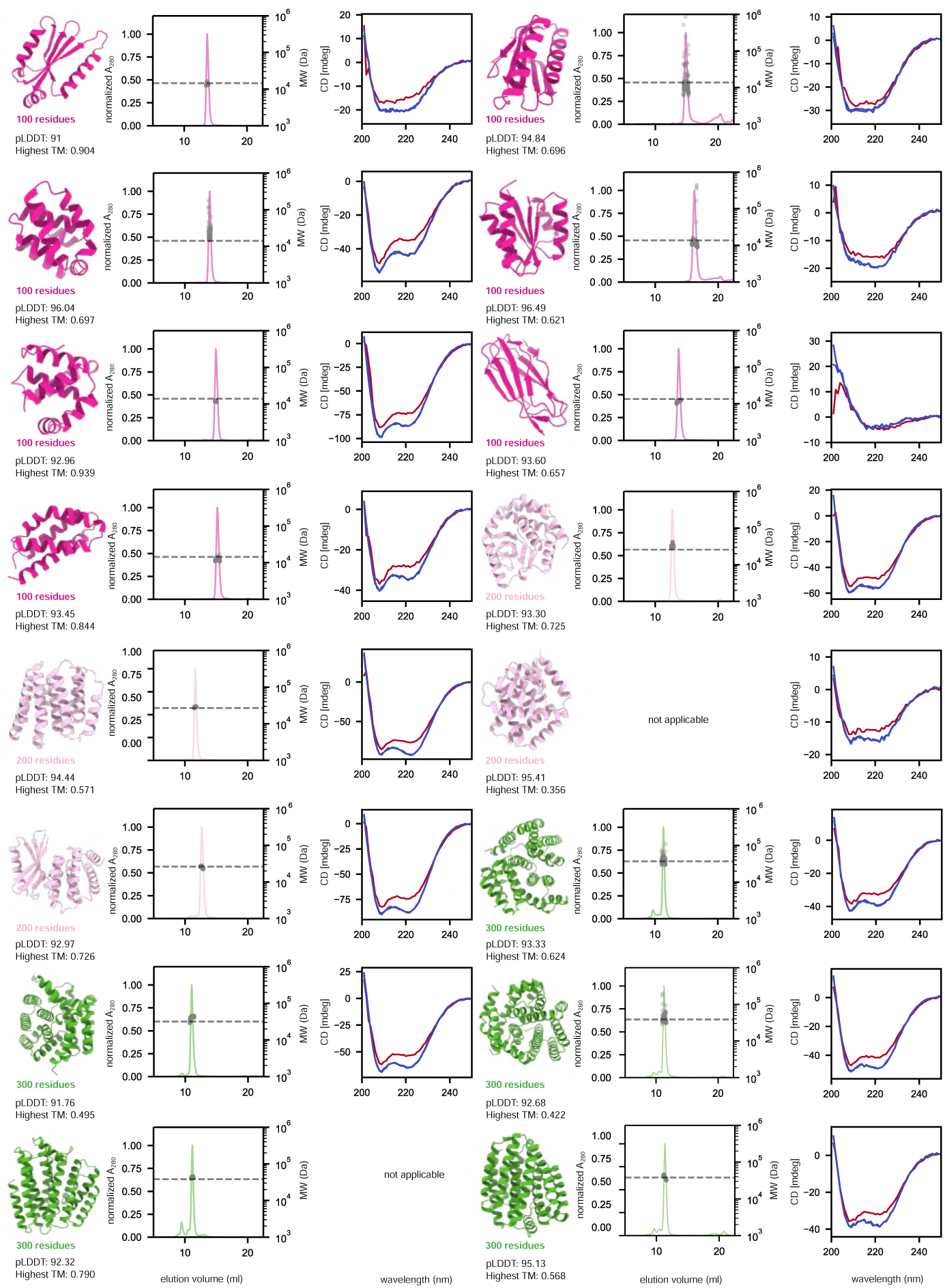
Biophysical characterization of designed monomeric folds: 100–300 residue designs. For each design, the predicted structure model (left), SEC profile overlaid with MALS-determined molecular weight (middle), and CD spectra at 20 °C (blue) and 95 °C (red) (right) are shown. Protein models are colored according to chain length. The average pLDDT from AF2 single-sequence predictions and the Foldseek TM-score to the closest known fold are reported for each design.

**Fig. S2.**
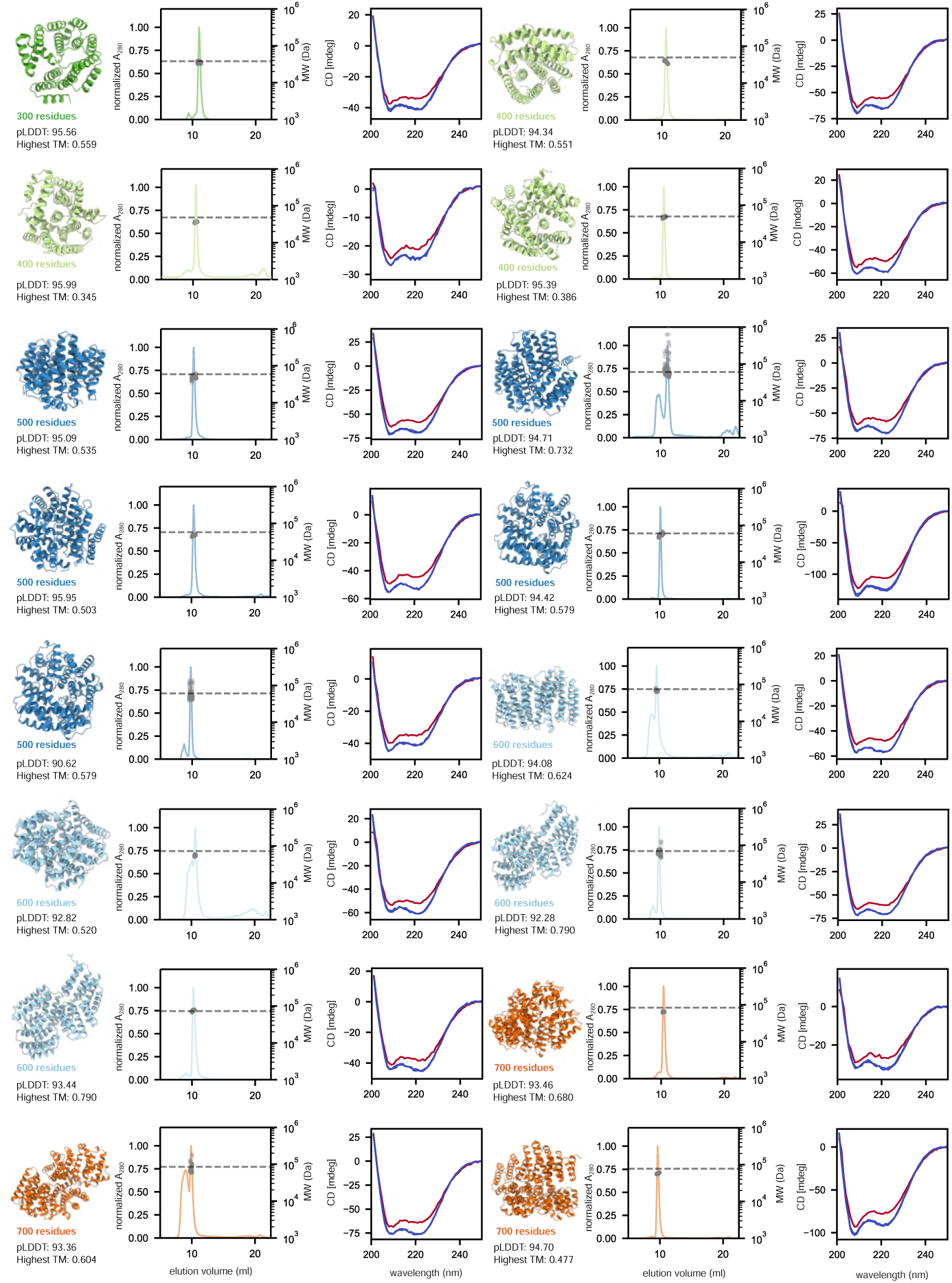
Biophysical characterization of designed monomeric folds: 300–700 residue designs. Layout as in Supplementary Fig. S1.

**Fig. S3.**
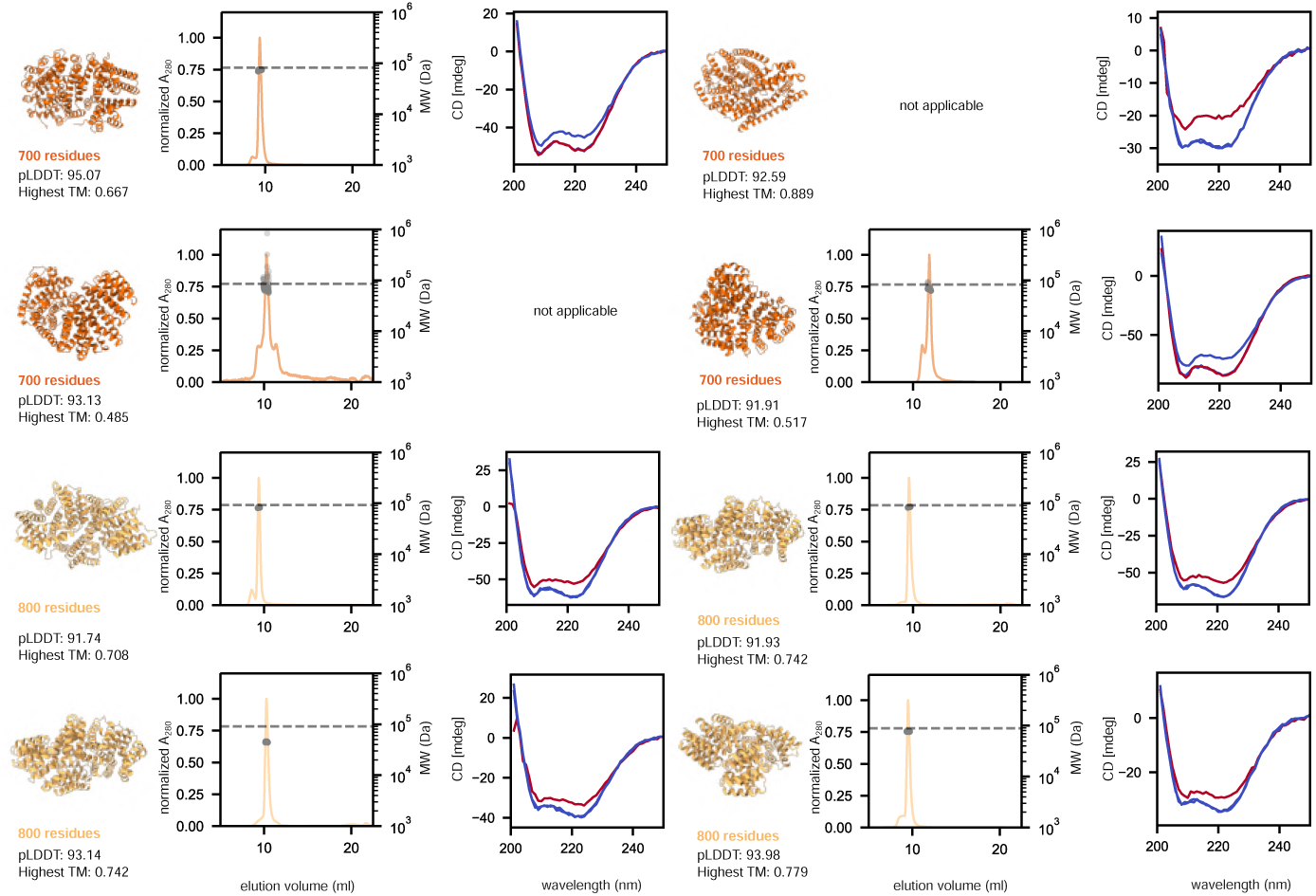
Biophysical characterization of designed monomeric folds: 700–800 residue designs. Layout as in Supplementary Fig. S1.

**Fig. S4.**
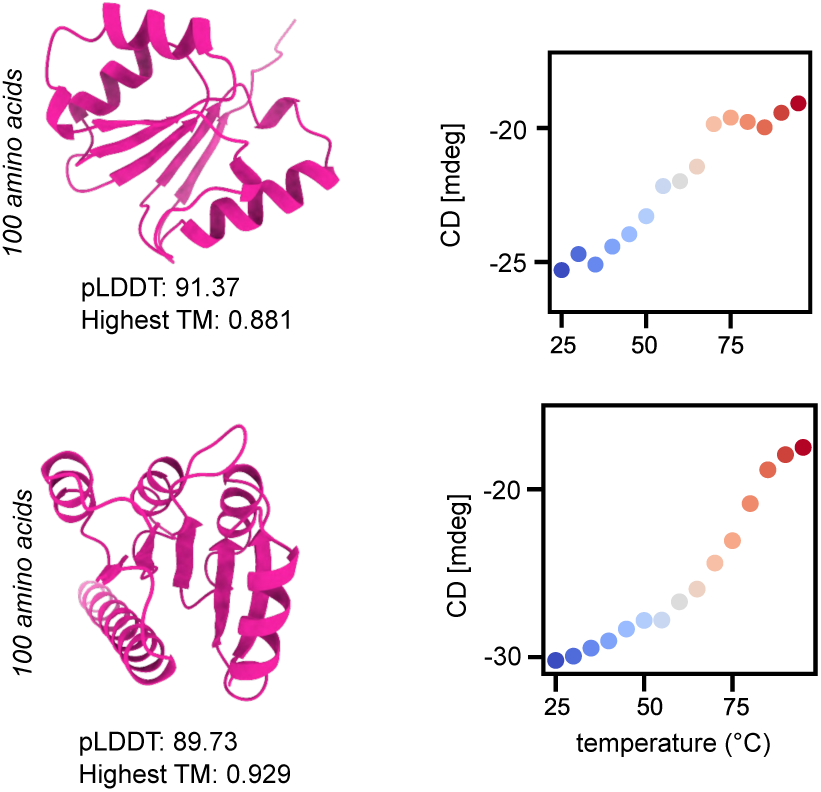
Biophysical characterization of monomers generated at high sampling temperature. Predicted structures and circular-dichroism thermal denaturation curves are shown for the two designs, from a selected set of 13 that expressed solubly. Both proteins undergo cooperative thermal transitions below 95 °C. The mean pLDDT from MSA-based AlphaFold2 predictions and the Fold-seek TM-score to the closest known fold are reported for each design.

**Fig. S5.**
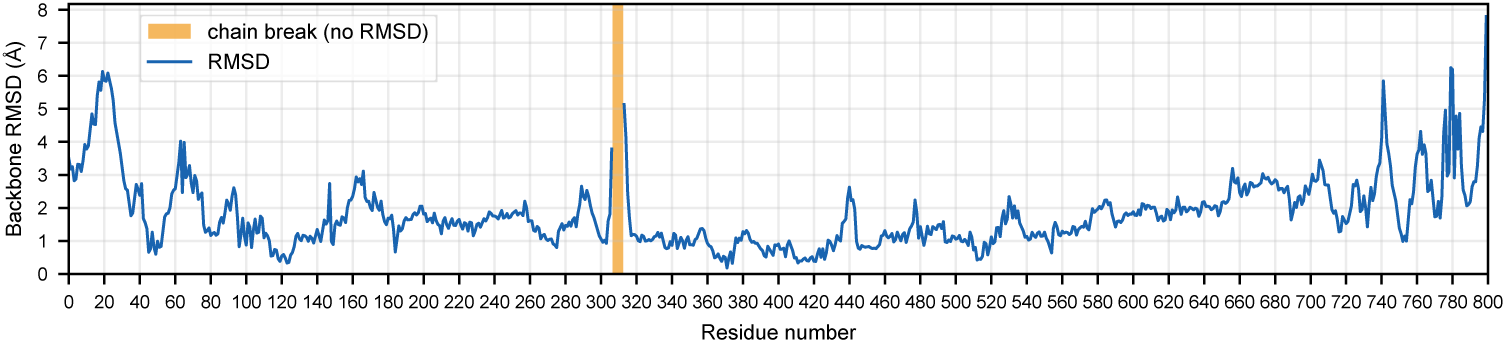
Root mean square deviation (RMSD) between crystal structure and design model. Plot shows the average per-residue RMSD between the backbone atom positions of the experimentally determined crystal structure and the generated design by La-Proteína. Residues 307–312 were not experimentally resolved.

**Fig. S6.**
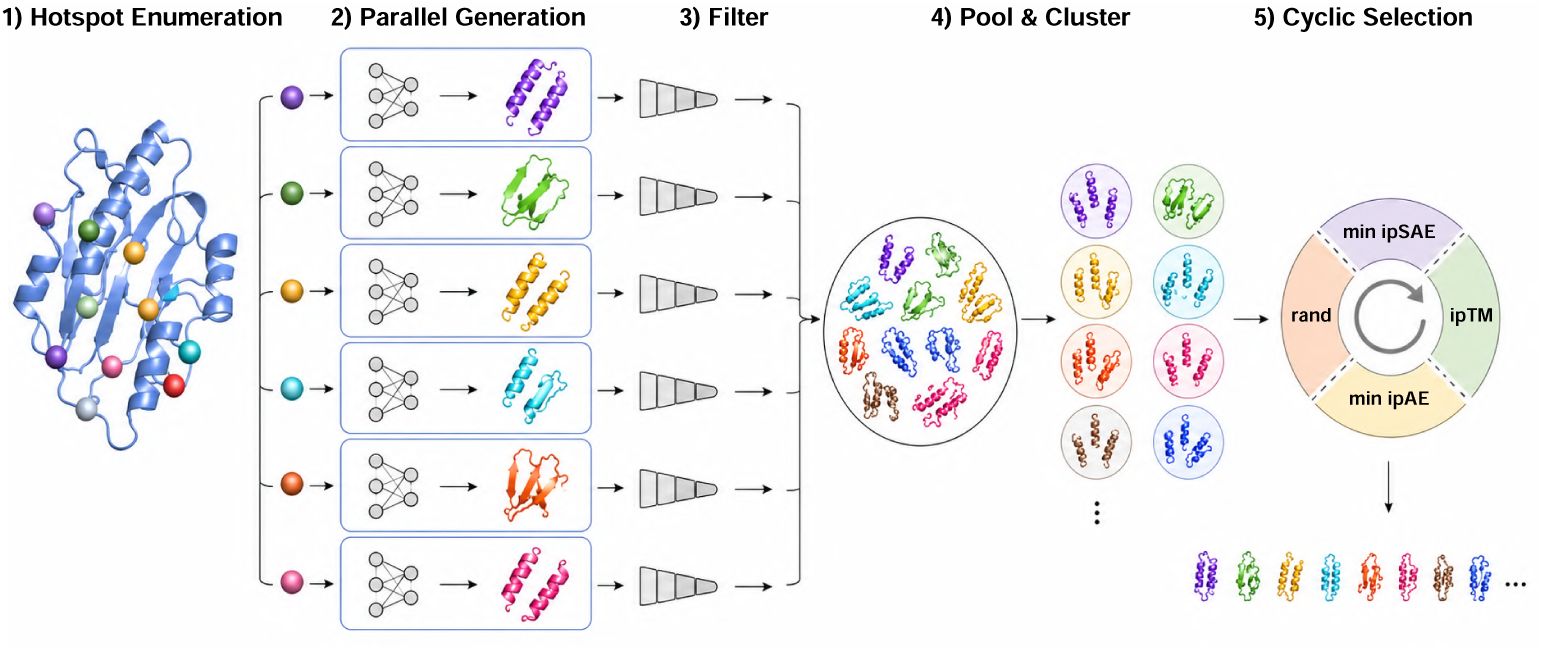
Massive-scale design and selection pipeline. The per-target procedure used to select the 492,636 Proteína-Complexa-generated sequence subset across the 127-target panel with diverse hotspots. (1) enumeration of candidate interface hotspot combinations; (2) parallel generation of candidates by beam-search-guided latent generative search; (3) in-silico filtering on interface confidence and physicochemical metrics; (4) pooling and structural clustering to remove redundancy; and (5) cyclic selection that rotates ranking criteria (minimum interface pAE, interface pTM, minimum interface pSAE and random selection) across clusters to assemble a diverse, size-matched set of designs for synthesis and multiplexed phage-display screening.

**Fig. S7.**
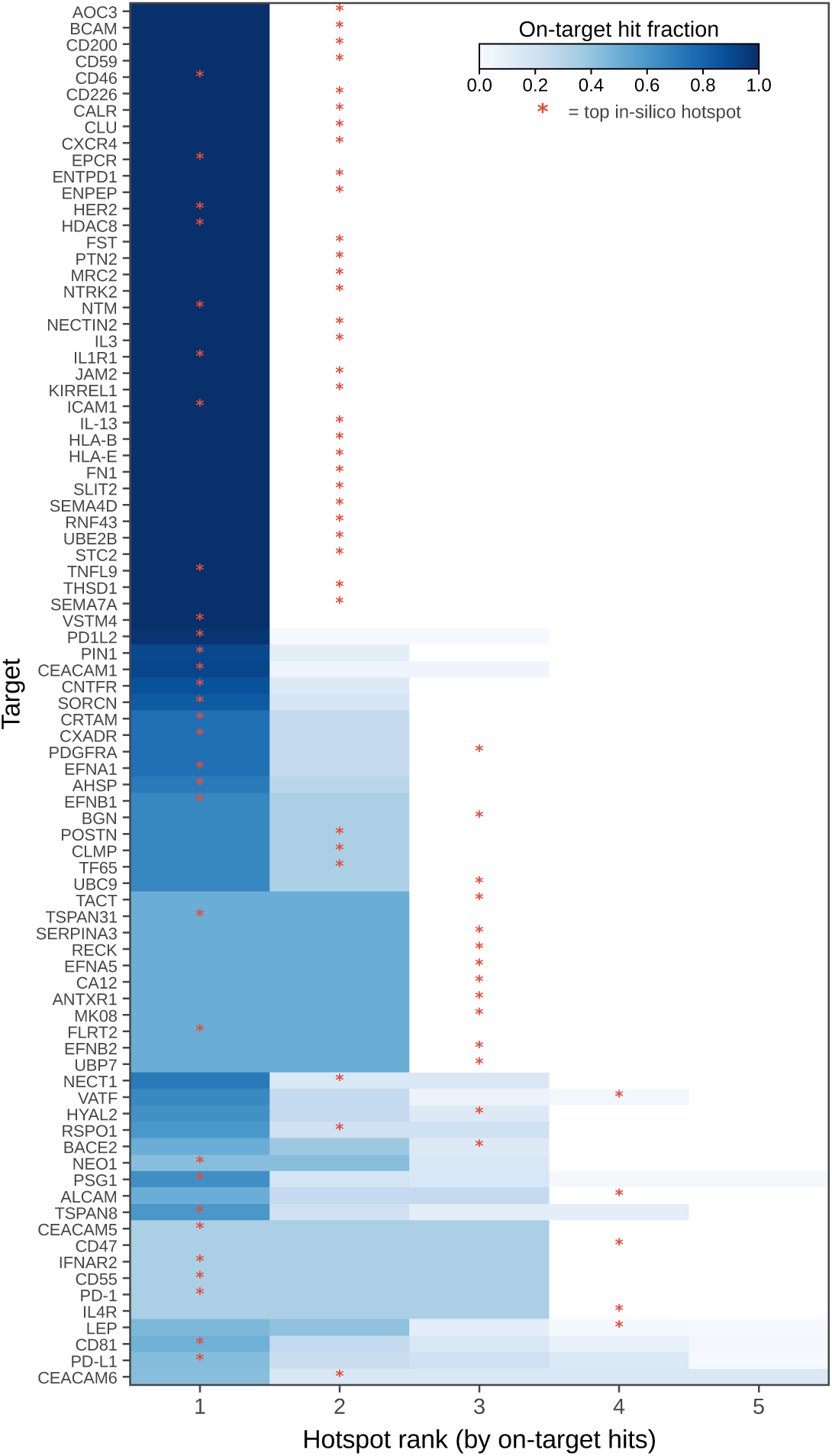
Per-target hotspot hit concentration analysis. Heatmap of on-target hit fraction across the top five hotspots for targets with at least one on-target hit. Targets are ordered by hotspot entropy. Red asterisks denote the top-ranked in-silico hotspot per target. Hit fractions are normalized over the top 10 hotspots per target.

**Fig. S8.**
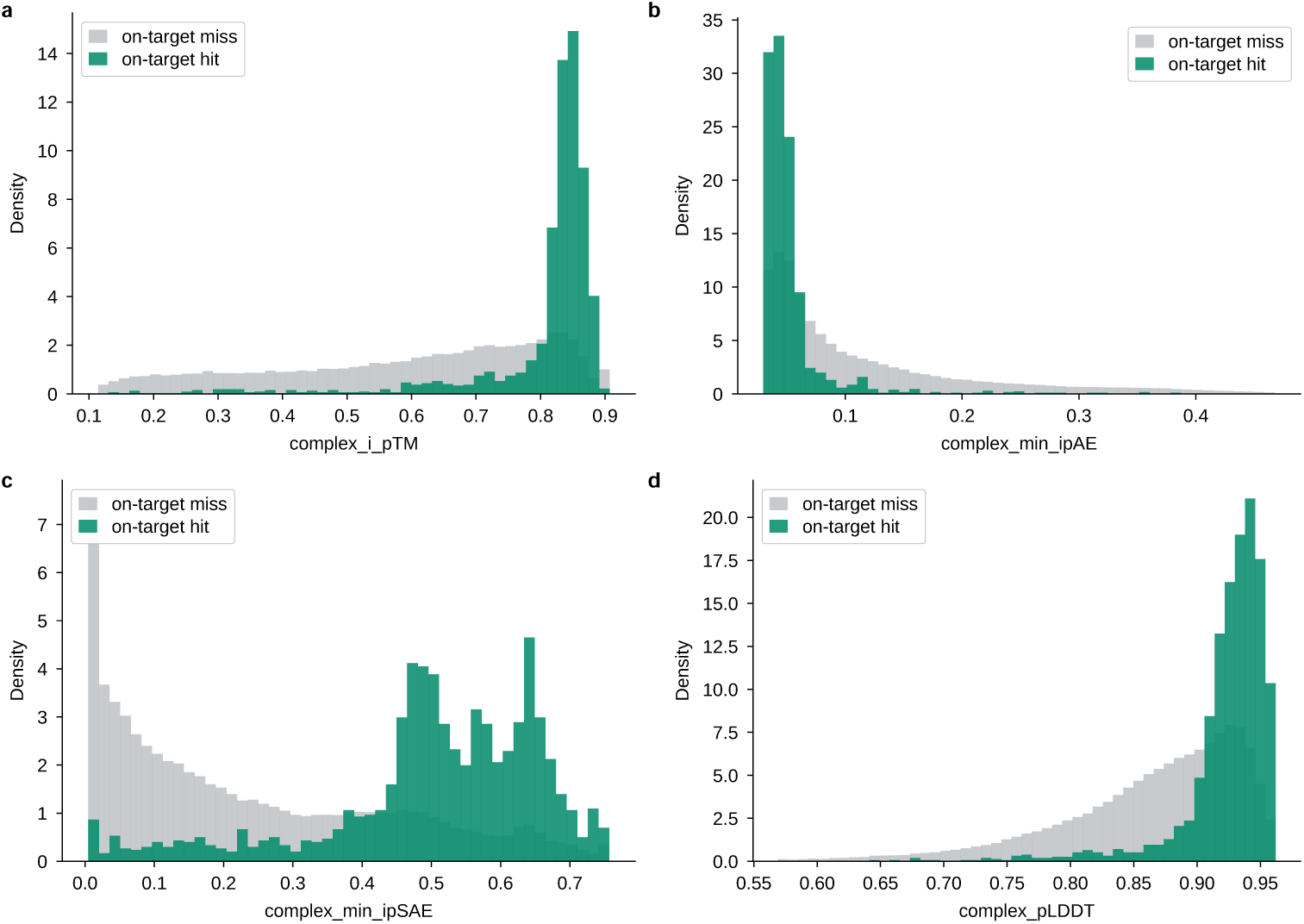
In-silico confidence scores separate experimental hits from misses. Distributions of AlphaFold2-Multimer complex confidence metrics — interface pTM (**a**), minimum interface pAE (**b**), minimum interface pSAE (**c**) and complex pLDDT (**d**) — for Proteína-Complexa designs classified as on-target hits (green) versus misses (grey) in the phage-display benchmark. Higher interface pTM, interface pSAE and pLDDT and lower interface pAE are enriched among experimental hits, indicating that these in-silico metrics are predictive of experimental success.

**Fig. S9.**
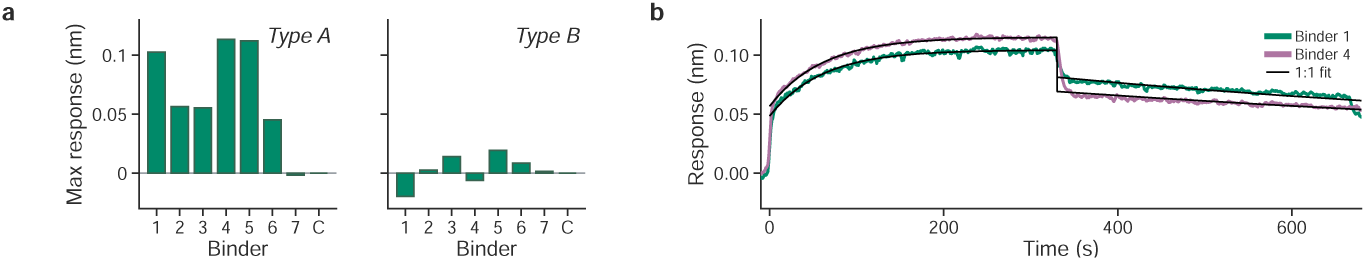
BLI specificity screen of carbohydrate-binder designs. **a**, Maximum biolayer-interferometry (BLI) association response of BLI-tested FP-positive designs, and a control (C), challenged with the type-A (left) versus the type-B (right) glycan conjugate, measured in *E. coli* lysate. Most designs respond strongly to the type-A glycan with little or no response to type-B, consistent with antigen-preferential binding. **b**, Representative association/dissociation sensorgrams for designs 1 and 4 against the type-A glycan, with 1:1 kinetic fits overlaid (black). Because the assay was run in crude lysate, these data are reported as supporting rather than definitive evidence of antigen specificity (see main text).

**Fig. S10.**
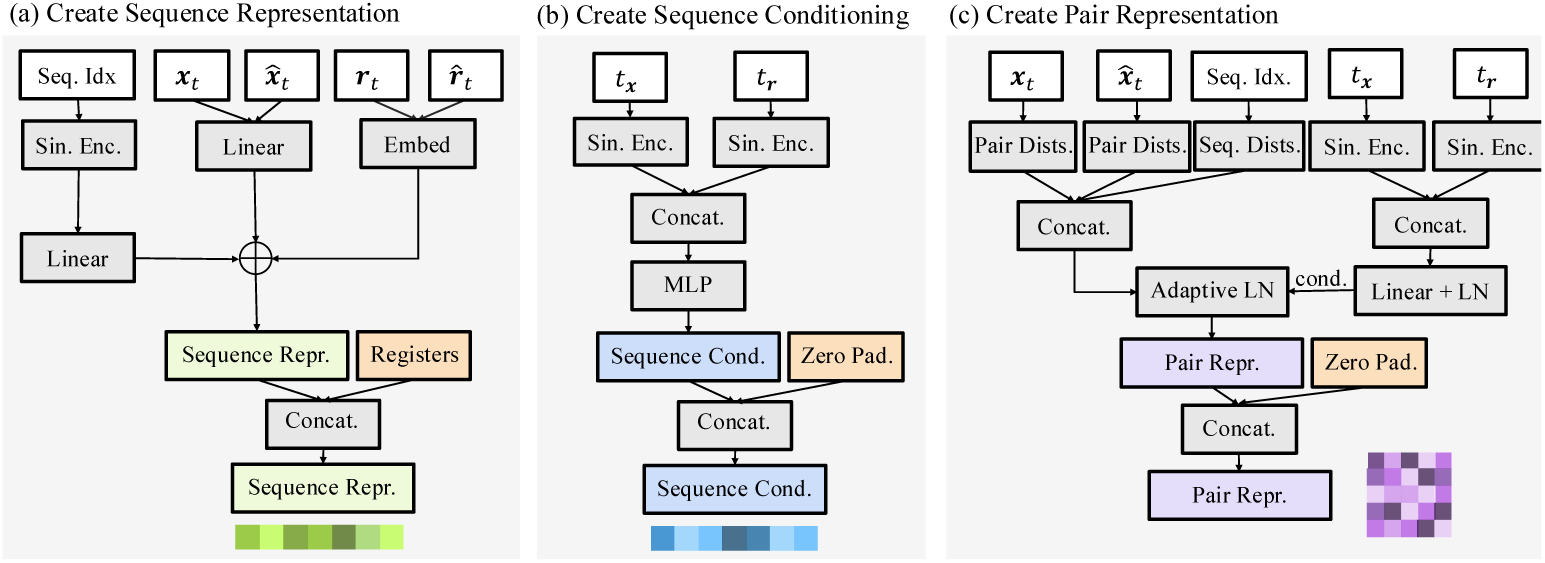
Construction of the inputs to the conditional flow denoiser. **a**, Sequence indices, coordinate channels, and latent or target-specific features are encoded and combined to form the initial sequence representation. Learned register tokens are then appended to the physical binder and target tokens. **b**, The backbone and latent interpolation times are sinusoidally encoded, concatenated, and passed through an MLP to produce the sequence conditioning representation. Zero-valued entries are appended for the register tokens. **c**, Pairwise coordinate distances, sequence separations, and the two time embeddings are combined to construct the initial pair representation. The geometric and sequence-derived pair features are adaptively normalized using the time conditioning before being concatenated with a transformed time embedding. Pair entries involving register tokens are zero-padded. The latent channel denoted by *r* and *t_r_* in the schematic corresponds to **z** and *t_z_* in the main text.

**Fig. S11.**
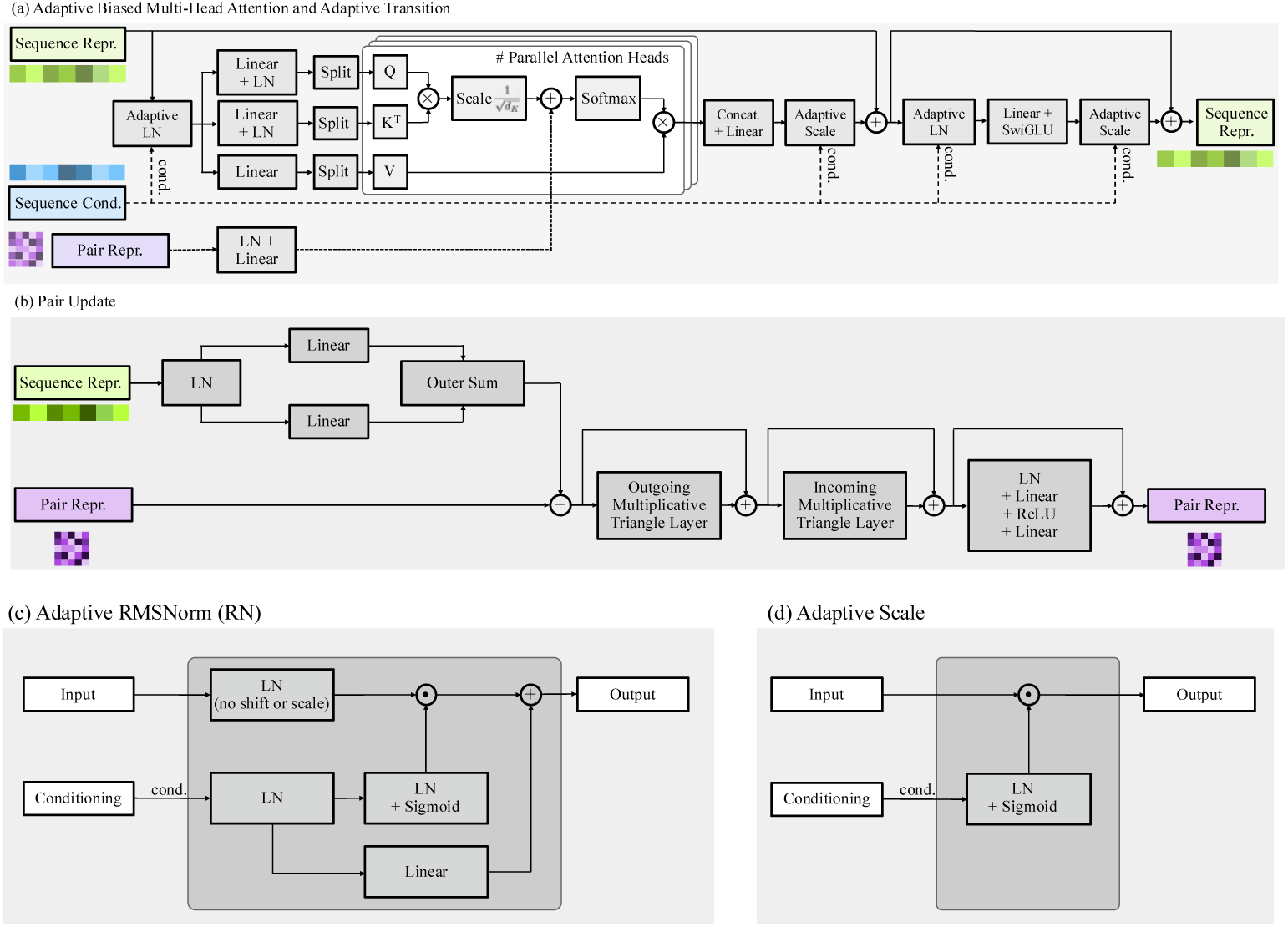
Architecture of one conditional sequence–pair transformer block. **a**, Adaptive pair-biased multi-head self-attention uses the pair representation to add a learned bias to the attention logits. Sequence conditioning modulates the normalization and residual scaling in both the attention and SwiGLU transition sublayers. **b**, The pair representation is updated from normalized sequence states through two learned projections and an elementwise product, followed by outgoing and incoming multiplicative triangle layers and a pair transition. Residual connections surround each pair update. **c**, Adaptive normalization uses the conditioning representation to generate a sigmoid gate and an additive shift for normalized activations. **d**, Adaptive scaling gates a residual branch using a sigmoid function of the normalized conditioning representation. In the diagram, RN denotes the base normalization operator, solid plus signs denote residual additions, and dashed connections carry conditioning signals or pair-derived attention biases.

**Fig. S12.**
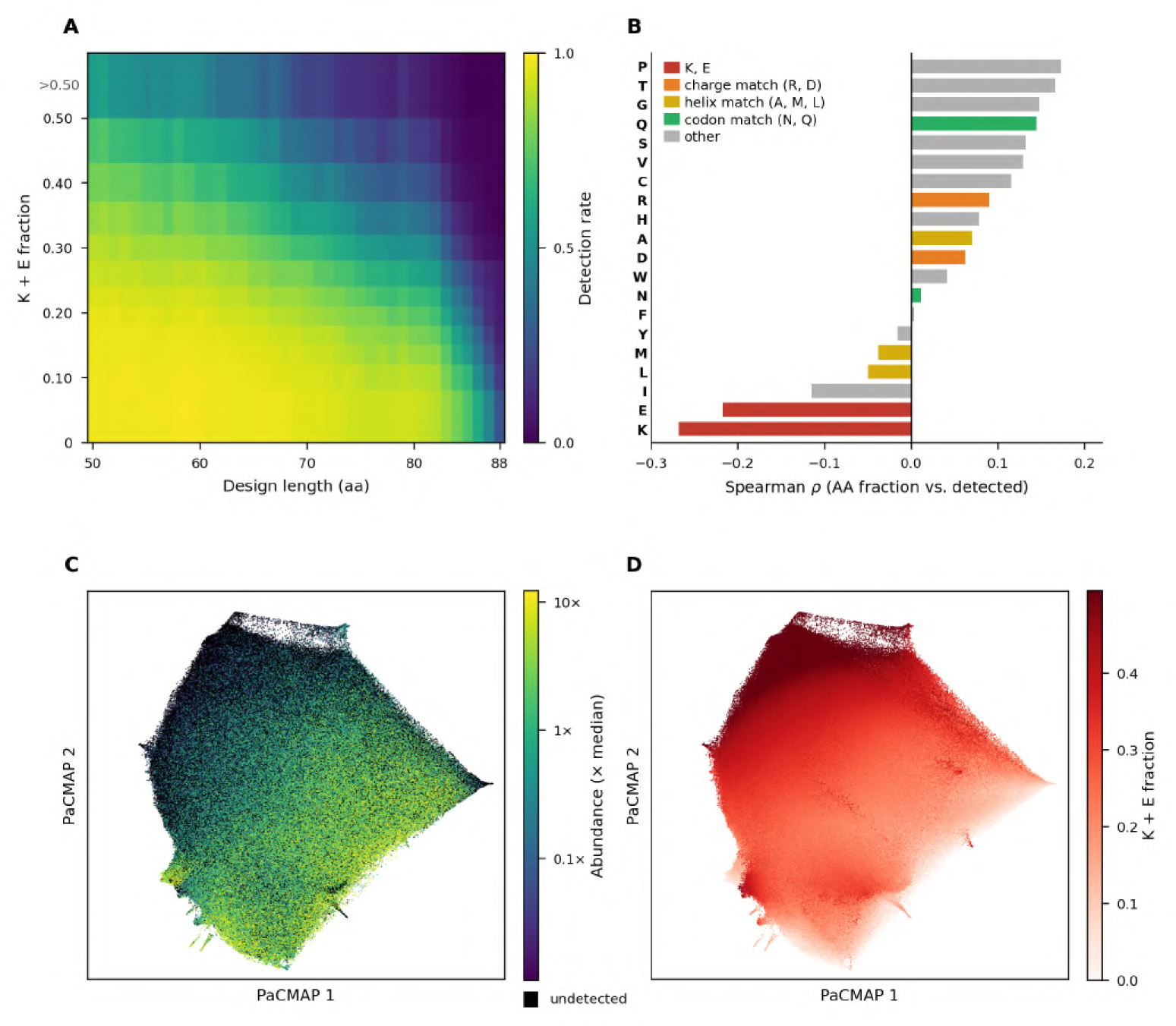
Lysine and glutamate content correlates with loss from the phage input pool. **a**, Detection rate as a function of design length and combined K+E fraction across 1.4 million *de novo* designs in four length-stratified libraries. **b**, Spearman correlation between per-residue fractional composition and detection. **c**, PaCMAP embedding of all designs, highlighting undetected sequences and relative abundance among detected sequences. **d**, Same embedding colored by K+E fraction.

**Fig. S13.**
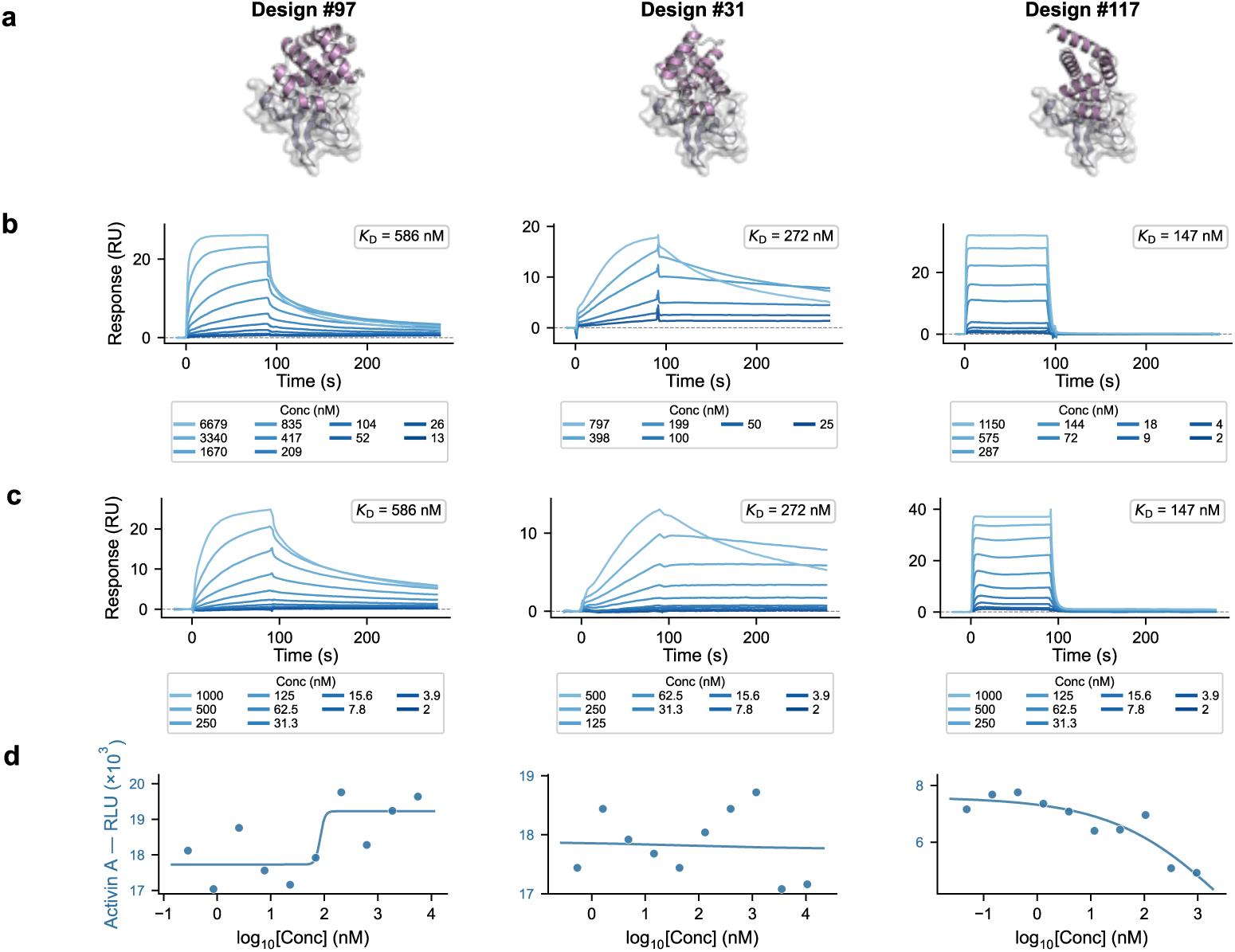
Experimental characterization of three additional ActRIIA binder designs. Designs #97, #31 and #117 achieved sub-*µ*M binding affinity but did not exhibit significant functional inhibition. **a**, Predicted complex structures of each binder (purple) engaged with the ActRIIA receptor (gray surface). **b**, SPR sensorgrams from the first independent measurement (raw traces only), with equilibrium *K*_D_ values indicated. **c**, SPR sensorgrams from the second independent measurement (raw traces only). **d**, Dose-response curves from the Smad2/3-responsive luciferase reporter assay under Activin A stimulation (20 ng/mL). Despite nanomolar-to-sub-*µ*M affinity for ActRIIA, these designs did not produce measurable functional inhibition at the tested concentrations, suggesting that binding orientation and/or epitope overlap was insufficient to block native ligand signaling.

**Fig. S14.**
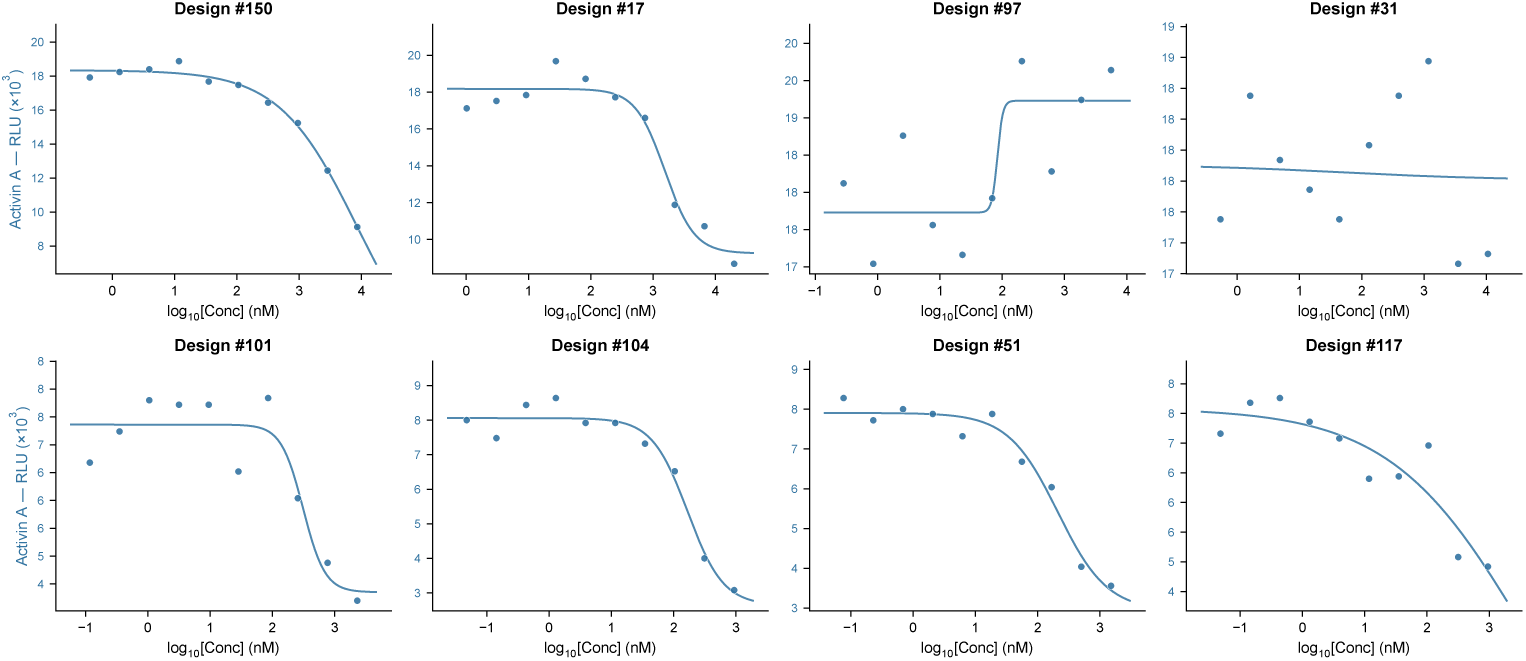
Activin A functional assay across all eight ActRIIA binder designs. Dose-response curves from the Smad2/3 luciferase reporter assay are shown for all eight designs tested in this campaign (blue curves). Luminescence values are plotted against log_10_ concentration (nM). This panel complements the focused main-text figure by showing the full set of experimentally characterized designs under Activin A stimulation conditions.

**Fig. S15.**
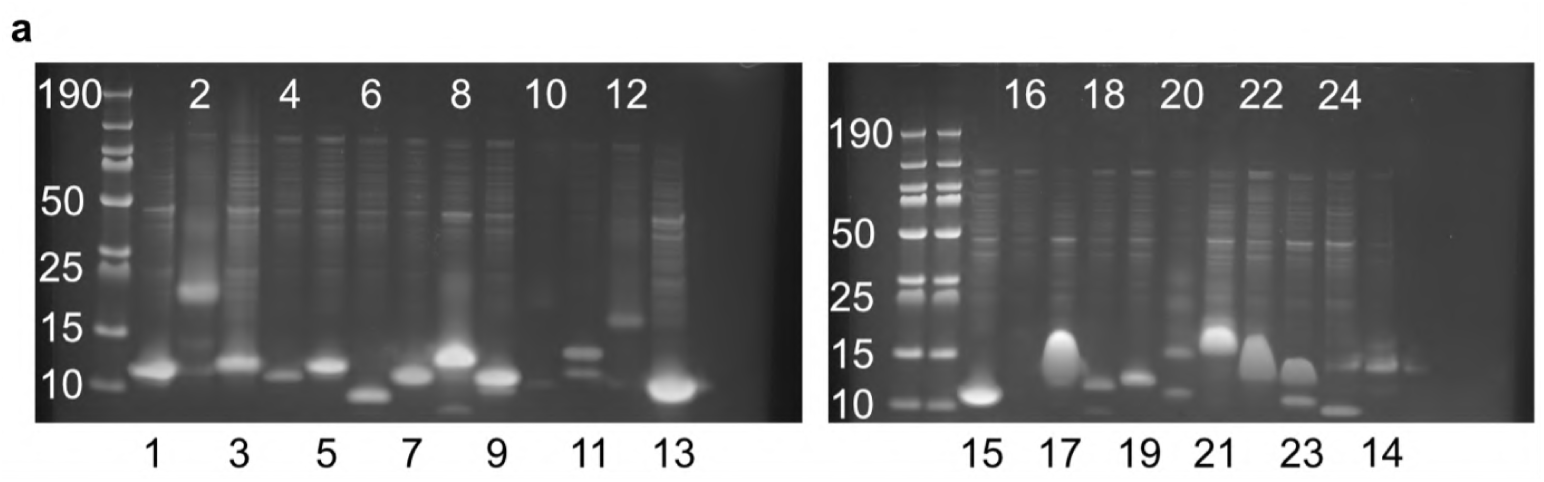
Expression of the initial carbohydrate-binder candidates. SDS-PAGE analysis of the 24 candidates expressed in *E. coli*. Lanes S1–S24 correspond to the expression-test panel; molecular-weight markers (kDa) are shown. Twenty-three designs advanced to the binding screen reported in the main text.

## S9 Supplementary tables

The following tables supplement the main text and Supplementary Methods.

**Table S1.** Temperature ablation for Proteina-Complexa (self-generated sequences) on baseline solvable targets (*n* = 83). On-target hit counts, specificity (specific/all on-target), and target coverage across backbone and local latent temperature combinations. New and lost columns indicate targets gained or lost relative to the default (0.1, 0.1) setting; unique denotes targets solved exclusively by that condition and no other.

| Temperature | On-target |  |  | Targets |  |  |  | # Seqs. |
| --- | --- | --- | --- | --- | --- | --- | --- | --- |
| (backbone, local latent) | All | Specific | Specificity | Solved | New | Lost | Unique |  |
| (0.1, 0.1) | **702** | **607** | **86.5%** | 35 | – | – | 4 | 49,263 |
| (0.1, 0.4) | 495 | 419 | 84.6% | 34 | +12 | –13 | 6 | 45,255 |
| (0.4, 0.1) | 492 | 412 | 83.7% | 30 | +6 | –11 | 3 | 45,904 |
| (0.4, 0.4) | 318 | 232 | 73.0% | 30 | +7 | –12 | 4 | 49,697 |

**Table S2.**
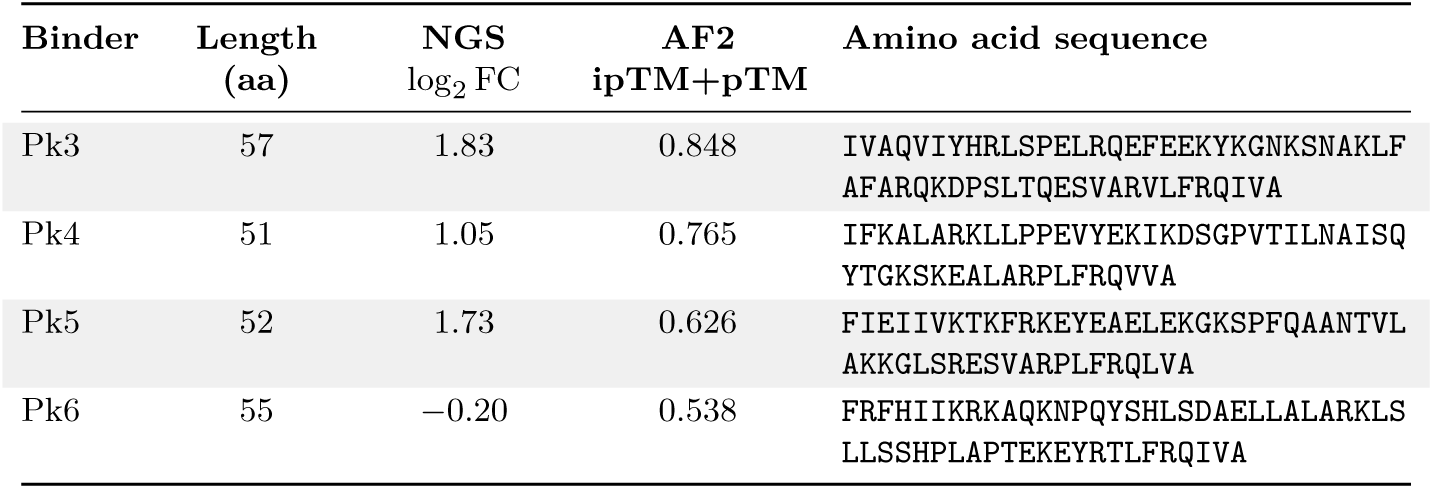
Properties of selected PAK1 mini-protein binder candidates. NGS log_2_ FC represents the log_2_ fold-change in read frequency relative to the input library after kanamycin selection. AF2 ipTM+pTM is the combined score from AlphaFold2 Multimer (best of 25 models).

**Table S3.**
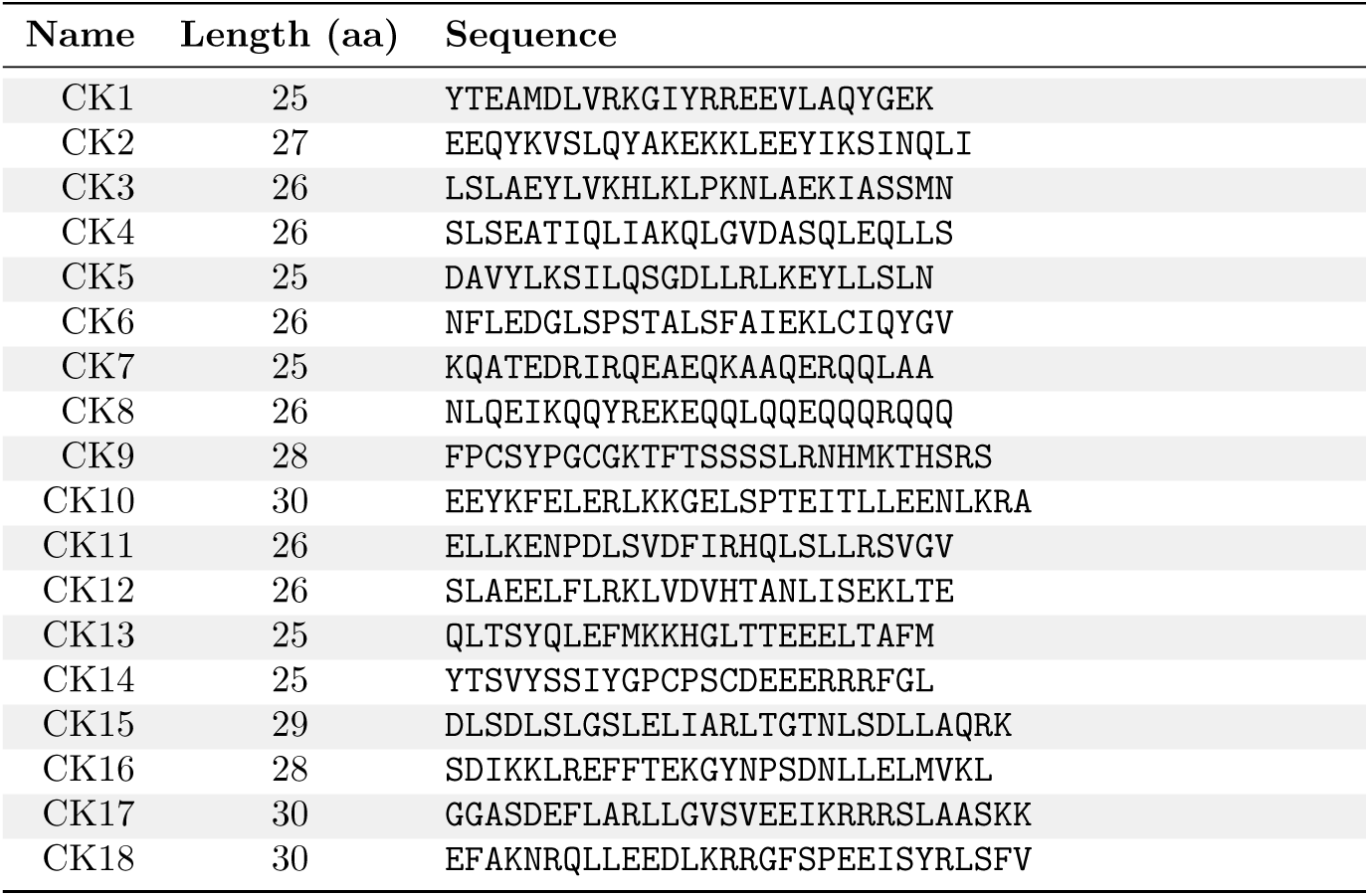
Predicted CK1*δ* peptide binders. All peptides were synthesized with an N-terminal biotinylated lysine followed by two glycine spacers.

**Table S4:**
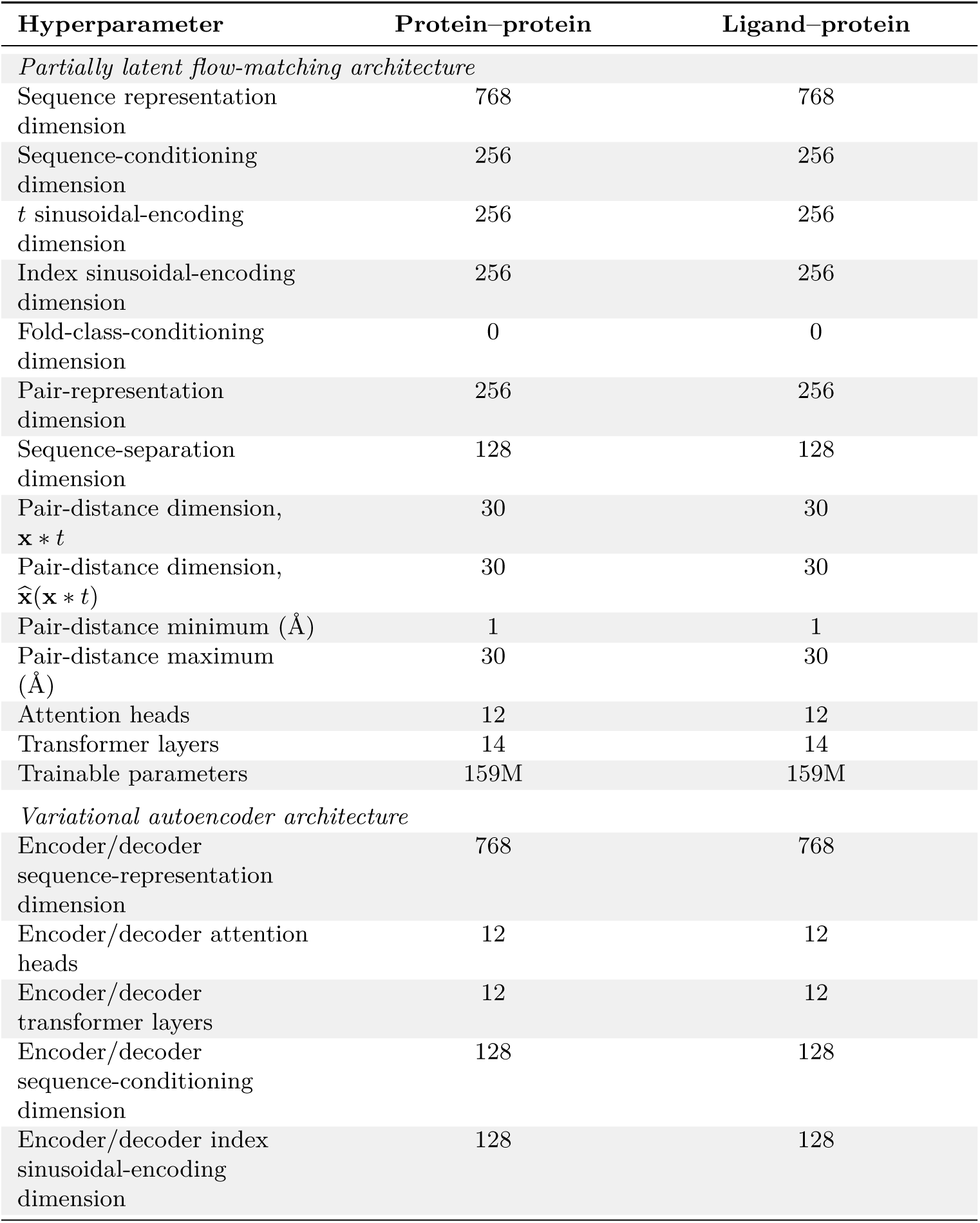
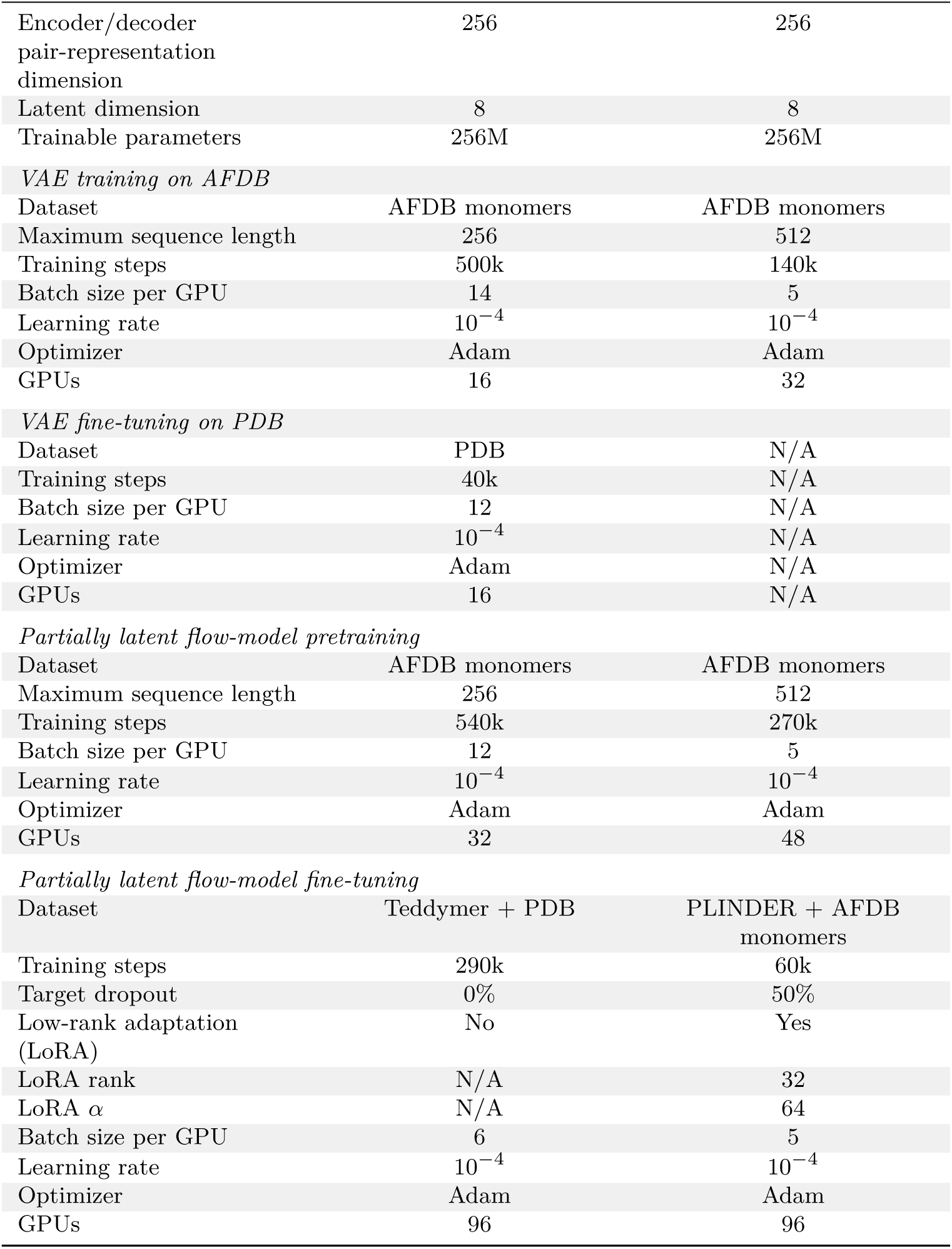
Architecture and training hyperparameters for Proteína-Complexa. Values are reproduced from Appendix G and Table 14 of ref. [18]. Protein–protein denotes the model used for protein-target binder generation, and ligand–protein the model used for small-molecule targets. All GPU counts refer to NVIDIA A100-80GB devices. N/A, not applicable.

**Table S5:** Reward functions and downstream selection criteria used in this study. The table consolidates the campaign-specific definitions reported in the main text and Methods. Unless stated otherwise, beam search used 400 denoising steps, beam width *N* = 4, branch factor *L* = 4 and update interval *K* = 100; Feynman–Kac steering used inverse temperature *β* = 10 [18].

| Campaign or task | Search procedure | Reward used during search | Downstream success or selection criteria |
| --- | --- | --- | --- |
| Designed monomers | Generation followed by filtering | None since pure generative model benchmarking | AlphaFold2 single-sequence mean pLDDT $> 90$ and backbone RMSD $< 2 \text{ \AA}$ to the design model |
| PDGFR and PD-L1 | Beam search | AlphaFold2-Multimer interface predicted aligned error (interface pAE) | Physicochemical and confidence filtering, TM-score clustering at 0.8, then OpenFold3 and Boltz-2 monomer validation with pLDDT $> 80$ and RMSD $< 2 \text{ \AA}$ |
| Nipah virus G, *de novo* and scaffold re-engineering | Beam search | Multi-objective combination of Boltz-2 maximum interface score, shape complementarity, interface hydrophobicity and hydrogen-bond satisfaction | Top 10 *de novo* and top 10 scaffold reengineering variants were selected ipAE, TMol hydrogen bonds, shape complementarity and visual inspection for key interactions |
| Massive-scale 127-target benchmark | Beam search | AlphaFold2-Multimer interface pAE | Minimum interface pAE $< 2$, complex pLDDT $> 0.8$, interface pTM $> 0.7$ and fewer than five $C_\alpha$ clashes; diversity selection cycled rankings across minimum interface pAE, interface pTM, minimum interface pSAE and random selection |
| ActRIIA | Beam search, Feynman–Kac steering or Monte Carlo tree search across seven rounds | AlphaFold2-Multimer interface pAE | Ranking by the averaged score followed by Foldseek scaffold clustering and experimental surface-plasmon-resonance testing |
| PAK1 mini-proteins | Beam search | AlphaFold2-Multimer interface pAE (lower is better) | Fifty best selected candidates by ipAE were tested |

**Table S5: Reward functions and downstream selection criteria used in this study** (continued).
| Campaign or task | Search procedure | Reward used during search | Downstream success or selection criteria |
| --- | --- | --- | --- |
| CK1δ peptides | Beam Search | AlphaFold2-Multimer interface pAE | 18 best candidates were selected by ipAE |
| Carbohydrate antigen | Beam search | Pocket burial, physics-based Tmol hydrogen-bond energies and RosettaFold3 interface confidence; after each search step the RosettaFold3-predicted sugar pose replaced the preceding pose | RosettaFold3 minimum interface pAE < 2, binder RMSD < 2 Å and ligand RMSD < 5 Å |

**Table S6:**
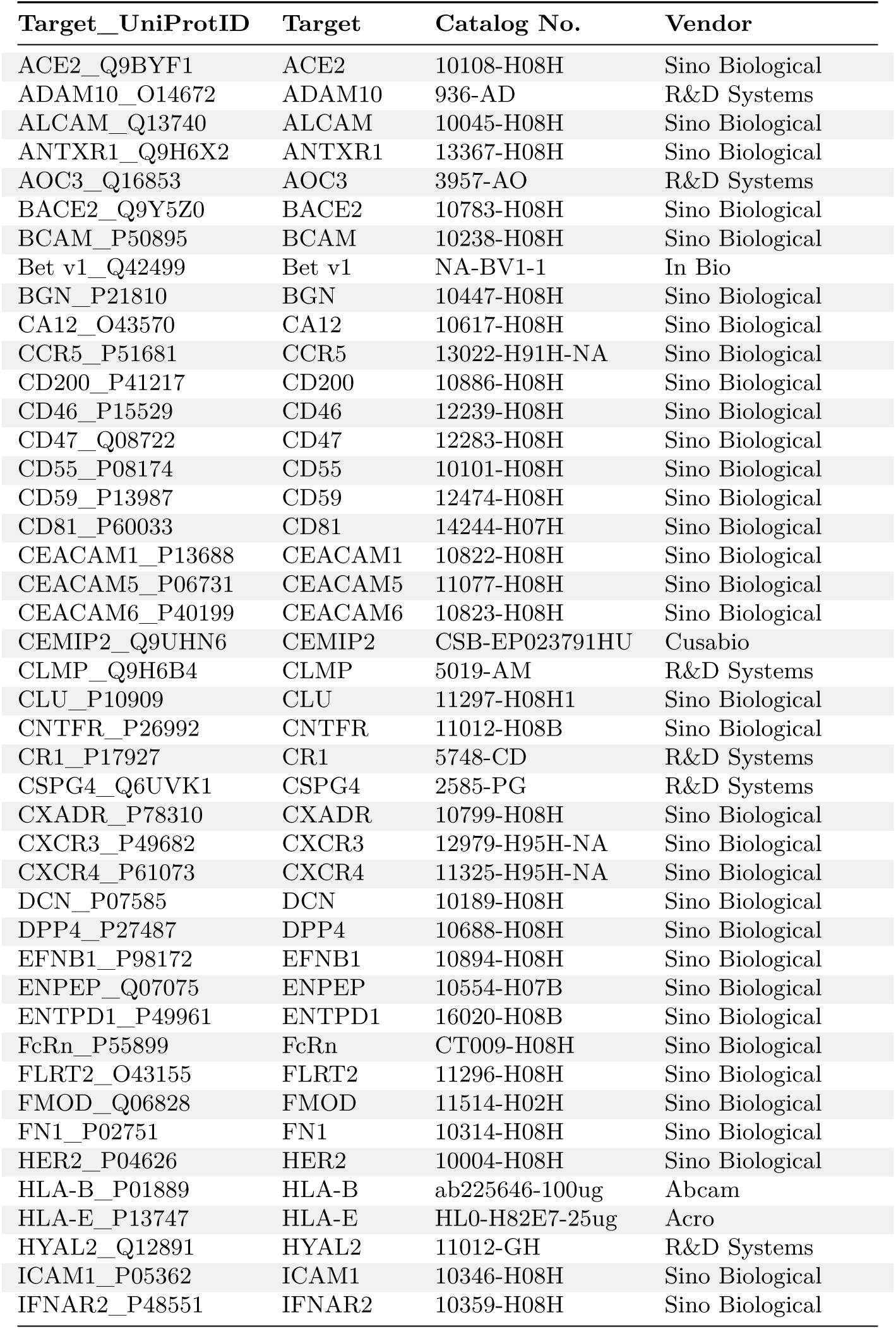
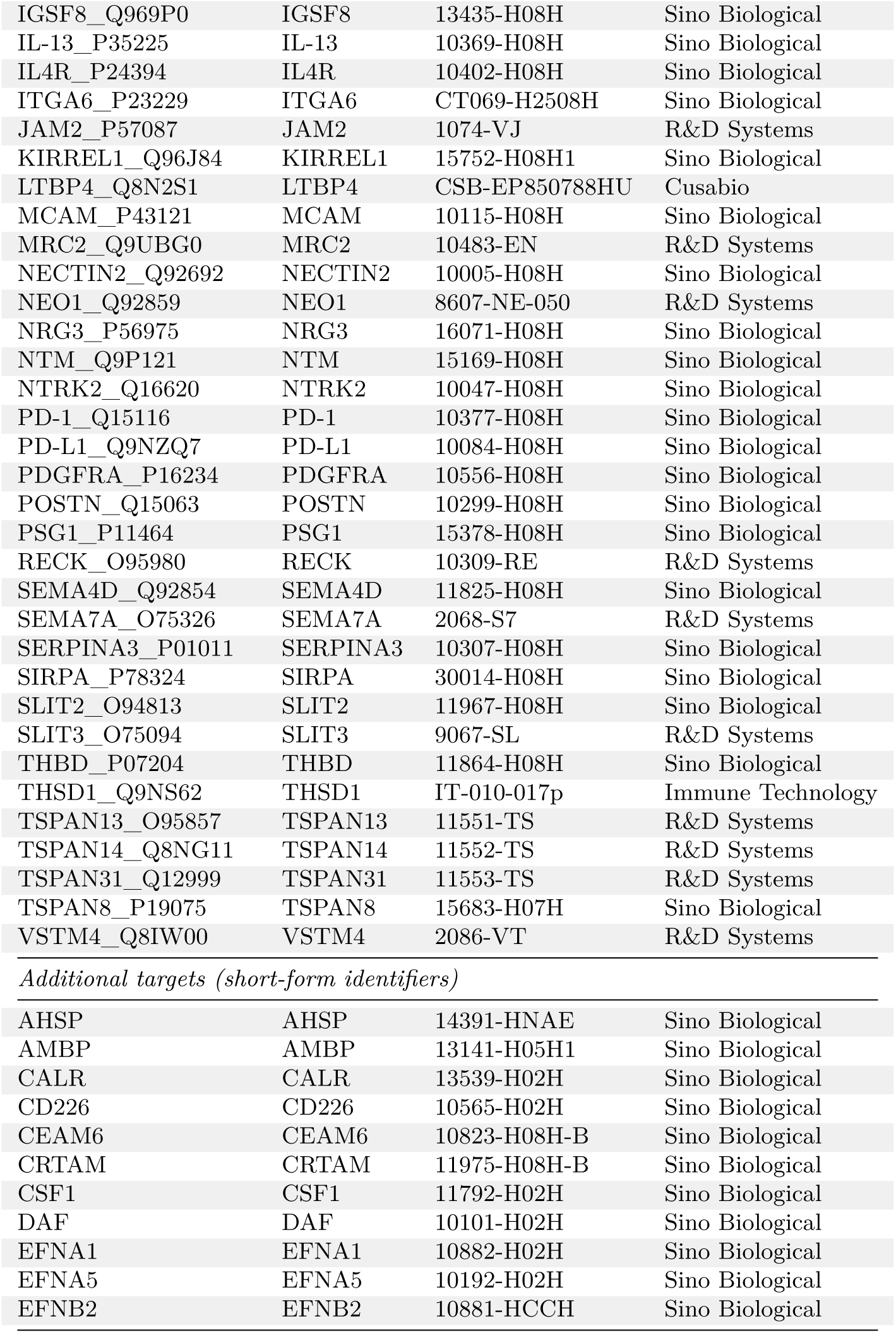
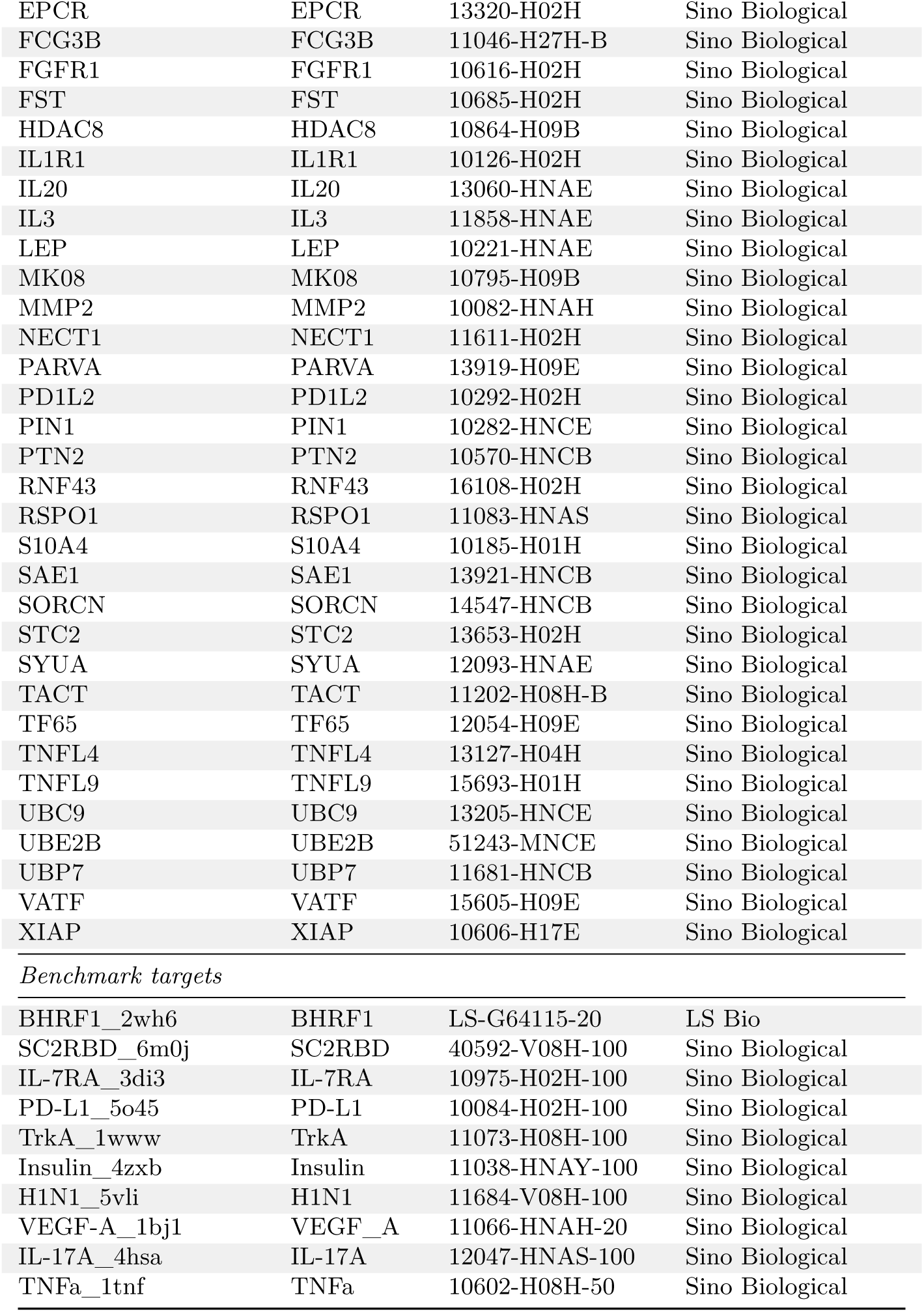
List of commercially sourced proteins used in the Manifold Bio large-scale all-by-all binder screen.

| Target_UniProtID | Target | Catalog No. | Vendor |
| --- | --- | --- | --- |
| ACE2_Q9BYF1 | ACE2 | 10108-H08H | Sino Biological |
| ADAM10_O14672 | ADAM10 | 936-AD | R&D Systems |
| ALCAM_Q13740 | ALCAM | 10045-H08H | Sino Biological |
| ANTXR1_Q9H6X2 | ANTXR1 | 13367-H08H | Sino Biological |
| AOC3_Q16853 | AOC3 | 3957-AO | R&D Systems |
| BACE2_Q9Y5Z0 | BACE2 | 10783-H08H | Sino Biological |
| BCAM_P50895 | BCAM | 10238-H08H | Sino Biological |
| Bet v1_Q42499 | Bet v1 | NA-BV1-1 | In Bio |
| BGN_P21810 | BGN | 10447-H08H | Sino Biological |
| CA12_O43570 | CA12 | 10617-H08H | Sino Biological |
| CCR5_P51681 | CCR5 | 13022-H91H-NA | Sino Biological |
| CD200_P41217 | CD200 | 10886-H08H | Sino Biological |
| CD46_P15529 | CD46 | 12239-H08H | Sino Biological |
| CD47_Q08722 | CD47 | 12283-H08H | Sino Biological |
| CD55_P08174 | CD55 | 10101-H08H | Sino Biological |
| CD59_P13987 | CD59 | 12474-H08H | Sino Biological |
| CD81_P60033 | CD81 | 14244-H07H | Sino Biological |
| CEACAM1_P13688 | CEACAM1 | 10822-H08H | Sino Biological |
| CEACAM5_P06731 | CEACAM5 | 11077-H08H | Sino Biological |
| CEACAM6_P40199 | CEACAM6 | 10823-H08H | Sino Biological |
| CEMIP2_Q9UHN6 | CEMIP2 | CSB-EP023791HU | Cusabio |
| CLMP_Q9H6B4 | CLMP | 5019-AM | R&D Systems |
| CLU_P10909 | CLU | 11297-H08H1 | Sino Biological |
| CNTFR_P26992 | CNTFR | 11012-H08B | Sino Biological |
| CR1_P17927 | CR1 | 5748-CD | R&D Systems |
| CSPG4_Q6UVK1 | CSPG4 | 2585-PG | R&D Systems |
| CXADR_P78310 | CXADR | 10799-H08H | Sino Biological |
| CXCR3_P49682 | CXCR3 | 12979-H95H-NA | Sino Biological |
| CXCR4_P61073 | CXCR4 | 11325-H95H-NA | Sino Biological |
| DCN_P07585 | DCN | 10189-H08H | Sino Biological |
| DPP4_P27487 | DPP4 | 10688-H08H | Sino Biological |
| EFNB1_P98172 | EFNB1 | 10894-H08H | Sino Biological |
| ENPEP_Q07075 | ENPEP | 10554-H07B | Sino Biological |
| ENTPD1_P49961 | ENTPD1 | 16020-H08B | Sino Biological |
| FcRn_P55899 | FcRn | CT009-H08H | Sino Biological |
| FLRT2_O43155 | FLRT2 | 11296-H08H | Sino Biological |
| FMOD_Q06828 | FMOD | 11514-H02H | Sino Biological |
| FN1_P02751 | FN1 | 10314-H08H | Sino Biological |
| HER2_P04626 | HER2 | 10004-H08H | Sino Biological |
| HLA-B_P01889 | HLA-B | ab225646-100ug | Abcam |
| HLA-E_P13747 | HLA-E | HL0-H82E7-25ug | Acro |
| HYAL2_Q12891 | HYAL2 | 11012-GH | R&D Systems |
| ICAM1_P05362 | ICAM1 | 10346-H08H | Sino Biological |
| IFNAR2_P48551 | IFNAR2 | 10359-H08H | Sino Biological |

Table S6 – *continued from previous page*
| Target_UniProtID | Target | Catalog No. | Vendor |
| --- | --- | --- | --- |
| IGSF8_Q969P0 | IGSF8 | 13435-H08H | Sino Biological |
| IL-13_P35225 | IL-13 | 10369-H08H | Sino Biological |
| IL4R_P24394 | IL4R | 10402-H08H | Sino Biological |
| ITGA6_P23229 | ITGA6 | CT069-H2508H | Sino Biological |
| JAM2_P57087 | JAM2 | 1074-VJ | R&D Systems |
| KIRREL1_Q96J84 | KIRREL1 | 15752-H08H1 | Sino Biological |
| LTBP4_Q8N2S1 | LTBP4 | CSB-EP850788HU | Cusabio |
| MCAM_P43121 | MCAM | 10115-H08H | Sino Biological |
| MRC2_Q9UBG0 | MRC2 | 10483-EN | R&D Systems |
| NECTIN2_Q92692 | NECTIN2 | 10005-H08H | Sino Biological |
| NEO1_Q92859 | NEO1 | 8607-NE-050 | R&D Systems |
| NRG3_P56975 | NRG3 | 16071-H08H | Sino Biological |
| NTM_Q9P121 | NTM | 15169-H08H | Sino Biological |
| NTRK2_Q16620 | NTRK2 | 10047-H08H | Sino Biological |
| PD-1_Q15116 | PD-1 | 10377-H08H | Sino Biological |
| PD-L1_Q9NZQ7 | PD-L1 | 10084-H08H | Sino Biological |
| PDGFRA_P16234 | PDGFRA | 10556-H08H | Sino Biological |
| POSTN_Q15063 | POSTN | 10299-H08H | Sino Biological |
| PSG1_P11464 | PSG1 | 15378-H08H | Sino Biological |
| RECK_O95980 | RECK | 10309-RE | R&D Systems |
| SEMA4D_Q92854 | SEMA4D | 11825-H08H | Sino Biological |
| SEMA7A_O75326 | SEMA7A | 2068-S7 | R&D Systems |
| SERPINA3_P01011 | SERPINA3 | 10307-H08H | Sino Biological |
| SIRPA_P78324 | SIRPA | 30014-H08H | Sino Biological |
| SLIT2_O94813 | SLIT2 | 11967-H08H | Sino Biological |
| SLIT3_O75094 | SLIT3 | 9067-SL | R&D Systems |
| THBD_P07204 | THBD | 11864-H08H | Sino Biological |
| THSD1_Q9NS62 | THSD1 | IT-010-017p | Immune Technology |
| TSPAN13_O95857 | TSPAN13 | 11551-TS | R&D Systems |
| TSPAN14_Q8NG11 | TSPAN14 | 11552-TS | R&D Systems |
| TSPAN31_Q12999 | TSPAN31 | 11553-TS | R&D Systems |
| TSPAN8_P19075 | TSPAN8 | 15683-H07H | Sino Biological |
| VSTM4_Q8IW00 | VSTM4 | 2086-VT | R&D Systems |
| *Additional targets (short-form identifiers)* |  |  |  |
| AHSP | AHSP | 14391-HNAE | Sino Biological |
| AMBP | AMBP | 13141-H05H1 | Sino Biological |
| CALR | CALR | 13539-H02H | Sino Biological |
| CD226 | CD226 | 10565-H02H | Sino Biological |
| CEAM6 | CEAM6 | 10823-H08H-B | Sino Biological |
| CRTAM | CRTAM | 11975-H08H-B | Sino Biological |
| CSF1 | CSF1 | 11792-H02H | Sino Biological |
| DAF | DAF | 10101-H02H | Sino Biological |
| EFNA1 | EFNA1 | 10882-H02H | Sino Biological |
| EFNA5 | EFNA5 | 10192-H02H | Sino Biological |
| EFNB2 | EFNB2 | 10881-HCCH | Sino Biological |

Table S6 – *continued from previous page*
| Target_UniProtID | Target | Catalog No. | Vendor |
| --- | --- | --- | --- |
| EPCR | EPCR | 13320-H02H | Sino Biological |
| FCG3B | FCG3B | 11046-H27H-B | Sino Biological |
| FGFR1 | FGFR1 | 10616-H02H | Sino Biological |
| FST | FST | 10685-H02H | Sino Biological |
| HDAC8 | HDAC8 | 10864-H09B | Sino Biological |
| IL1R1 | IL1R1 | 10126-H02H | Sino Biological |
| IL20 | IL20 | 13060-HNAE | Sino Biological |
| IL3 | IL3 | 11858-HNAE | Sino Biological |
| LEP | LEP | 10221-HNAE | Sino Biological |
| MK08 | MK08 | 10795-H09B | Sino Biological |
| MMP2 | MMP2 | 10082-HNAH | Sino Biological |
| NECT1 | NECT1 | 11611-H02H | Sino Biological |
| PARVA | PARVA | 13919-H09E | Sino Biological |
| PD1L2 | PD1L2 | 10292-H02H | Sino Biological |
| PIN1 | PIN1 | 10282-HNCE | Sino Biological |
| PTN2 | PTN2 | 10570-HNCB | Sino Biological |
| RNF43 | RNF43 | 16108-H02H | Sino Biological |
| RSPO1 | RSPO1 | 11083-HNAS | Sino Biological |
| S10A4 | S10A4 | 10185-H01H | Sino Biological |
| SAE1 | SAE1 | 13921-HNCB | Sino Biological |
| SORCN | SORCN | 14547-HNCB | Sino Biological |
| STC2 | STC2 | 13653-H02H | Sino Biological |
| SYUA | SYUA | 12093-HNAE | Sino Biological |
| TACT | TACT | 11202-H08H-B | Sino Biological |
| TF65 | TF65 | 12054-H09E | Sino Biological |
| TNFL4 | TNFL4 | 13127-H04H | Sino Biological |
| TNFL9 | TNFL9 | 15693-H01H | Sino Biological |
| UBC9 | UBC9 | 13205-HNCE | Sino Biological |
| UBE2B | UBE2B | 51243-MNCE | Sino Biological |
| UBP7 | UBP7 | 11681-HNCB | Sino Biological |
| VATF | VATF | 15605-H09E | Sino Biological |
| XIAP | XIAP | 10606-H17E | Sino Biological |
| *Benchmark targets* |  |  |  |
| BHRF1_2wh6 | BHRF1 | LS-G64115-20 | LS Bio |
| SC2RBD_6m0j | SC2RBD | 40592-V08H-100 | Sino Biological |
| IL-7RA_3di3 | IL-7RA | 10975-H02H-100 | Sino Biological |
| PD-L1_5o45 | PD-L1 | 10084-H02H-100 | Sino Biological |
| TrkA_1www | TrkA | 11073-H08H-100 | Sino Biological |
| Insulin_4zxb | Insulin | 11038-HNAY-100 | Sino Biological |
| H1N1_5vli | H1N1 | 11684-V08H-100 | Sino Biological |
| VEGF-A_1bj1 | VEGF_A | 11066-HNAH-20 | Sino Biological |
| IL-17A_4hsa | IL-17A | 12047-HNAS-100 | Sino Biological |
| TNFa_1tnf | TNFa | 10602-H08H-50 | Sino Biological |

**Table S7.**
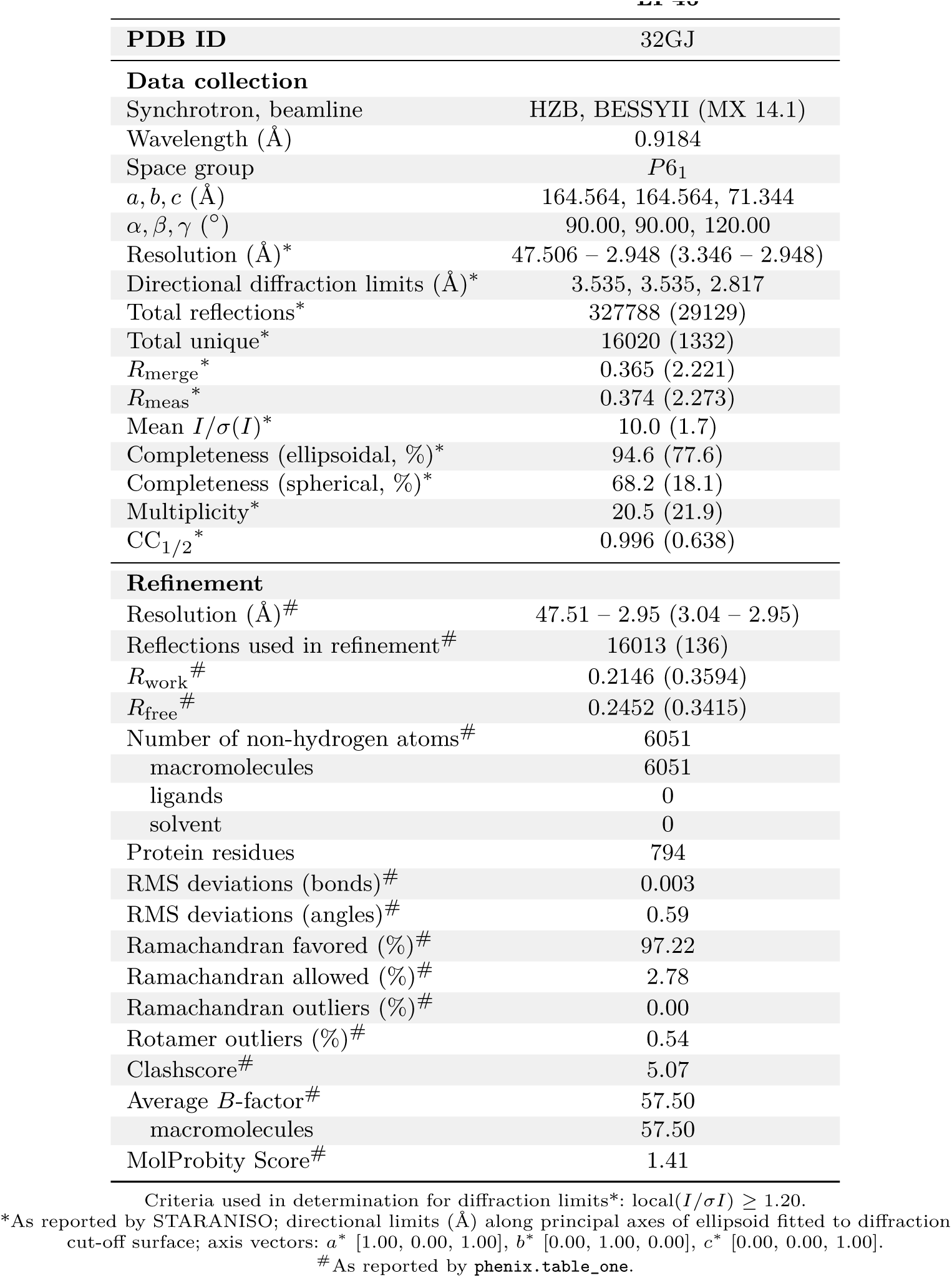
Data collection and refinement statistics for LP46.

**Table S8.**
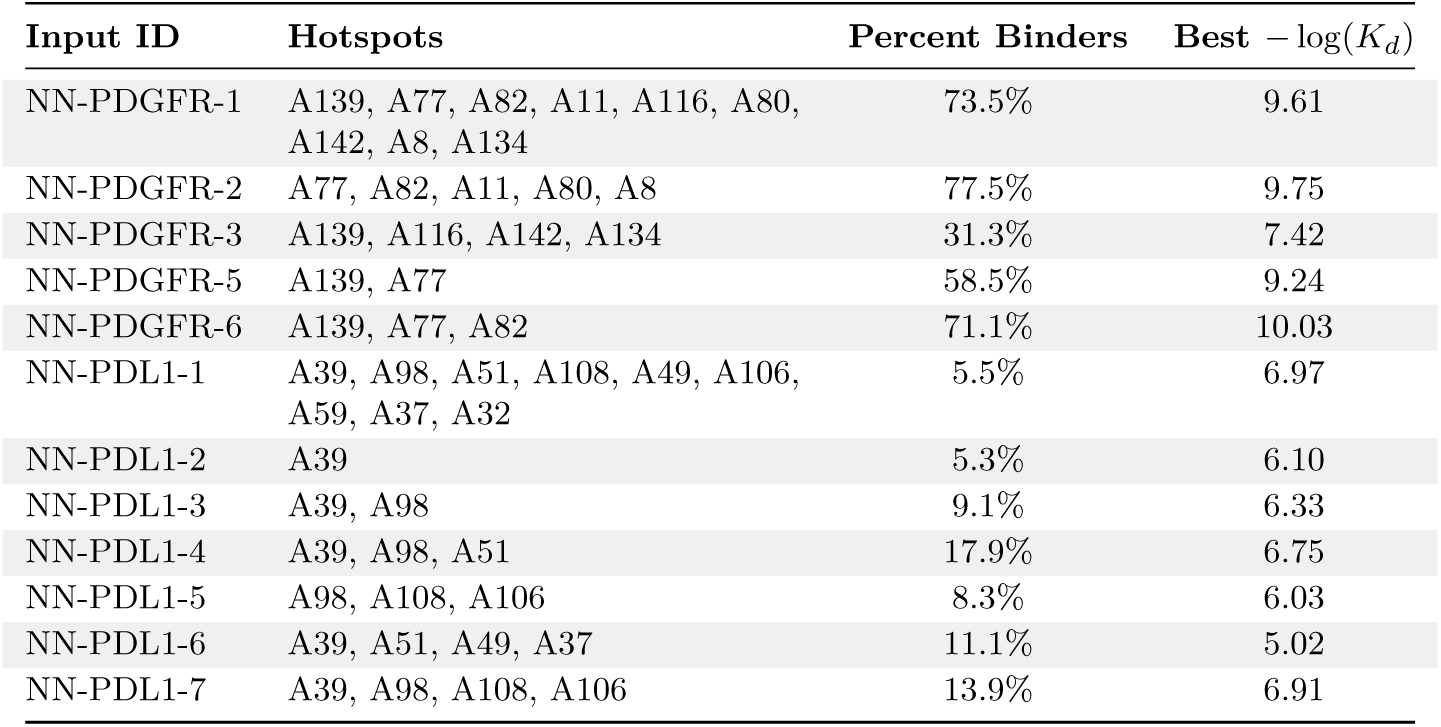
Evaluation of hotspot performance for binders to PDGFR and PDL1 targets.

**Table S9.** Expression and purification yields for the 200 ActRIIA binder designs. Soluble expression yields are reported after the first nickel-affinity purification step for both His-tagged (tagged) and TEV-cleaved (tagless) constructs. Of the 200 designs, 192 expressed solubly and 110 achieved yields exceeding 50 mg/L in the tagged format. Tag removal by TEV protease was unsuccessful for 22 constructs.

| Yield (after 1<sup>st</sup> nickel) | Tagged Constructs | Tagless |
| --- | --- | --- |
| No soluble expression | 9 | 9 |
| Tag can't be cleaved | – | 22 |
| <10 mg/L | 2 | 59 |
| 10–20 mg/L | 9 | 52 |
| 20–30 mg/L | 12 | 36 |
| 30–50 mg/L | 58 | 20 |
| >50 mg/L | 110 | 2 |
| **Total** | **200** | **200** |

